# Mapping the genomic landscape of multidrug resistance in *Plasmodium falciparum* and its impact on parasite fitness

**DOI:** 10.1101/2023.06.02.543338

**Authors:** Sachel Mok, Tomas Yeo, Davin Hong, Melanie J. Shears, Leila S. Ross, Kurt E. Ward, Satish K. Dhingra, Mariko Kanai, Jessica L. Bridgford, Abhai K. Tripathi, Godfree Mlambo, Anna Y. Burkhard, Kate J. Fairhurst, Eva Gil-Iturbe, Heekuk Park, Felix D. Rozenberg, Jonathan Kim, Filippo Mancia, Matthias Quick, Anne-Catrin Uhlemann, Photini Sinnis, David A. Fidock

## Abstract

Drug-resistant *Plasmodium falciparum* parasites have swept across Southeast Asia and now threaten Africa. By implementing a *P. falciparum* genetic cross using humanized mice, we report the identification of key determinants of resistance to artemisinin (ART) and piperaquine (PPQ) in the dominant Asian KEL1/PLA1 lineage. We mapped *k13* as the central mediator of ART resistance and identified secondary markers. Applying bulk segregant analysis, quantitative trait loci mapping and gene editing, our data reveal an epistatic interaction between mutant PfCRT and multicopy plasmepsins 2/3 in mediating high-grade PPQ resistance. Susceptibility and parasite fitness assays implicate PPQ as a driver of selection for KEL1/PLA1 parasites. Mutant PfCRT enhanced susceptibility to lumefantrine, the first-line partner drug in Africa, highlighting a potential benefit of opposing selective pressures with this drug and PPQ. We also identified that the ABCI3 transporter can operate in concert with PfCRT and plasmepsins 2/3 in mediating multigenic resistance to antimalarial agents.

## Introduction

Malaria caused an estimated 247 million cases and 619,000 deaths in 2021, mostly in sub-Saharan Africa (*1*). Efforts to reduce the impact of malaria have repeatedly been stymied by the emergence and spread of *Plasmodium falciparum* resistance to antimalarial drugs (*2, 3*). Resistance to artemisinin (ART) derivatives, the fast-acting backbone of ART-based combination therapies (ACTs), was first reported in western Cambodia in the late 2000s and has since spread across the Greater Mekong Subregion (GMS) (*4, 5*). In patients, resistance manifests as delayed parasite clearance following ART treatment (*6*). Resistance to piperaquine (PPQ), the long-lasting partner drug of the widely-used first-line treatment dihydroartemisinin (DHA)+PPQ, was also first detected in western Cambodia and resulted in rapid loss of treatment efficacy across the GMS, with failure rates as high as 87% in northern Thailand (*7, 8*).

Single point mutations in Kelch13 (K13) have been identified as the primary mediator of ART resistance (*9–12*). Mechanistically, K13 mutations are thought to reduce endocytosis of host hemoglobin, effectively lowering ART activation by hemoglobin-derived heme (*13–15*), thus attenuating drug-mediated redox perturbations, protein alkylation and proteotoxic stress (*16–18*). Worryingly, mutant K13 variants have now emerged independently in Rwanda and Uganda, with evidence of delayed parasite clearance as well as increased survival of DHA-treated ring-stage parasites (*19–21*). Gene editing studies have identified a substantial role for the genetic background, implicating a role for additional determinants (*12, 22*).

Investigations into PPQ resistance initially identified multicopy *plasmepsins 2* and *3* (*pm2/3*) as biomarkers (*23, 24*). These aspartic proteases contribute to hemoglobin proteolysis that releases globin (a source of amino acids) and toxic heme. Subsequent studies identified a critical role for novel point mutations in the chloroquine resistance determinant PfCRT (such as M343L or F145I), which evolved on the background of the chloroquine-resistant Southeast Asian Dd2 mutant haplotype (**Fig. 1** legend; (*25–27*)). These novel variants are thought to mediate PPQ resistance via a gain of drug efflux through the central cavity of this digestive vacuole membrane-spanning transporter (*28, 29*). Efflux removes PPQ from its site of action in the digestive vacuole, where it normally accumulates (by >1,000 fold) and acts to inhibit heme detoxification and hemozoin formation (*30*). PPQ-resistant parasites display an unusual dose-response profile, in which the IC_50_ (i.e., the concentration that yield half-maximal growth inhibition) shows minimal difference relative to PPQ-sensitive parasites, yet at high PPQ concentrations parasites show a biphasic survival curve (*31*). Defining the relative contributions of PfCRT and *pm2/3*, in terms of resistance and fitness costs, has until now been elusive.

**Fig 1.**
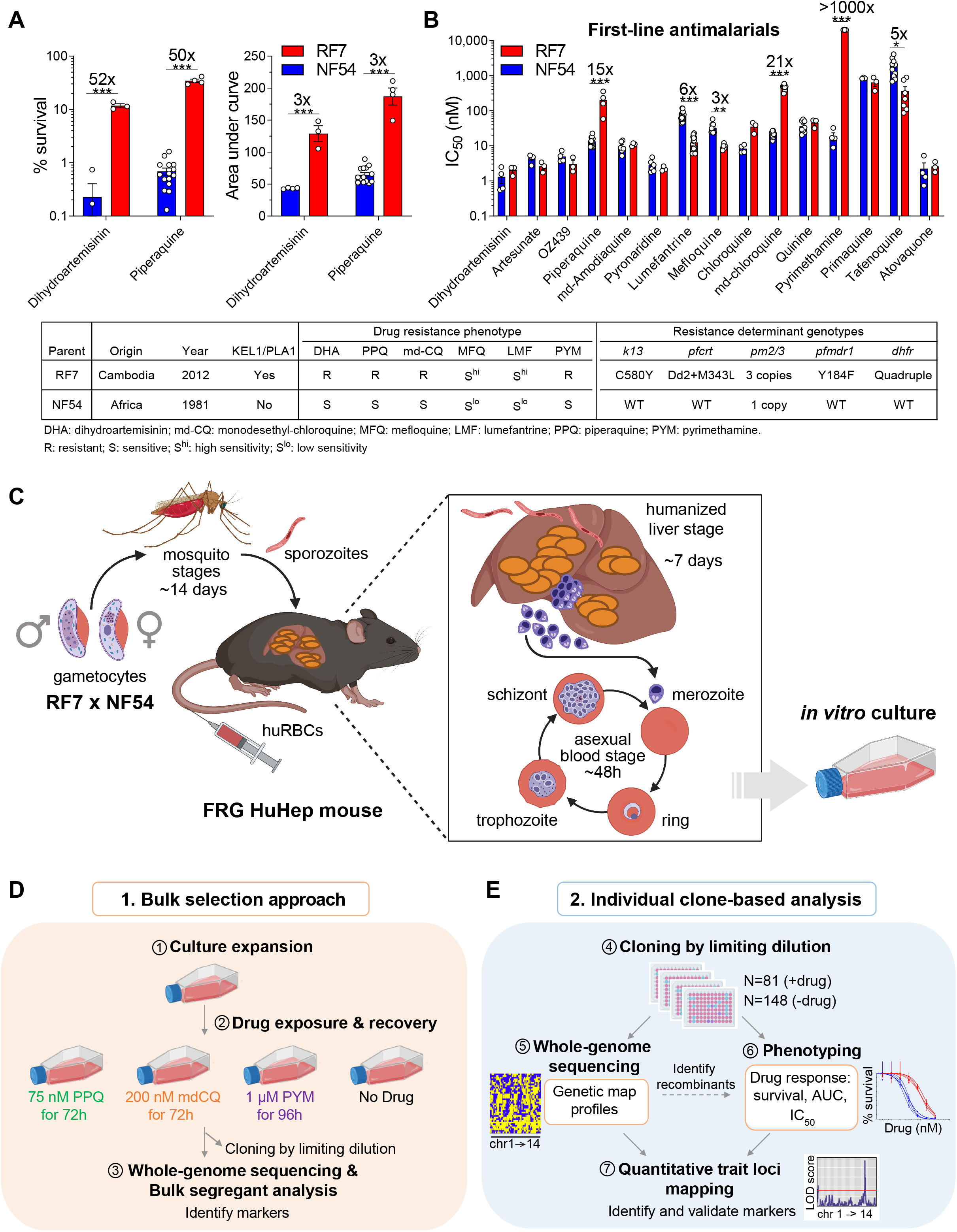
Antimalarial susceptibilities of the genetic cross parents RF7 × NF54 and the experimental workflow for bulk selection vs. individual clone-based linkage mapping. (**A**) Bar plots of % survival derived from the RSA or the PSA and corresponding AUC values for the genetic cross parents RF7 and NF54. In the RSA and PSA, early ring-stage parasites are exposed to the pharmacologically-relevant concentrations of 700 nM DHA for 4h or 200 nM PPQ for 72h, respectively, and survival measured as a percentage of mock-treated cultures. (B) IC_50_ values for first-line antimalarials of RF7 and NF54. Values represent the means ± SE across 3-17 independent experiments performed with technical duplicates. Statistical significance was determined by unpaired Student’s t-tests, with a Holm-Sidak post-test to correct for multiple comparisons. The IC_50_ fold shifts are indicated above the bars. \*\**P*<0.01, \*\*\**P*<0.001. Numbers listed above the statistics indicate the fold differences between parental IC_50_ values. The table summarizes parental phenotypic and genotypic characteristics. WT, wild-type. Mutant genotypes include: Dd2 *pfcrt* mutations: M74I, N75E, K76T, A220S, Q271E, N326S, I356T, and R371I; Quadruple *dhfr* mutations: N51I, C59R, S108N, and I164L. (C) Overview of the genetic cross pipeline performed in humanized FRG-NOD mice. (D) Bulk selection approach used for bulk segregant analysis, and (E) Individual clone-based linkage approach, used to identify genetic determinants of drug resistance. Images were created with BioRender.com.

To explore the genetic basis of *P. falciparum* resistance to DHA and PPQ, we conducted a genetic cross between a Cambodian multidrug-resistant parasite and a drug-sensitive African parasite. This cross leveraged the FRG-NOD humanized mouse model that permits pre-erythrocytic liver and asexual blood stage (ABS) replication of *P. falciparum* parasites (*32, 33*). Our data uncover secondary markers of ART resistance and elucidate the epistatic relationship between *pfcrt* and *pm2/3* in modulating PPQ resistance. We also identify the ABC transporter *abci3* as an additional member of this gene interaction network involved in differential susceptibility to other antimalarial compounds. Lastly, we identify PPQ-resistant *pfcrt* as a driver of enhanced sensitivity to lumefantrine (LMF) and mefloquine (MFQ), supporting their combined use in the field.

## Results

### Implementation of a RF7 × NF54 genetic cross to map determinants of drug resistance

To identify determinants of *P. falciparum* resistance to DHA and PPQ, we conducted a genetic cross between the PPQ- and DHA-resistant, contemporary Cambodian clinical isolate RF7, and the sensitive African parasite line NF54 (**Fig. 1**). RF7 was selected as representative of the KEL1/PLA1/Pailin (KelPP) lineage that has dominated the GMS (*5, 10*). We first characterized RF7 and NF54 susceptibility profiles to DHA, PPQ and a panel of other antimalarials (**Fig. 1A**). Using the ring-stage survival assay (RSA), we observed a high percentage of surviving DHA-treated parasites (mean±SEM, 11.8±1.0%), whereas NF54 showed marginal survival (0.2±0.2%). Likewise, using the PPQ survival assay (PSA), ring-stage RF7 parasites gave a high survival rate (34.5±2.6%), contrasting with minimal survival (0.7±0.1%) in NF54 (**Fig. 1A** and **Table S1**). RF7 parasites showed elevated Area Under the Curve (AUC) values for both drugs (**Fig. 1A**). RF7 was also highly resistant to monodesethyl chloroquine (mdCQ; the active metabolite of CQ) and pyrimethamine (PYM). In contrast, RF7 exhibited 3- to 6-fold increased sensitivity to the ACT partner drugs MFQ and LMF compared to NF54 (**Fig. 1B** and **Table S1**). There were no significant differences in IC_50_ values for other antimalarials tested, including amodiaquine, pyronaridine, quinine and atovaquone.

To achieve a genetic cross, RF7 and NF54 gametocytes were fed as 1:1 pools to female *Anopheles stephensi* mosquitoes to initiate sexual stage recombination. We then infected four human liver-chimeric FRG-NOD mice with sporozoites, either via intravenous inoculation of manually dissected sporozoites or via mosquito bites (**Fig. 1C** and **Table S2**). Haploid ABS parasites were recovered on day 7.5 post-infection and subsequently cultured *in vitro* (**Fig. 1C**).

### Bulk segregant analyses identify major determinants of *P. falciparum* resistance to piperaquine, chloroquine and pyrimethamine

Whole-genome sequencing (WGS) of the RF7 and NF54 parents identified 18,489 (“18k”) single nucleotide polymorphisms (SNPs) between their core genomes (**Tables S3** and **S4**). These SNPs were located in 2,439 *P. falciparum* coding genes, corresponding to nearly half of the core genome of ∼5,100 genes, with an average of ∼1 kb distance between SNPs. These SNPs were distributed quite evenly across coding and non-coding regions (**fig. S1, A** to **C**). This extensive diversity between these geographically distinct parents enabled high-resolution genetic mapping of drug susceptibility profiles in the cross progeny.

We applied a bulk selection approach to enrich for recombinant ABS progeny resistant to PPQ, mdCQ or PYM (**Fig. 1D**). Surviving parasites from these drug-pressured bulk cultures were then subjected to WGS, using this 18k SNP set to measure parental allele frequencies across the 14 chromosomes (**Fig. 1D**). Using bulk segregant analysis, we then compared the parental frequencies at each SNP position between drug-treated or untreated cultures (**Fig. 2** and **fig. S2**). Comparing the PYM-treated bulk against the mdCQ-treated sample or the untreated bulk revealed a significant quantitative trait locus (QTL) on chr4 (**Fig. 2, Table S5** and **fig. S2A**). This locus, spanning 141 kb, contained dihydrofolate reductase (*dhfr*) that in RF7 parasites encoded a quadruple mutant PYM-resistant allele. By comparison, mdCQ selected for a dominant 856 kb QTL on chr7 that contains the mutant Dd2+M343L allele of the *P. falciparum* chloroquine resistance transporter *(pfcrt)*, as well as two smaller peaks on chr6 and 12 that containing the putative amino acid transporter *aat1* and the v-type pyrophosphatase *vp2*. mdCQ and PPQ showed similar allelic frequency profiles, consistent with their similar modes of action (*30*). Nonetheless, PPQ selected for a 117 kb chr14 segment containing the tandem *pm2/3* amplicons, whereas this drug selected against a 484 kb segment on chr13 harboring mutant E415G exonuclease-I, previously proposed to be a molecular marker of PPQ resistance (*24*) (**Fig. 2** and **fig. S2**).

**Fig. 2.**
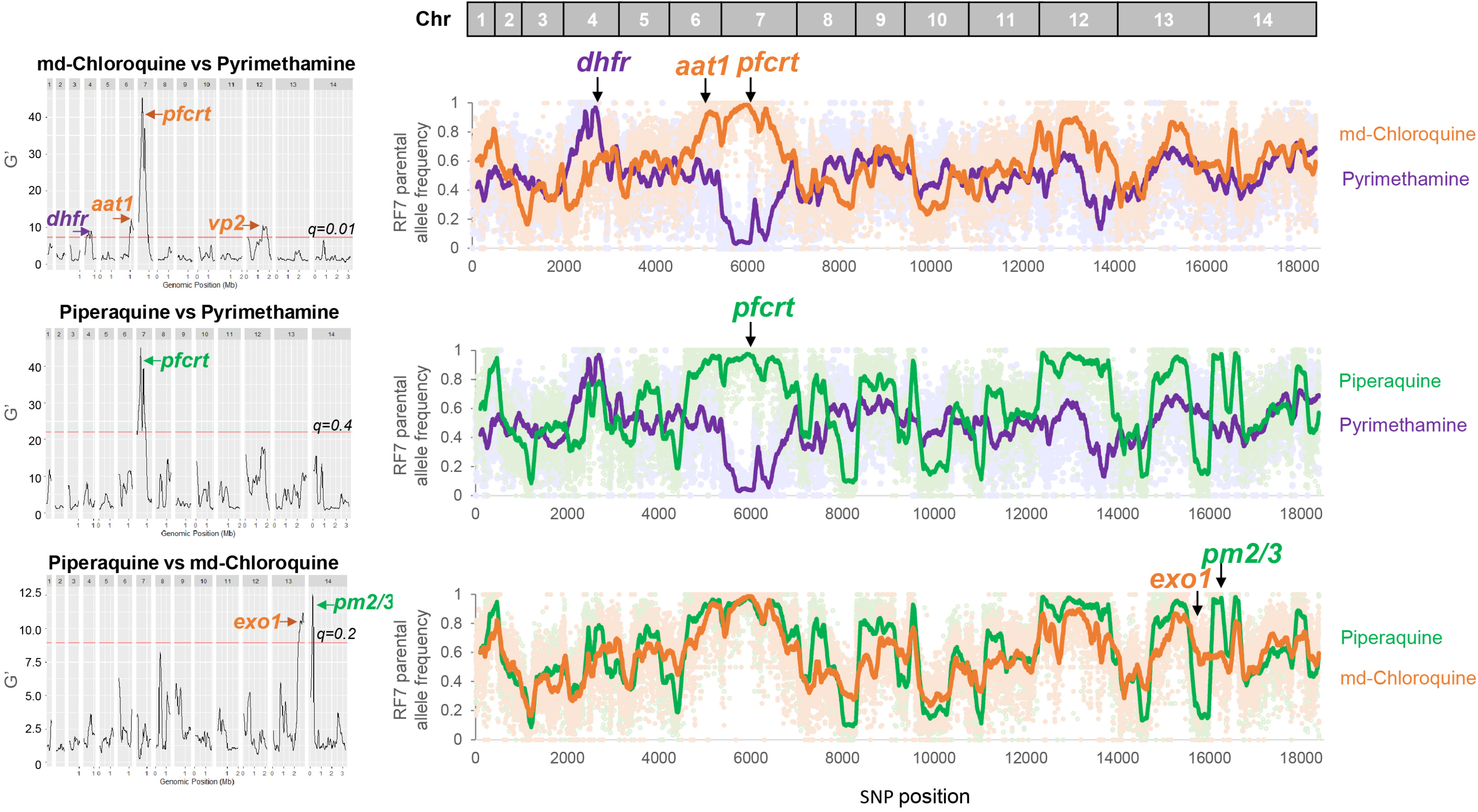
Bulk segregant analyses of progeny pools pressured with PPQ, mdCQ or PYM. Shown are the significant genetic loci enriched by each drug in individual drug-drug pairwise comparisons, and their corresponding RF7 parental allele frequencies for the set of “18k” SNPs that differ between the two parents. Regions with a False Discovery Rate (FDR) *q*-value < 0.01 (mdCQ vs. PYM), 0.4 (PPQ vs. PYM), or 0.2 (PPQ vs. mdCQ), were considered statistically significant QTLs.

Although bulk segregant analysis could identify loci containing the major resistance mediators for these drugs, this approach can be constrained by the fitness cost of certain genotypes that show reduced expansion in bulk cultures. To refine the identification of resistance markers and map secondary determinants, as well as understand epistatic interactions between genes, we therefore performed QTL analysis with individual clones (**Fig. 1E**).

### Enrichment of genetically & phenotypically diverse progeny using drug pulses

#### Identification of genetically distinct recombinant progeny

We cloned 148 individual progeny by limiting dilution of untreated cultures, and identified 66 recombinant progeny using WGS analysis of the 18k SNP set. The remaining 82 progeny proved to be genetically identical to NF54, an indication of a high selfing rate for this parent in our cross. The 66 recombinant clones segregated into 12 unique haplotypes (**Fig. 3A**). An additional 81 recombinant clones were obtained from drug-pressured bulk cultures, yielding an additional 22 unique haplotypes (**Fig. 3, A** and **B**). Subsequent analysis focused on these 34 genetically distinct haplotype groups (**Fig. 3B**), in addition to HapA and HapN, corresponding to NF54 and RF7, respectively. Surprisingly, we observed minimal overlap in the sets of recombinant progeny selected using PPQ and mdCQ pressure. Also, no common haplotypes were obtained using PYM pressure compared with either of the other two drugs (**Fig. 3B**). PPQ and mdCQ, respectively, generated five and six unique recombinant haplotypes, and one additional shared haplotype. This finding suggests that resistance to each drug requires distinct combinations of genetic determinants.

**Fig. 3.**
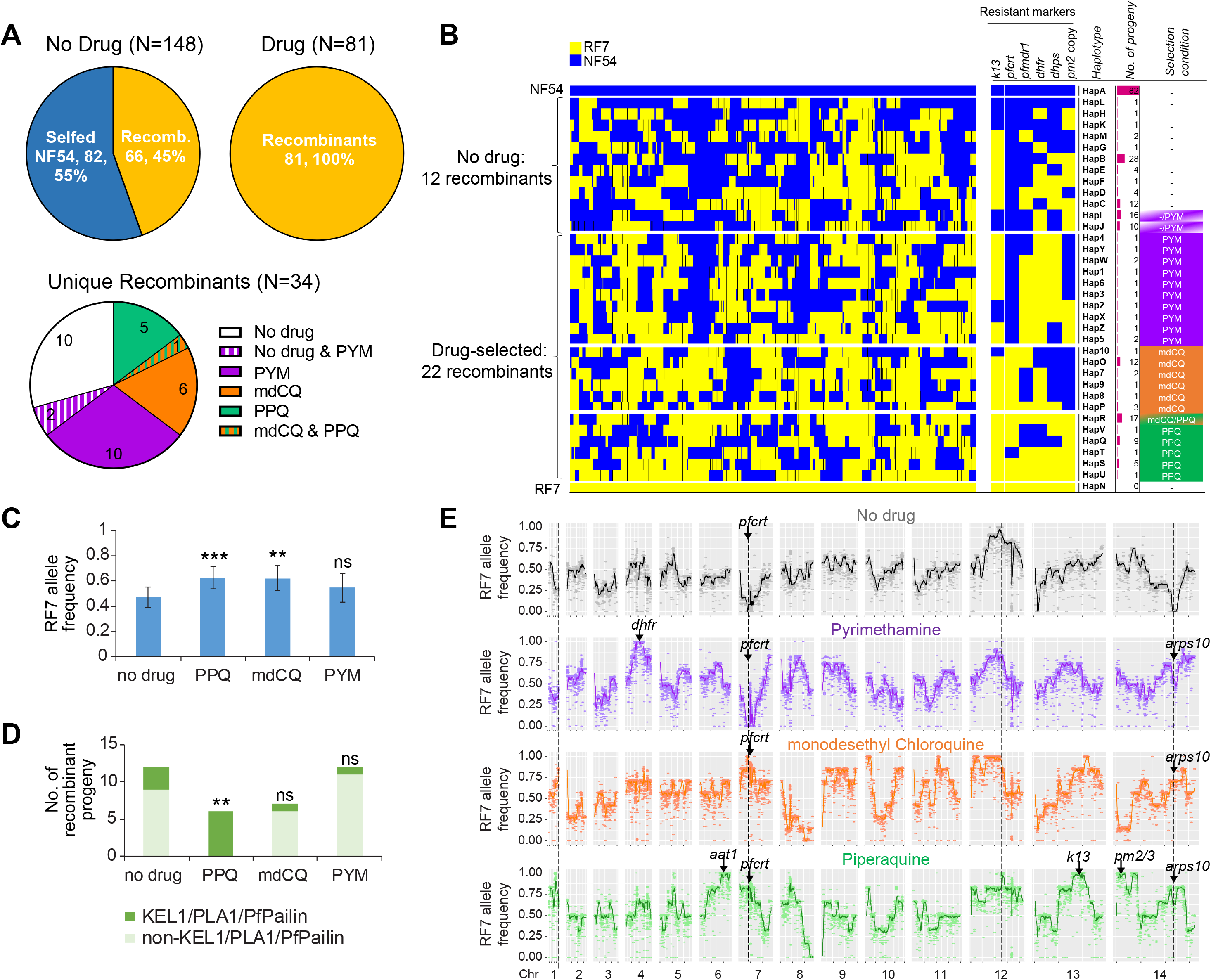
Genetic analysis of recombinant progeny and enrichment of diverse progeny using drug pulses. (**A**) Number of selfed progeny vs. genetic recombinants obtained from cloning in the absence of drug or following drug exposure. Distribution of the 34 unique recombinants selected by the various ± drug conditions of PPQ, monodesethyl chloroquine (CQ) or pyrimethamine (PYM). (**B**) Allelic map for the 34 recombinant progeny grouped by drug-treatment condition. Shown are the parental alleles inherited by each progeny for 14,476 SNPs (after excluding SNPs missing in any of the 36 haplotypes), the genotypes of known resistant markers (*k13, pfcrt, pfmdr1, dhfr, dhps* and *pm2* copy) and the number of derived progeny clones per haplotype sorted by selection condition. (**C**) Mean RF7 allele frequency inherited by the 34 recombinant progeny clones for each drug-treatment group. (**D**) Count of KEL1/PLA1/PfPailin haplotypes in progeny clones that were selected by each drug condition. Statistical significance was tested for the drug-selected progeny clones against the “no drug” group by Fisher’s exact test. \*\**P*<0.01; ns, not significant. (**E**) Frequency of RF7 parental allele in progeny clones derived post-treatment with PPQ (N=6), mdCQ (N=7) or PYM (N=12) or in the absence of drug (N=12), showing the skews in allele frequency at specific loci in the genome, as indicated by dashed lines.

#### Analysis of recombination rates, inheritance patterns and genetic relatedness among progeny

Our analysis of the 34 unique recombinants revealed a recombination rate of 15.1 kb/cM, and a positive correlation (R^2^=0.47) between number of crossover events and physical lengths (**Table S6** and **fig. S1, D** and **E**), similar to previous crosses (12.1 – 14.3 kb/cM; (*33–35*). Most clones shared ∼50% similarity in parental allelic composition across the genome, as measured by identity-by-descent analysis (**fig. S1F**). Nonetheless, using multidimensional scaling of pairwise genetic distances, we observed that the recombinant progeny clustered based on their selection conditions (**fig. S1G**). Progeny derived with PPQ or mdCQ pressure clustered more closely and were distinct from the progeny pressured with PYM or derived without drug. This observation agrees with our bulk segregant results and drug chemical relatedness. There was also significantly higher inheritance of RF7 SNPs in the recombinants derived following PPQ or mdCQ exposure compared to those obtained in the absence of any drug pressure (**Fig. 3, B** and **C** and **fig. S1H**), suggesting linkage disequilibrium and the possibility of multiple determinants.

#### PPQ and mdCQ show differential selective pressure for the KEL1/PLA1/PfPailin haplotype

Interestingly, we observed that all progeny selected by PPQ were of the KelPP haplotype (*k13* C580Y, *k13*-flanking region, and multicopy *pm2/3*), identical to RF7 (**Fig. 3D** and **fig. S3A** and **Table S7**). However, only one of the seven mdCQ-selected progeny inherited this haplotype. While mdCQ treatment enriched for progeny clones harboring mutant *pfcrt*, this drug selected against a segment on chr14 encompassing *pm2/3* (**Fig. 3B**). This observation suggests that amplified *pm2/3* may have a fitness cost in the presence of RF7 mutant *pfcrt*. Likewise, the majority of recombinants obtained under PYM or drug-free conditions did not harbor the KelPP lineage (**Fig. 3D**). This haplotype’s significant association with PPQ-selected progeny (*P*=0.009, Fisher’s exact test) but not with the mdCQ or PYM-selected progeny suggests that PPQ favors this particular lineage.

#### Genomic segregation distortions map to mutant pfcrt and a chr14 loci

To identify loci under segregation distortion in the progeny, we aggregated the distinct sets of clones recovered without or with drug selection *in silico* and examined their allele frequencies. In the drug-negative clones, there was extreme inheritance bias towards NF54 wild-type (WT) SNPs at multiple loci on chromosomes 1, 7, 13, and 14. These skews were reversed by PPQ, mdCQ or PYM treatments that enriched for RF7 SNPs in these segments (**Fig. 3E**). This reversal was most prominent for *pfcrt* found on chr7 in mdCQ- or PPQ-pressured parasites, implying a strong fitness cost for mutant *pfcrt* parasites in the absence of selective pressure. Using the “pooled” drug-selected progeny clones, we also identified an additional 251 kb segment on chr14 that was enriched by all three drugs (**Fig. 3E**). Besides *pfcrt*, the PPQ-selected clones showed 100% inheritance of RF7 SNPs for chr14 (377 kb) and chr13 (516 kb) segments containing *pm2/3* and *k13*, respectively (**Fig. 3E**). This observation supports our finding that PPQ positively selected for the KelPP lineage. In addition, PPQ but not mdCQ or PYM, selected for a 416 kb chr6 segment encompassing *aat1*, suggesting a possible involvement of this marker with PPQ resistance.

#### Drug exposure enriches for progeny with diverse phenotypic responses

To characterize the DHA and PPQ phenotypic responses of the 34 recombinant haplotypes, we profiled a panel of representative progeny and their RF7 x NF54 parents (**Fig. 4** and **Table S8**). Interestingly, we observed a wider range of phenotypic responses and higher survival and AUC values for the clones derived post-drug selection compared to the non-selected group (**Fig. 4, C** and **D**). Several of these drug-selected progeny clones had even higher RSA survival and AUC values compared to parental RF7. This contrasted with the PPQ response where none of the progeny were more resistant than RF7 in their PSA survival and AUC levels. All non-selected clones remained as equally susceptible as NF54. This suggests a large fitness cost among the PPQ-resistant progeny and reinforces the importance of applying drug pressures to diversify genetic cross progeny.

**Fig. 4.**
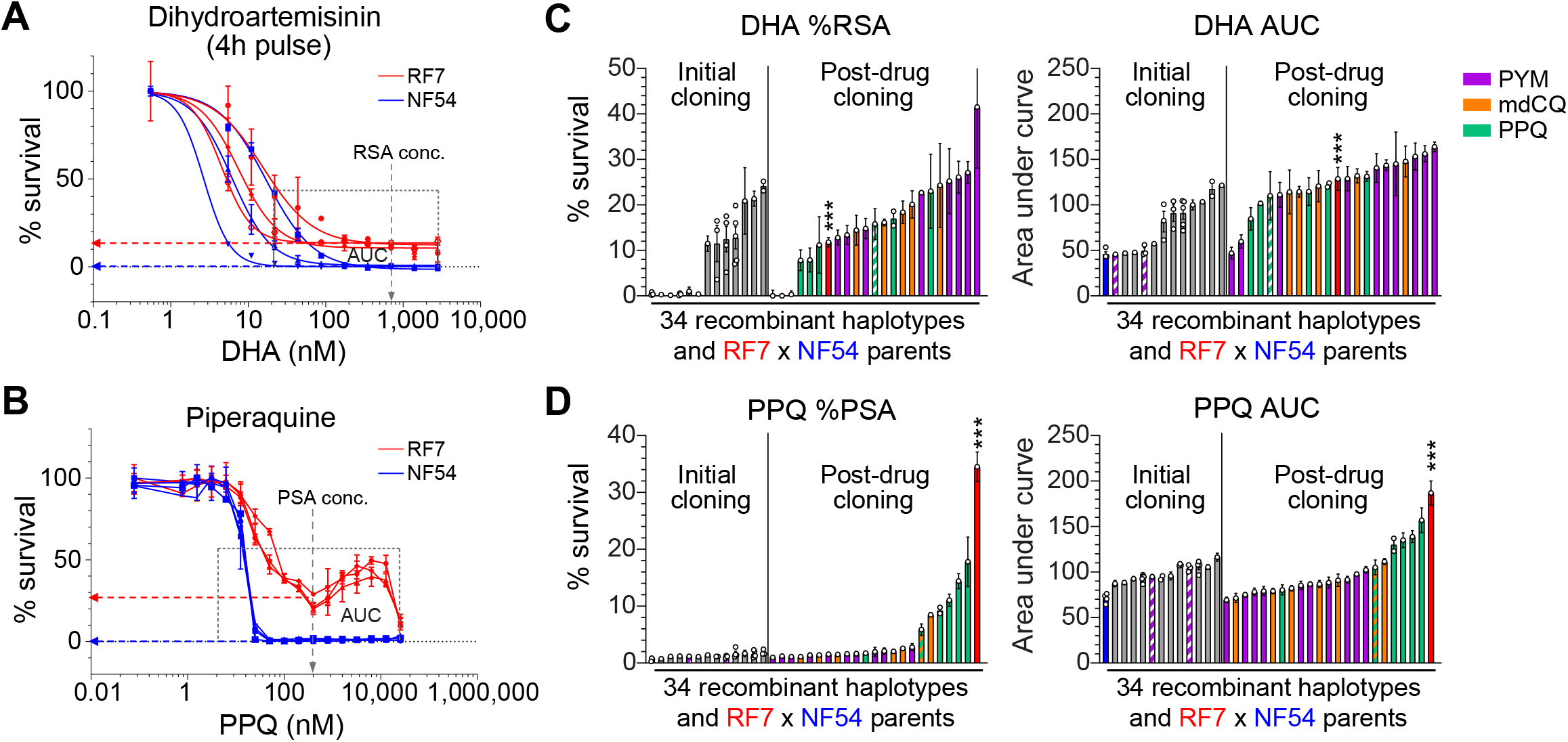
Phenotypic response of parents and progeny to DHA and PPQ. (**A** and **B**) Dose-response curves for RF7 and NF54 parents across a range of DHA (**A**) and PPQ (**B**) concentrations (N,n=3,2). The % survival at the RSA and PSA concentrations of 700 nM DHA and 200 nM PPQ for 4h or 72h, respectively, were used to determine DHA and PPQ resistance levels, whereas AUC values were measured as total survival across a range of concentrations (22 nM to 2.8 µM for DHA and 1.6 nM to 25.6 µM for PPQ). (**C** and **D**) DHA and PPQ response measured by % RSA and DHA AUC values (**C**), and % PSA and PPQ AUC values (**D**), respectively, in the 46 recombinant and three NF54-selfed progeny that were derived from cloning ± drug exposure. Each bar represents the mean % survival or AUC ± SE for a recombinant haplotype and ordered by the parasites’ resistance levels in ascending order independently for each drug index. For certain haplotype groups in which identical clones were obtained, multiple points depict more than one sibling progeny being phenotyped. Significance between the genetic cross parents’ responses were tested by unpaired Student’s t-tests (N,n=3-4,2). \*\*\**P*<0.001. RSA, ring-stage survival assay. PSA, piperaquine survival assay. AUC, area under the curve.

### DHA resistance associates with *k13* and other secondary determinants

#### QTL mapping of DHA resistance reveals k13 as the primary determinant

We next performed linkage analysis to uncover QTLs associated with DHA resistance, using the phenotype-genotype data from the progeny clones (**Tables S4** and **S8**). There was a good correlation (R=0.85) between RSA and AUC values (**fig. S4**). QTL mapping using either sets of values revealed a dominant peak on chr13. This 183 kb segment contained 109 SNPs including 47 non-synonymous mutations within 20 genes, and included *k13,* the primary ART resistance mediator (**Fig. 5, A** to **D**). We also observed co-inheritance of this segment flanking the region -133 kb to +47 kb of the *k13* gene in all mutant progeny (**fig. S3**), suggesting the possible presence of additional loci that have an impact on ART resistance or compensate for altered mutant K13 function. Grouping the recombinant progeny by their *k13* genotypes (C580Y vs. WT) showed a clear segregation in DHA response with mutant progeny showing higher RSA survival and AUC levels (**Fig. 5E**). Reverting C580Y to WT in RF7 parasites, using CRISPR/Cas9-based gene editing, resulted in a significant reduction in RSA and AUC levels (**Fig. 5F**).

**Fig. 5.**
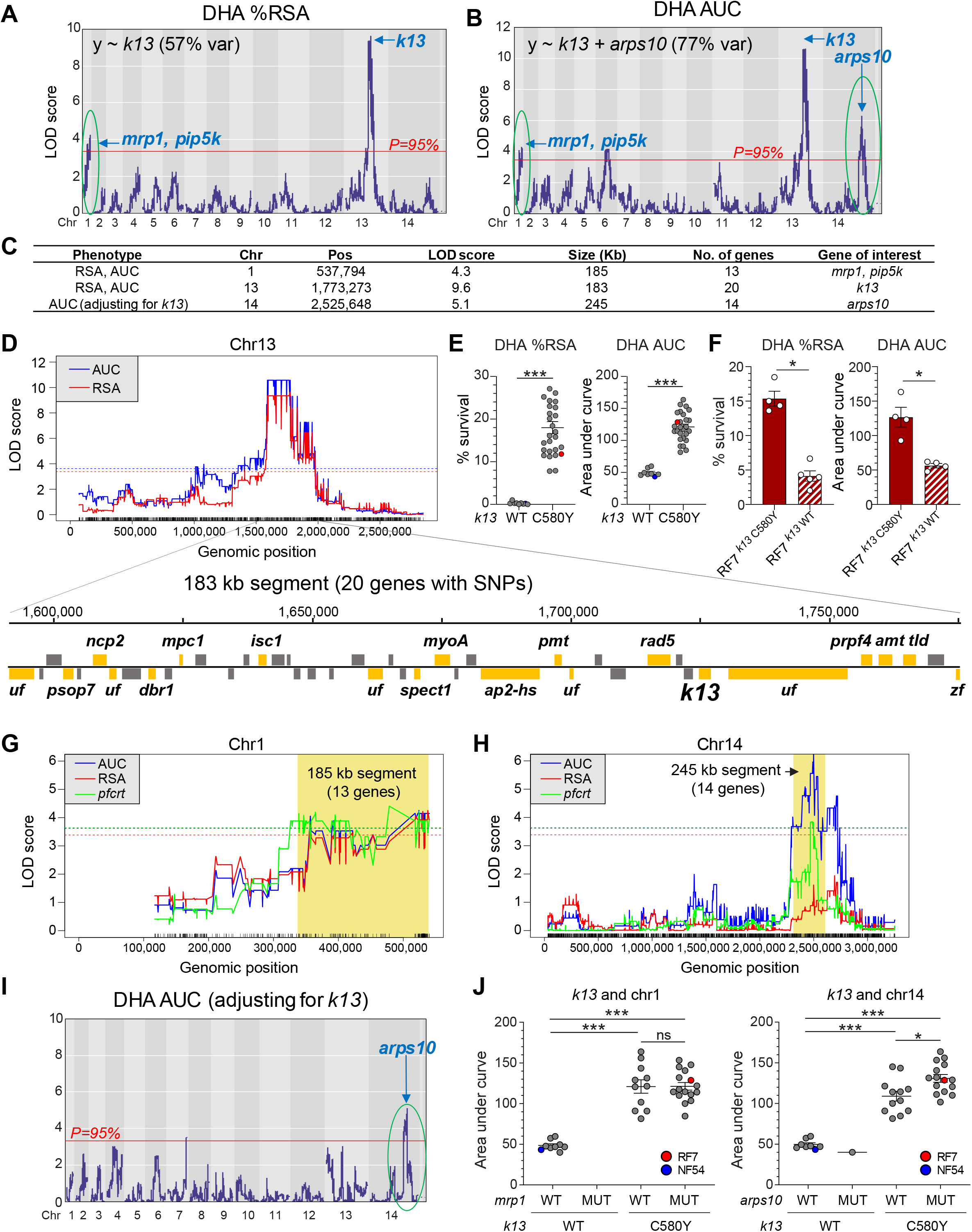
QTL mapping of DHA response in progeny identifies *k13* as the primary DHA resistance locus. (**A** and **B**) LOD plots for % RSA (**A**) and AUC levels (**B**) showing the significant QTLs above the 95% probability threshold (red line). (**C**) List of QTL segments for % RSA and AUC levels. (**D**) Genes in QTL segment on chr13, with those having non-synonymous mutations between RF7 and NF54 colored in orange (n=20), or grey where mutations were absent in RF7 and NF54. (**E**) The % RSA and AUC levels in recombinant progeny segregated by *k13* parental allele, C580Y (RF7) and WT (NF54). Significance was tested using Mann-Whitney U. \*\*\**P*<0.001. (**F**) DHA response in *k13*-edited isogenic RF7 clones showed that *k13* WT allele reduces % RSA and AUC levels. Bars represent the means ± SE for four independent experiments performed in technical replicates. Significance was tested using Mann-Whitney U. \**P*<0.05. (**G** and **H**) Linkage of chr1 (**G**) and chr14 (**H**) segments with *pfcrt* from QTL analysis using the *pfcrt* genotype as an outcome. Shown are the significant QTLs above the 95% probability threshold for each analysis. (**I**) LOD plot for AUC levels after adjusting for *k13* as a covariate suggests independent inheritance of chr14 segment with *k13* and co-inheritance of chr1 segment with *k13*. (**J**) Scatter plot of AUC levels in progeny segregated by *k13*, *mrp1* (chr1) and *arps1* (chr14) genotypes showing that the RF7 chr14 segment increases DHA AUC levels. Significance between groups were tested using Mann-Whitney U. \**P*<0.05, \*\*\**P*<0.001.

#### QTL mapping uncovers potential secondary determinants on chr1 and chr14

Besides the chr13 segment, we observed two additional smaller QTLs on chr1 and chr14 that were significantly associated with DHA resistance (**Fig. 5, A, B, G, H,** and **Table S9**). We then examined whether these associations were additive or epistatic in modulating DHA response. Applying *k13* as a covariate resulted in the loss of the chr1 peak for both RSA and AUC, suggesting that the chr1 segment was likely co-inherited with *k13* (**Fig. 5I** and **fig. S5A**). This was evident by the lack of any difference in RSA levels when stratified by the *k13* and chr1 genotypes (**Fig. 5J**). In contrast, the chr14 segment showed a significant QTL peak for AUC after correcting for *k13* genotypes or the subset of *k13* mutants (**Fig. 5I**, **fig. S5B** and **Table S9**). Mutant *k13* progeny exhibited higher mean AUC levels when harboring RF7 alleles for chr14, suggesting that this segment might enhance parasite survival at higher doses of DHA (**Fig. 5J** and **Table S9**).

Within the chr1 185 kb segment, we observed non-synonymous mutations in 13 genes, including the drug-resistant candidates *mrp1* and *pip5k* (**Table S9**). The chr14 245 kb segment, significant in only the AUC outcome, comprised 14 genes including *arps10*. Examination of sequence diversity in these three genes, across the set of ∼2,500 sequenced *P. falciparum* genomes (*36*), suggested geographic differences with the RF7 parental mutations being prevalent in clinical isolates from Cambodia, Mali and Ghana (**fig. S5C**). Interestingly, these chr1 and chr14 segments also showed linkage disequilibrium with *pfcrt* (**Fig. 5, G** and **H,** and **fig. S5, D** and **E**).

### PPQ resistance is conferred by mutant Dd2+M343L *pfcrt* and multicopy *pm2/3*

To identify the markers of PPQ resistance, we performed QTL mapping using the recombinant progeny values obtained from the PSA and AUC data (**Table S8**). Both data sets identified a major QTL on chr7, which mapped to *pfcrt* (**Fig. 6, A** to **C** and **Table S10**). Our AUC analyses also identified an additional locus on chr14 harboring the conserved *pm2/3* amplicon (**Fig. 6, B** and **C, fig. S6A** and **Table S10**). Controlling for *pfcrt* as a covariate revealed no other peaks in the PSA analysis, suggesting that the *pfcrt* RF7 allele is the main determinant of survival at 200 nM PPQ (**fig. S6B**). In contrast, controlling for *pfcrt* improved the logarithm of the odds (LOD) score for the *pm2/3* segment for AUC. Reciprocally, controlling for *pm2/3* revealed a higher LOD score for the *pfcrt* segment (**fig. S6B**). Our results suggest a link between *pfcrt, pm2/3* and PPQ AUC.

**Fig. 6.**
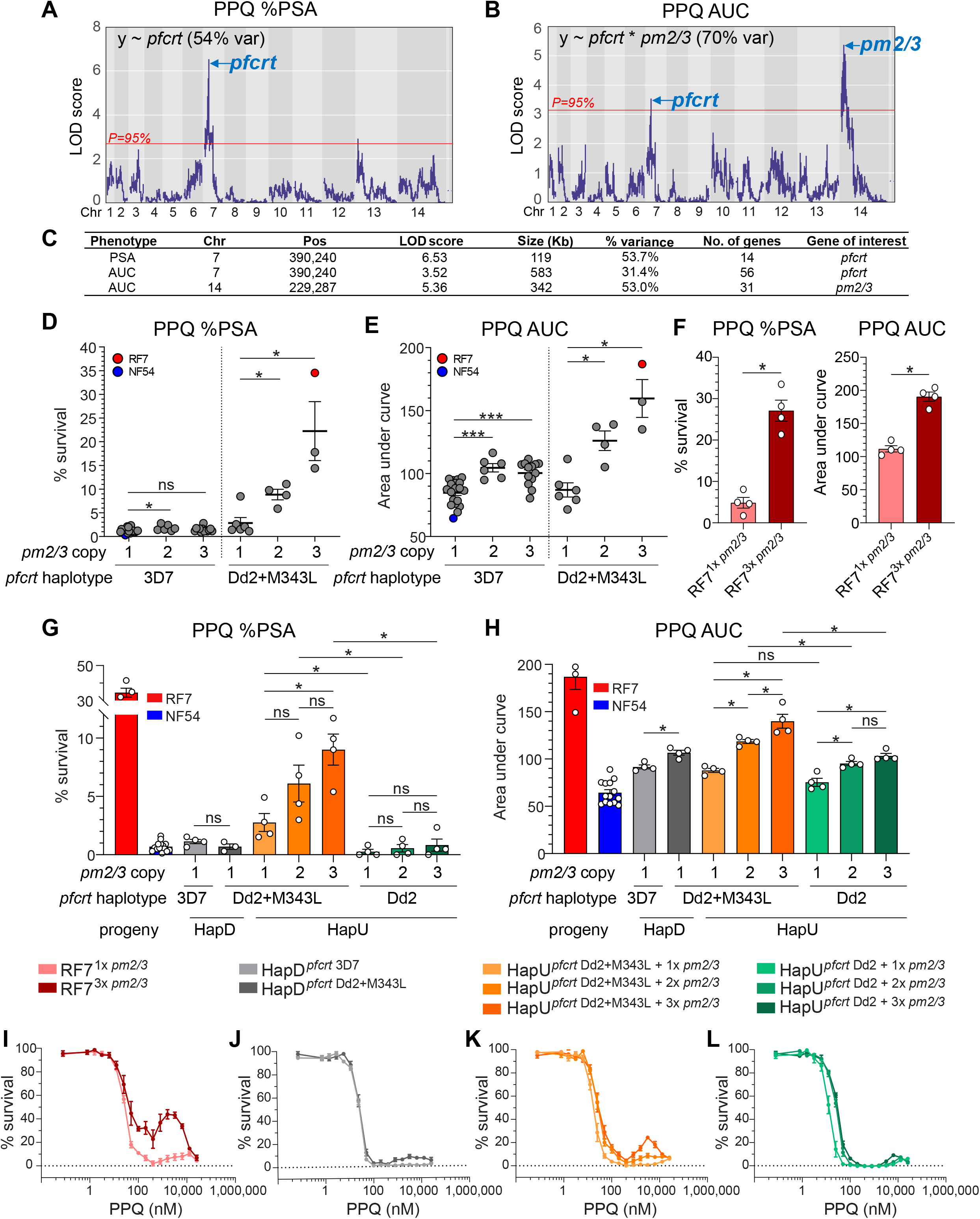
QTL mapping of PPQ resistance in progeny and response in *pfcrt*-edited progeny reveals an association of PPQ-R with mutant Dd2+M343L *pfcrt* and multicopy *pm2/3*. (**A** and **B**) LOD plots for % PSA (**A**) and AUC levels (**B**) showing QTLs detected above the 95% probability threshold (red line). (**C**) List of QTL segments for % PSA and AUC levels. (**D** and **E**) Scatter plot of the % PSA (**D**) or AUC levels (**E**) for 34 independent recombinant progeny and parents segregated by *pm2/3* copy and *pfcrt* genotypes. Significant differences in PPQ response between the recombinant groups harboring different *pm2/3* copies were tested by Mann-Whitney U. \**P*<0.05, \*\*\**P*<0.001. (**F**) PPQ response in isogenic RF7 clones containing 1 vs. 3 copies of *pm2/3* in the Dd2+M343L *pfcrt* background showing that single copy *pm2/3* reduces % PSA and AUC levels. Bars represent the means ± SE for four independent experiments performed in technical replicates. Significance was tested using Mann-Whitney U. \**P*<0.05. (**G**) PPQ % PSA and (**H**) AUC levels in *pfcrt*-edited HapD progeny (3D7 vs. Dd2+M343L) having single *pm2/3* copy and in edited HapU progeny (Dd2 vs. Dd2+M343L) having either 1, 2 or 3 copies of *pm2/3*. Bars represent the means ± SE for four independent experiments performed in technical replicates. Significance between the isogenic edited progeny lines was tested using Mann-Whitney U. \**P*<0.05; ns, non-significant. The colored key for parasites lines applies to panels **G** to **L**. (**I** to **L**) Dose response curve in RF7 clones (**I**), HapD (**J**), unedited HapU (**K**) and edited HapU (**L**) progeny showing that multicopy *pm2/3* and mutant *pfcrt* are necessary for the PPQ biphasic response. In parallel studies, serum was noted to lower % PSA (**fig. S8, B** and **C**). Here, each line depicts the mean % parasite survival ± SE across 3-4 independent experiments performed in technical duplicates.

Examining the PPQ response in WT *pfcrt* progeny revealed a small but significant association between *pm2/3* amplifications and higher AUC levels (**Fig. 6, D** and **E**). Mutant *pfcrt* progeny harboring multicopy *pm2/3* showed a larger increase in both PSA and AUC levels, coinciding with a biphasic response at high PPQ concentrations (**fig. S7**). These findings suggest an epistatic interaction between *pm2/3 and pfcrt* in modulating high-grade PPQ resistance. To further examine the relative contributions of *pm2/3* and *pfcrt*, we took advantage of a loss of *pm2/3* copy number in the RF7 parental line over ∼3 months of continuous *in vitro* culture, and used limiting dilution to obtain clones with 3 or 1 copies of *pm2/3*. The multicopy *pm2/3* clone exhibited significantly higher PSA and AUC levels and displayed a biphasic PPQ dose-response curve (**Fig. 6, F** and **I** and **fig. S8**). These data confirm that *pm2/3* amplifications can augment PPQ resistance in the presence of mutant *pfcrt*.

We validated these markers using gene-editing of progeny with differing *pm2/3* copy numbers. Introducing the mutant *pfcrt* allele into HapD WT *pfcrt* parasites having single copy *pm2/3* did not affect PSA levels and led to a minor increase in AUC levels, suggesting that the Dd2+M343L *pfcrt* allele did not suffice to confer PPQ resistance in this progeny (**Fig. 6, G, H** and **J**). We also tested HapU mutant *pfcrt* clones that displayed 1, 2 or 3 copies of *pm2/3*, showing higher PSA and AUC levels with increasing *pm2/3* copy numbers (**Fig. 6, G, H** and **K**). Removing the *pfcrt* M343L mutation from these parasites yielded a significant reduction in PSA and AUC levels irrespective of *pm2/3* copy number (**Fig. 6, G**, **H** and **L**). Our data also provided evidence that the amplification of *pm2/3* might constitute a first step of resistance as shown by enhanced parasite survival at lower PPQ concentrations (up to 100 nM). In the presence of multicopy *pm2/3*, the *pfcrt* M343L mutation further boosted parasite survival upon exposure to higher PPQ concentrations and generated a biphasic response (**Fig. 6, H** and **K**). Our results reveal an epistatic genetic interaction between these two markers, *pfcrt* and *pm2/3*, in modulating levels of PPQ resistance.

### Multicopy *pm2/3* and mutant *k13* associate with reduced fitness in the RF7 KEL1/PLA1 parasite

We next investigated the impact the *k13* and *pm2/3* resistance determinants on parasite fitness in the absence of drug pressure. To study this, we performed long-term pairwise competitive assays between *k13* C580Y mutant vs. WT clones that were edited in isogenic RF7 clones harboring 1 or 3 copies of *pm2/3* (**Fig. 7A**). The *k13* C580Y mutation exhibited a small fitness cost in the presence of multicopy *pm2/3*, whereas this k13 mutation was fitness neutral on the single copy *pm2/3* background (**Fig. 7, B** and **C**). A greater fitness cost was observed with multicopy *pm2/3* compared to single copy *pm2/3*, in *k13* C580Y RF7 parasites (**Fig. 7, B** and **C**). These data are consistent with our observation of spontaneous deamplification of *pm2/3* in RF7 and HapU parasites in the absence of selection pressure that associated with a decrease in PPQ resistance (**fig. S8**).

**Fig. 7.**
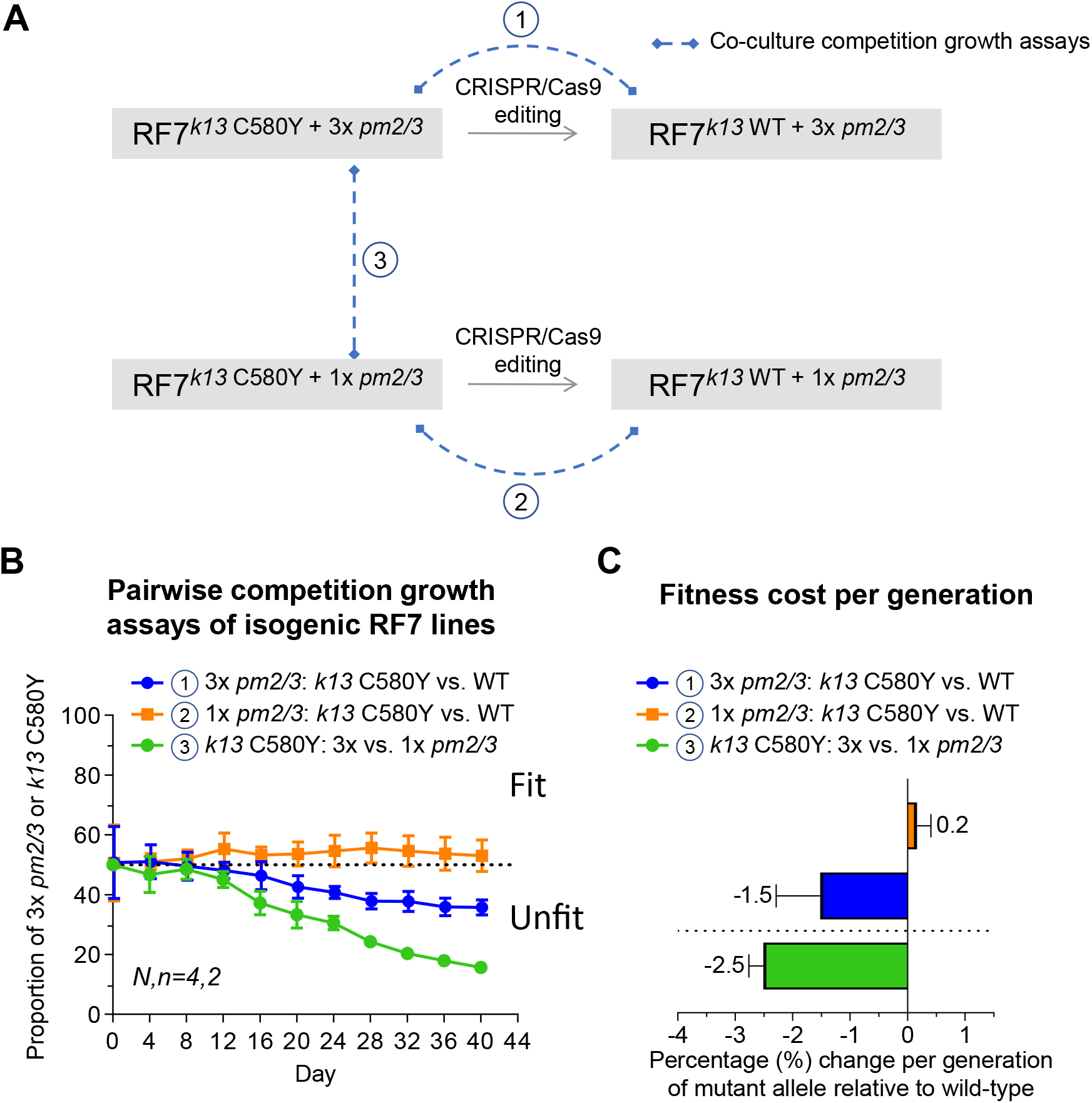
Impact of mutant *k13* and multicopy *pm2/3* on parasite’s asexual fitness. (**A**) Experimental design showing the generation of the panel of isogenic RF7 lines used in the co-culture competitive fitness assays. (**B**) Pairwise competitive growth assays showing the proportion of *k13* C580Y or WT alleles in the RF7 line harboring either 3 copies or 1 copy of *pm2/3*, and the proportion of RF7 lines carrying either 3 or 1 copy of *pm2/3* measured at 4 day intervals over 40 days. Values shown are the averaged % from four independent experiments with two technical replicates per pairwise competition assay. (**C**) Fitness cost per generation showing the relative change in percentage of the *k13* and *pm2/3* genotypes for each pairwise comparison.

### The Dd2 *pfcrt* allele increases parasite susceptibility to the ACT partner drugs LMF and MFQ

When testing other drugs, RF7 displayed 3 to 6-fold higher sensitivity to both LMF and MFQ when compared with NF54 (**Figs. 1B** and **8, A** and **B**). IC_50_ values of the progeny assayed against these two drugs were positively correlated (R=0.66), indicative of similar although not identical modes of resistance (**Fig. 8, A** and **B** and **fig. S4**). This contrasted with the negative correlations (in the range of - 0.35 to -0.45) observed between LMF and MFQ IC_50_ responses and either PPQ IC_50_ or AUC levels (**fig. S4**). This may suggest different markers of resistance and/or opposing alleles for the same set of markers. QTL analyses for LMF IC_50_ values revealed a dominant peak on chr7 within 11 kb of *pfcrt* (**Fig. 8C** and **Table S11**). For MFQ, we observed multiple QTL peaks (including the same chr7 segment), even though their LOD scores were just below the threshold for 95% significance, suggesting a multigenic trait (**Fig. 8D**). When controlling for *pm2/3* as a covariate, we observed an increased association between the chr7 *pfcrt* QTL and MFQ IC_50_ values (**fig. S9**). Interestingly, analysis of the IC_50_ values and *pfcrt* genotypes of the recombinant progeny revealed an association between mutant *pfcrt* and lower IC_50_ values for both LMF and MFQ (**Fig. 8E**).

**Fig. 8.**
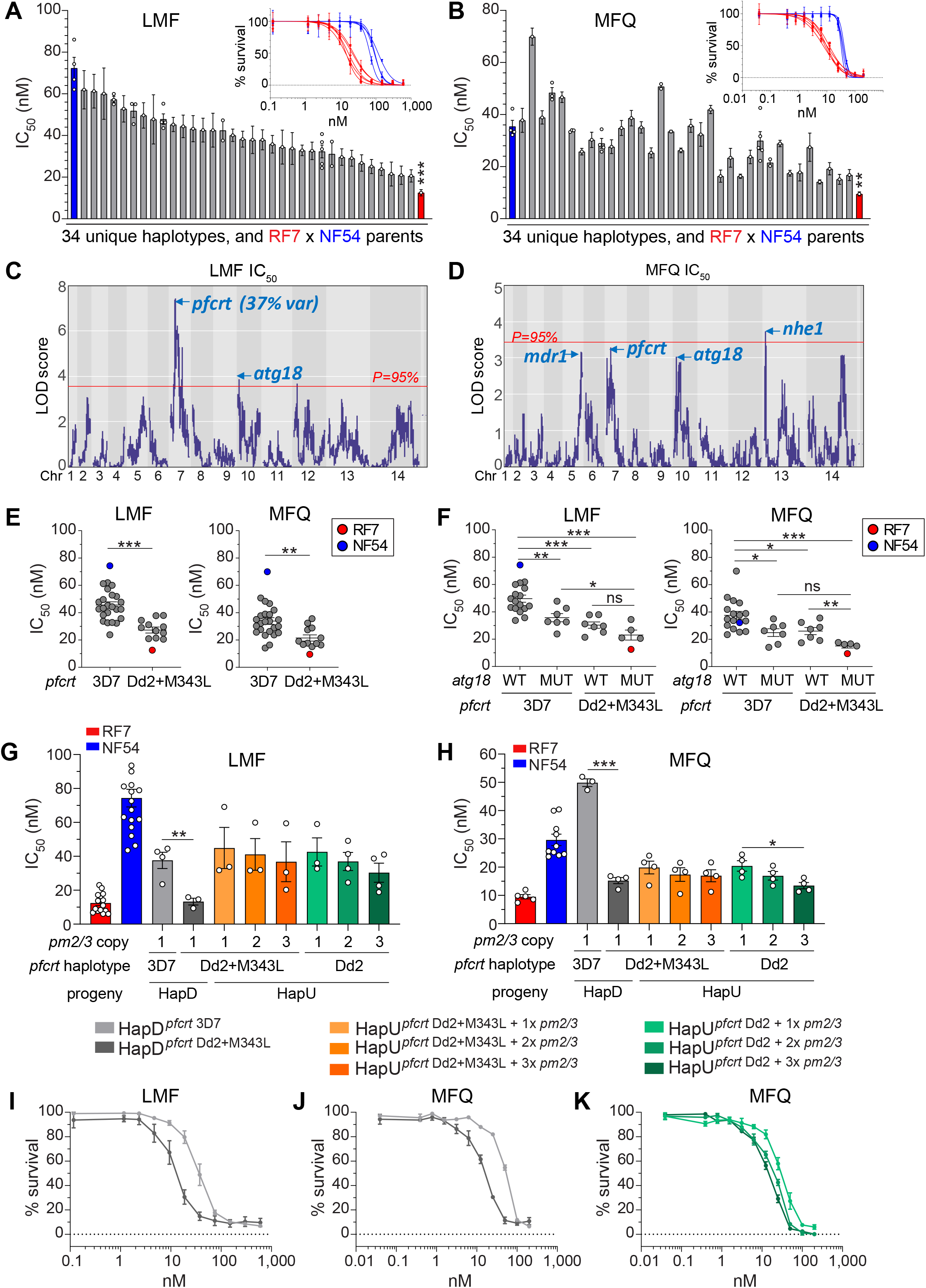
QTL mapping of LMF and MFQ response identifies multiple peaks including mutant Dd2 *pfcrt*. (**A**) LMF IC_50_ and (**B**) MFQ IC_50_ values of the 34 recombinant haplotypes and two parents. Each bar represents the mean IC_50_ ± SE for a haplotype group, and haplotypes were ordered based on descending IC_50_ levels for LMF. For certain haplotype groups in which identical clones were obtained, multiple points depict more than one sibling progeny being phenotyped. Significance between the genetic cross parents’ responses were tested by Mann-Whitney U (N,n=10-14,2 for LMF and N,n=11,2 for MFQ). \*\*\**P*<0.001. Inset panels depict dose response curves for the RF7 (red) or NF54 (blue) parents across a range of LMF and MFQ concentrations. Each curve represents the mean ± SD for an independent experiment performed in technical duplicates. (**C**) LOD plots showing significant QTLs for LMF above the 95% probability threshold (red line). (**D**) LOD plots showing multiple QTLs for MFQ falling just below the 95% probability threshold (red line). (**E**) LMF IC_50_ and MFQ IC_50_ values for the 34 independent recombinant progeny and two parents, grouped by the *pfcrt* haplotypes. Significant differences in drug response between the *pfcrt* haplotypes were tested by Mann-Whitney U. \*\**P*<0.01, \*\*\**P*<0.001. (**F**) LMF IC_50_ and MFQ IC_50_ values for the 34 independent recombinant progeny and two parents, grouped by their *atg18* genotypes and *pfcrt* haplotypes. WT: wild-type, MUT: mutant T38I. Significant differences in drug response between the recombinant groups with different *atg18* or *pfcrt* alleles were tested by Mann-Whitney U. \**P*<0.05, \*\**P*<0.01, \*\*\**P*<0.001. (**G**) LMF IC_50_ values in *pfcrt*-edited HapD progeny (3D7 vs. Dd2+M343L) having single *pm2/3* copy and in edited HapU progeny (Dd2 vs. Dd2+M343L) having either 1, 2 or 3 copies of *pm2/3*. Bars represent the means ± SE for 3-4 independent experiments performed in technical replicates. Significance between the isogenic edited progeny lines were tested using unpaired Student’s t-tests. \*\**P*<0.01. (**H**) MFQ IC_50_ values in *pfcrt*-edited HapD progeny (3D7 vs. Dd2+M343L) having single *pm2/3* copy and in edited HapU progeny (Dd2 vs. Dd2+M343L) having either 1, 2 or 3 copies of *pm2/3*. Bars represent the means ± SE for 3-4 independent experiments performed in technical replicates. Significance between the isogenic edited progeny lines were tested using unpaired Student’s t-tests. *P<0.05, \*\*\**P*<0.001. The colored key for parasites lines applies to panels **G** to **K**. (**I**) LMF and (**J**) MFQ dose response curve in edited HapD progeny showing that mutant Dd2+M343L *pfcrt* confers increased tolerance to these drugs. Each line depicts the mean % parasite survival ± SE across 3-4 independent experiments performed in technical duplicates. (**K**) MFQ dose response curve in edited HapU *pfcrt*-edited Dd2 progeny with variable *pm2/3* copies showing that single *pm2/3* copy associates with reduced MFQ sensitivity. Each line depicts the mean % parasite survival ± SE across 4 independent experiments performed in technical duplicates.

To identify secondary markers for LMF and MFQ susceptibility, we performed a QTL analysis for LMF IC_50_, controlling for *pfcrt.* These data revealed significant QTLs on chr2 for LMF and chr10 for LMF and MFQ (**fig. S9A** and **Table S11**). These segments contain *atg11* and *atg18*, respectively. Of note, the *atg18* T38I mutation (inherited from the RF7 parent) was associated with lower LMF and MFQ IC_50_ values, in both WT and mutant *pfcrt* progeny (**Fig. 8F**). Progeny harboring both mutant *atg18* and mutant *pfcrt* were the most sensitive to these two drugs, whereas the double WT haplotypes were the least sensitive (**Fig. 8F**).

We next tested whether *pfcrt* might directly affect LMF and MFQ responses by assaying *pfcrt*-edited isogenic progeny (**Fig. 8, G** and **H**). Gene*-*edited HapD progeny carrying mutant Dd2+M343L *pfcrt* showed a significant three-fold reduction in LMF and MFQ IC_50_ values compared with the WT *pfcrt* counterpart (**Fig. 8, G** to **J**). However, the *pfcrt* Dd2+M343L variant showed no change in its LMF and MFQ IC_50_ values when compared to the Dd2 isogenic comparator (**Fig. 8, G** and **H**). suggesting that the 8 amino acid Dd2 *pfcrt* allele is solely responsible for heightened sensitivity to LMF and MFQ. We note a small but significant decrease in MFQ IC_50_ with higher *pm2/3* copies in the HapU *pfcrt*-edited progeny (**Fig. 8, H** and **K)**.

### RF7 and NF54 exhibit differential susceptibility to preclinical compounds

We tested the RF7 and NF54 response to a panel of 15 clinical and preclinical antimalarial compounds to map modulators of parasite susceptibility (**Fig. 9A**). Four of these compounds showed higher IC_50_ values in RF7. These include the antifolate P218 (*37*), which targets *P. falciparum* DHFR that in the case of RF7 carries the four mutations mediating high-level resistance to the related antifolate PYM (*2*). We also observed higher IC_50_ values for GNF-Pf-5640 (*38*), previously found to select for point mutations in *pfmdr1* that in RF7 has the Y184F mutation. Four to 10-fold higher IC_50_ values were also seen for MMV665939 and MMV675939 (labelled collectively as “MMV6x”), which were previously associated with *abci3* point mutations or CNVs (*39, 40*). Previously known MMV6x resistance-conferring *abci3* mutations or CNVs were absent in RF7 (**fig. S10A**). Nonetheless, the two point mutations S2106C/S2227G in RF7 showed evidence of selection in field isolates (**fig. S10B**). This double mutant was present in ∼37% of Cambodian or Thai isolates (n=718), based on our analysis of 2,512 isolates sequenced from Asia and Africa (*41*).

**Fig. 9.**
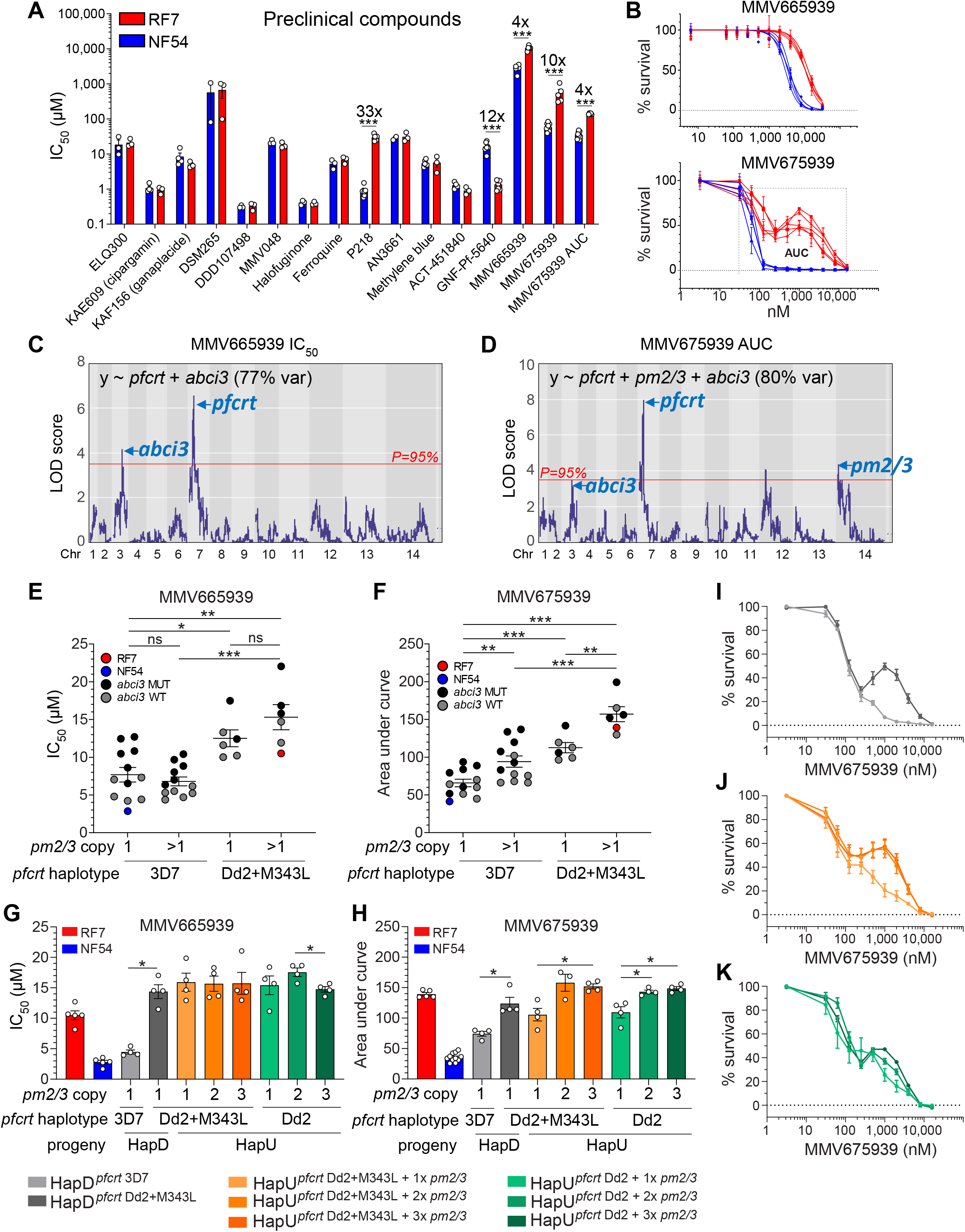
Phenotypic response of genetic cross parents, RF7 × NF54, to preclinical compounds and identification of *pfcrt, abci3, pm2/3* in MMV665939 and MMV675939 resistance. (**A**) 72h drug susceptibility IC_50_ values for RF7 x NF54 to a panel of experimental compounds. Values represent the means ± SE across 3-7 independent replicates performed in technical duplicates. Statistical significance was determined by unpaired Student’s t-tests and adjusted by multiple testing and the IC_50_ fold-shifts are indicated above the bars. \*\*\**P*<0.001. (**B**) Dose response curves for parents across a range of MMV665939 and MMV675939 showing a sigmoidal survival curve for MMV665939 and a biphasic response for MMV675939 in the drug-resistant RF7 parent. Each line represents the mean % parasite survival ± SE for one independent experiment performed in technical duplicates. N,n=4,2. (**C** and **D**) LOD plots for MMV665939 IC_50_ (**C**) and MMV675993 AUC levels (**D**) showing QTLs identified above the 95% confidence threshold. (**E**) MMV665939 IC_50_ and (**F**) MMV675939 AUC levels for the 34 independent recombinant progeny and two parents, grouped by the *pm2/3* copy, *pfcrt* and *abci3* genotypes. Significant differences in drug response between the recombinant groups harboring 1 vs. >1 *pm2/3* copies or between *pfcrt* haplotypes were tested by Mann-Whitney U. \**P*<0.05, \*\**P*<0.01, \*\*\**P*<0.001. (**G**) MMV665939 IC_50_ and (**H**) MMV675939 AUC levels in *pfcrt*-edited HapD progeny (3D7 vs. Dd2+M343L) and in edited HapU progeny (Dd2 vs. Dd2+M343L) having either 1, 2 or 3 copies of *pm2/3*. Values represent the means ± SE across 4 independent experiments performed in technical duplicates. Statistical significance was determined by Mann-Whitney U, \**P*<0.05. The colored key for parasites lines applies to panels **G** to **K**. (**I**) MMV675939 dose response curves in HapD (3D7 WT) vs. edited HapD (Dd2+M343L) progeny showing that mutant *pfcrt* is sufficient to generate a biphasic curve on a single copy *pm2/3* background. Each line depicts the mean % parasite survival ± SE across 4 independent experiments performed in technical duplicates. (**J**) MMV675939 dose response curves of HapU (Dd2+M343L) with 1-3 copies of *pm2/3* showing that multicopy *pm2/3* can augment the biphasic response. Each line depicts the mean % parasite survival ± SE across 4 independent experiments performed in technical duplicates. (**K**) MMV675939 dose response curves in edited HapU (Dd2 *pfcrt*) progeny with 1-3 copies of *pm2/3* showing that the M343L mutation is not required to generate a biphasic response on a Dd2 *pfcrt* background. Each line depicts the mean % parasite survival ± SE across 4 independent experiments performed in technical duplicates.

#### QTL mapping identifies pfcrt as a major determinant of reduced susceptibility to MMV6x compounds

To map determinants of resistance to these MMV6x compounds, we phenotyped the cross progeny and performed QTL analysis (**fig. S11A** and **Fig. 9B**). QTL analysis showed that a narrow chr7 peak harboring *pfcrt* was the strongest predictor of MMV665939 IC_50_ and MMV675939 AUC levels (**Fig. 9, C** and **D, Table S12**). For both compounds, we also noted a smaller secondary QTL on chr3 containing the *abci3* locus. No other QTL peaks were observed for MMV665939 when controlling for *pfcrt* or *pm2/3* (**fig. S9B**). This observation suggests that MMV665939 resistance is associated with chr7 and chr3 segments, likely *pfcrt* and *abci3*. For MMV675939, we observed an additional peak on chr 14 that included *pm2/3* (**Fig. 9D** and **fig. S9B**). These data suggest that these three loci combine to underpin differential MMV675939 susceptibility.

#### Multicopy pm2/3, abci3 and pfcrt are combinatorial in modulating parasite susceptibility to MMV6x compounds

In our progeny, higher MMV665939 IC_50_ values and MMV675939 AUC levels were associated with both mutant *pfcrt* and *abci3* (**Fig. 9, E** and **F**). Higher MMV675939 AUC levels were also observed in progeny harboring multicopy *pm2/3* (**Fig. 9F**). For MMV675939, biphasic responses were observed either in progeny harboring mutant *pfcrt* as well as in progeny harboring mutant *abci3* in the presence of multicopy *pm2/3* (**fig. S12**). Gene editing of HapD by replacing WT with mutant *pfcrt* confirmed the role of this gene in MMV6x response (**Fig. 9, G** and **H**) and resulted in a MMV675939 biphasic response (**Fig. 9I**). Amplified *pm2/3* decreased parasite susceptibility to MMV675939 in HapU and HapU edited lines, whereas MMV665939 was unaffected (**Fig. 9, G** and **H**). Genetically-edited HapU parasites showed that the *pfcrt* M343L mutation had no impact on either MMV6x compound (**Fig. 9, G** and **H**), leading us to conclude that the Dd2 *pfcrt* allele itself can cause reduced susceptibility to MMV6x compounds and produce a biphasic response to MMV675939 (**Fig. 9, J** and **K**).

#### MMV6x compounds can interfere with mutant PfCRT-mediated efflux of PPQ and CQ

To investigate whether resistance to the two MMV compounds is mediated by PfCRT transport, we performed competitive inhibition assays to test their ability to inhibit CQ or PPQ uptake using PfCRT-loaded proteoliposomes (*28*). Both compounds competed with both PPQ and CQ for mutant PfCRT-mediated drug uptake, suggesting that the two MMV6x compounds can interact directly with PfCRT (**fig. S11B**).

## Discussion

By leveraging a new *P. falciparum* genetic cross, we identified parasite loci associated with resistance to multiple first-line antimalarial drugs. Our studies of ART resistance identify two unlinked chromosomal segments that appear to contribute to ART resistance, in addition to mutant *k13* that our genotype-phenotype mapping localized as the primary determinant. We also report that PPQ resistance is dictated by an epistatic relationship between mutant *pfcrt* and multicopy *pm2/3*, which together achieve high-level resistance but with a fitness cost that leaves them vulnerable to replacement with WT *pfcrt* or single-copy *pm2/3* in the absence of PPQ pressure. We also show that the first-line partner drug LMF, and its related arylaminoalcohol MFQ, select against mutant *pfcrt*, underscoring a strategic advantage of implementing LMF and PPQ in the same field settings. Finally, our data suggest that mutations in *abci3* form a network with *pfcrt* and multicopy *pm2/3* in regulating parasite susceptibility to diverse chemotypes, as illustrated with the two antiplasmodial drug discovery candidates MMV665939 and MMV675939.

Recent reports that *k13* mutations have emerged in several sites in east Africa and the Horn of Africa are a cause for concern, as reduced ART efficacy increases selective pressure on the partner drug and narrows the horizon towards treatment failures (*42*). Intravenous artesunate is also first-line treatment for severe malaria (*43*), with mutant *k13*-mediated delayed parasite clearance potentially compromising treatment efficacy. Mutant *k13* parasites in the GMS are often more ART-resistant and display improved fitness compared with African strains, suggesting that Asian strains have acquired secondary determinants (*12, 22*). Our genetic cross data identify two candidate loci – a chr1 segment harboring *mrp1* and a chr14 segment harboring *arps10*. The four *mrp1* mutations present in RF7 (H191Y, S437A, I876V, F1390I) are quite prevalent in the GMS and have been associated with altered susceptibility to ART derivatives (*44, 45*). The most common of these mutations in Africa, I876V, was earlier associated with parasites recrudescence following artemether-LMF treatment (*46*). This transporter is non-essential in *P. falciparum* ABS parasites cultured *in vitro*, with the knockout parasites showing increased parasite susceptibility to ART (*47*). In our genetic cross progeny, mutant *mrp1* was only found in parasites harboring mutant *k13*.

Our *arps10* marker on chr14, which harbors the V127M+D128H mutations, has been previously associated with prolonged parasite clearance half-lives following artesunate treatment and with increased RSA survival in K13 mutant parasites (*48, 49*). This gene is thought to be a key founder mutation of the KEL1/PLA1 lineage on which ART resistance emerged in the GMS (*48*). Other mutations in this chr14 segment include exonuclease-V R25K and the alpha/beta hydrolase I301F, which were earlier identified as a potential markers of DHA-PPQ treatment failures in Cambodian isolates (*24*). It is noteworthy that this segment overlaps the QTL associated with enhanced fitness from separate genetic cross studies with GMS parasites (*50*). These genes on chr 1 and 14 merit further examination as potential contributors to ART resistance emerging in African settings.

Our PPQ studies showed unexpected connections to ART resistance. Upon applying PPQ or CQ pressure to bulk progeny, we recovered resistant *pfcrt*-mutant progeny that were all mutant *k13*. PPQ also selected for the ART-resistant chr14 segment. These observations suggest that the selection of *pfcrt* mutants by PPQ in the field might have evolved on the background of mutant *k13* and favored the inheritance of the chr14 segment which can augment DHA resistance. Our results provide support that PPQ may have contributed to the dominance of the combined KEL1/PLA1 (*k13* C580Y and multicopy *pm2/3*) co-lineage across the GMS where DHA-PPQ is used (*51*).

Our PPQ studies resolve debate about the relative roles of mutant *pfcrt* and multicopy *pm2/3* in driving resistance to this drug (*27, 31, 51*). At PPQ concentrations <100 nM, we observed a decrease in parasite susceptibility upon amplification of *pm2/3*, in isogenic progeny expressing either Dd2+M343L or Dd2 PfCRT isoforms. At 200 nM PPQ or above, the Dd2+M343L variant afforded a clear gain of PPQ resistance which increased with increasing *pm2/3* copy number. The biphasic nature of the PPQ dose-response curve was most pronounced with multicopy *pm2/3* on a mutant *pfcrt* background. The PPQ C_max_ in plasma is ∼200 nM (*52*), which based on our data is consistent with both determinants contributing to resistance. The long terminal half-life of PPQ (∼3 weeks) would also explain a selection advantage for multicopy *pm2/3*, which we found to enhance parasite survival at low PPQ concentrations. This partial gain of resistance likely favored the emergence of novel PfCRT variants that dominate PPQ resistance at higher PPQ concentrations. Biophysical studies attribute this to a mutant PfCRT-mediated gain of PPQ efflux away from its heme target in the digestive vacuole (*28*). For *pm2/3*, the basis for resistance might relate to its impact on the rate of hemoglobin proteolysis or conversion of heme to hemozoin, effectively depleting levels of heme that act as the receptor for PPQ binding. The earlier observation that *pm2* and *pm3* form part of a multi-component hemozoin formation complex (*53*) may explain why amplification of both tandem genes is associated with PPQ resistance yet overexpression of each gene individually did not affect the PPQ dose-response profile (*54*). The interconnection between these mediators and ART resistance presumably centers on heme.

Our studies revealed that multicopy *pm2/3* exerts a significant fitness cost in the Cambodian RF7 *k13* C580Y parasite, exceeding that observed with the *k13* mutation itself. We also observed spontaneous deamplification of *pm2/3* in parasites cultured for two or more months in the absence of PPQ selection. PPQ resistance-associated mutant PfCRT haplotypes are also known to exert a fitness cost (*27, 55, 56*). This growth defect has been attributed to an excessive buildup of globin-derived short peptides in the digestive vacuole, which serve as a natural substrate, thereby restricting protein synthesis (*57–59*). These observations help explain why the prevalence of both multicopy *pm2/3* and PPQ-resistant mutant *pfcrt* decreased rapidly following withdrawal of DHA-PPQ from first-line use in Cambodia (*60*).

In the GMS, DHA-PPQ has been largely replaced by artemether-LMF, which to date has remained effective as LMF has not encountered resistance. Artesunate-MFQ has also been widely implemented since the early 2000s. Our QTL analysis data show a strong association between LMF or MFQ IC_50_ values and *pfcrt*, with these drugs selecting against the Southeast Asian mutant Dd2+M343L *pfcrt*. We confirmed these results in *pfcrt* gene-edited parasites. These data provide evidence that LMF and PPQ can exert opposing selective pressures on the PfCRT locus, in agreement with earlier observations that LMF selects against CQ-resistant *pfcrt* as detected using the K76T marker (*61, 62*).

In Africa, artemether-LMF accounts for 75% of the antimalarial drug market (*1*). Recent studies have shown excellent efficacy of DHA-PPQ either for intermittent preventive treatment of infants or pregnant women or for seasonal malaria chemoprevention (*63*). Our data lend support to the introduction of DHA-PPQ into areas with artemether-LMF use. Clinical teams have also documented excellent clinical efficacy of triple therapies that combine DHA-PPQ with MFQ in the GMS, with the rationale that use of these triple ACTs can suppress the emergence of multidrug resistance (*64*). Interestingly, we observed a slight sensitization of our multicopy *pm2/3* progeny towards MFQ, a drug that is known to select for multicopy *pfmdr1.* However, no association was observed with LMF. These data may help explain why *P. falciparum* parasites in the GMS very rarely show amplification of both *pfmdr1* and *pm2/3* (*60*).

Our genetic mapping of candidate loci is constrained by the choice of resistant parent and the resolution of the genetic crossover events. Nonetheless, our data illustrate the power of genetic crosses for identifying polygenic networks of *P. falciparum* multidrug resistance. Crosses provide an important complement to genome-wide association studies as well as *in vitro* evolution and whole-genome sequence studies that can identify candidate chromosomal segments and specific loci (*23, 24, 39, 65*). These approaches will be key to identifying the genetic basis of *P. falciparum* resistance to first-line combination therapies, which appears to be imminent in Africa, and leveraging that knowledge to map the spread of resistance and develop strategies to mitigate its impact.

## Materials and Methods

### *In vitro P. falciparum* cell culture

Asexual blood stage *P. falciparum* parasites were cultured in human O^+^ RBCs obtained from the Interstate Blood Bank (Memphis, TN) at 3% hematocrit, using RPMI 1640 medium supplemented with 25 mM HEPES, 2.1 g/liter sodium bicarbonate, 10 μg/ml gentamicin, 50 μM hypoxanthine, 0.5% (wt/vol) AlbuMAX II (Thermo Fisher Scientific), and 10% (v/v) human serum (Interstate Blood Bank). Parasites were maintained at 37°C under 5% O_2_/5% CO_2_/90% N_2_ gas conditions. The RF7 and NF54 parasites were obtained respectively from Drs Rick Fairhurst and Chanaki Amaratunga (NIAID, NIH) and Dr Photini Sinnis (Johns Hopkins Bloomberg School of Public Health). The RF7 (Sanger ID: PH1008-C (*27*)) isolate was a recrudescent sample of PH1356-C (genetically identical except that the latter had only a single copy of *pm2/3*), which was collected on day 0 of patient admission. Single clones of the RF7 parent and HapU progeny were obtained by cloning at 0.5 parasite/well using the limiting dilution technique. Media was replenished twice a week and fresh blood was added once per week. Clones were expanded on day 20-24 after parasitemias were above the detection threshold and the *pm2* copy number was measured by TaqMan-based real-time PCR on isolated gDNA (see section below).

### *P. falciparum* infections in *An. stephensi* mosquitoes and humanized chimeric liver mice

#### Gametocyte cultures and mosquito feeding

RF7 and NF54 gametocytes were cultured as previously described (*66*). Briefly, parasites were grown in RPMI 1640 containing 10% v/v human serum and 4% hematocrit at 37°C in a glass candle jar. Cultures were initiated at 0.5% asynchronous asexual parasitemia from a low-passage stock and maintained up to day 18 with daily media changes but without any addition of fresh erythrocytes. Day 15 to 18 cultures, containing largely mature gametocytes, were used for mosquito feeds. Cultures were centrifuged (108 x *g,* 4 min) and the parasite pellet was resuspended to 0.6% gametocytemia in a mixture of human O^+^ RBCs supplemented with 50% v/v human serum. To genetically cross RF7 and NF54, we used a 1:1 ratio of gametocytes at a final gametocytemia of 0.6%. Gametocytes were fed to 2 to 4-day old *Anopheles stephensi* mosquitoes (Liston strain) that had been starved overnight, using glass membrane feeders. Unfed mosquitoes were removed after feeding. Fed mosquitoes were then maintained on a 10% sugar solution at 25°C and 80% humidity with a 14: 10h (light: dark) cycle including a 2h dawn/dusk transition. Human erythrocytes used to set up the cultures were collected weekly from healthy volunteers, with informed consent, under a Johns Hopkins University Institutional Review Board approved protocol number NA_00019050. All experiments were performed in accordance with institutional guidelines and regulations.

#### Oocyst and salivary gland sporozoite quantification

On day 7 post-infected blood meal, mosquito midguts were harvested and stained with mercurochrome. Oocysts were counted by brightfield microscopy using a Nikon E600 microscope with a PlanApo 10× objective. We observed a 75% prevalence of infection, with an average of 10.7 oocysts per mosquito. Between days 14 to 16, salivary glands were harvested from a minimum of 20 mosquitoes, homogenized and the sporozoites counted using a hemocytometer. On average we obtained 25,000 salivary gland sporozoites per mosquito.

#### Mouse infections

All animal experiments were performed in accordance with the Animal Care and Use Committee (ACUC) guidelines and approved by the Johns Hopkins ACUC (Protocol M017H325). Hu-Hep mice were purchased and shipped from Yecuris. Upon arrival we began withdrawal of the mice from 2-(2-nitro-4-trifluoromethylbenzoyl)-1,3-cyclohexanedione (NTBC). Mice were kept in a sterile facility with sterile bedding, food, and water. Mice were weighed daily and those that showed any weight loss >10% (compared to pre-shipment weight) were treated with 300 µL of sterile saline via intraperitoneal injection and given a nutritional supplement in water (STAT; https://www.prnpharmacal.com/products/nutritional-supplements/stat/). Mice were allowed to recover from shipping for 2 to 3 days prior to infection. Infection with *P. falciparum* sporozoites was performed either by mosquito bite or by intravenous (IV) inoculation. For the mosquito bite approach, mosquitoes were allowed to probe/feed on the isoflurane-anaesthetized mice for 15 to 20 min. Each of the three mice received approximately 50 to 60 infectious mosquito bites. A fourth mouse was IV-inoculated with 3 million sporozoites dissected from mosquito salivary glands. Mice infected by mosquito bite remained stable and healthy-appearing for the remainder of the experiment. In contrast, the IV-inoculated mouse lost >10% body weight and on day four post-inoculation appeared ill. This latter mouse was given 3,000U penicillin via the intramuscular route, 1000U penicillin and 1 mg streptomycin intraperitoneally, with additional daily doses of penicillin and streptomycin until sacrifice. On days 6 and 7 post-sporozoite infection, 500 µL of RPMI-washed human RBCs were inoculated IV into each mouse. At day 7.5 post-infection, ABS parasites were detected at up to 0.3% parasitemia and blood recovered by cardiac puncture of mice anesthetized with ketamine/xylazine. One mouse, M46056, which received 3 million sporozoites via the IV route, corresponded to the highest parasitemia on day 7 and was used for bulk selection experiments (**Table S2**).

### WGS of bulk pools and individual progeny

Parasite cultures at 1-5% parasitemia were lysed with 0.1% saponin in 1× PBS, and genomic DNA was then extracted using the QIAamp DNA Blood Mini Kit (Qiagen). DNA concentrations were determined using the Qubit dsDNA BR assay kit (Thermo Fisher Scientific). WGS libraries were prepared using the Illumina Nextera DNA Flex protocol. Briefly, 150 ng of gDNA was fragmented and tagmented using bead-linked transposons and subsequently amplified by five cycles of PCR to add dual-index adapter sequences to the DNA fragments. The libraries were quantified, pooled, and sequenced on Illumina MiSeq or NextSeq platforms to generate 300 bp or 150 bp paired-end reads, respectively. Drug-selected bulk cultures yielded a mean 44× depth of coverage by Illumina MiSeq sequencing. Cloned progeny were subjected to WGS by Illumina NextSeq or MiSeq to identify selfed vs. genetic recombinants. Thereafter, representative progeny from each of the 34 recombinant haplotypes, as well as the two parents RF7 and NF54, were resequenced on the Illumina MiSeq platform to obtain between 2.7M to 4.6M reads, corresponding to an average of 40.7× depth of coverage, with on average at least 94.1% of each genome having 10 or more reads (**Table S3**). Sequence data were aligned to the *P. falciparum* 3D7 genome (PlasmoDB version 36.0) using Burrow-Wheeler Alignment (BWA). Reads that did not map to the reference genome and PCR duplicates were removed using SAMtools and Picard. The reads were realigned around indels using Genome Analyses Tool Kit (GATK) RealignerTargetCreator and base quality scores were recalibrated using GATK BaseRecalibrator.

All variants were called using *mpileup* and filtered based on quality scores (minimum base quality score ≥20, mapping quality >30, read depth ≥5) and multiallelic = FALSE to obtain high-quality SNPs, which were then annotated with snpEFF. SNPs for highly polymorphic surface antigens and multi-gene families (notably var, stevor and rifin, located mostly at the sub-telomeric ends of chromosomes) were as these are prone to stochastic changes during *in vitro* culture. To obtain the list of SNPs that differed between the parents in the core genome we retained only the SNPs that were homozygous, based on >90% alternate allele frequency in RF7 and >90% reference allele frequency in NF54. For QTL mapping, *pm2/3* copy was manually imputed as an additional marker “Pf3D7_14_v3_295261_PM2”. This analysis gave a total of 18,490 SNPs (“18k” SNP set). SAMTools mpileup was used to find SNPs in the “18k” SNP set for progeny or in targeted genomic regions including genes mediating known drug resistance phenotypes. SNPs that showed an allele frequency of >10% and <90% in the progeny clones were classified as heterozygous and defined as “missing/undetermined”. BIC-Seq was used to check for copy number variations using the Bayesian statistical model (*67*).

### Microsatellites for identifying recombinant progeny and for determining PfPailin genotypes within the *k13*-flanking regions

To establish a list of microsatellites (MS) that can be used to differentiate the RF7 and NF54 parents and provide a MS signature for each recombinant progeny, a custom Python script utilizing the pysam module was written to call WGS reads that harbored insertions or deletions at specified genomic loci within the respective windows, as described previously (*68*). Integrated Genome Viewer was used to visually verify the presence of MS. The MS markers used herein were C2M18, TAA81, BM5, C3M67, TA1 and C13M87; **fig. S13)**. To identify MS sizes surrounding the *k13* gene, we referred to the MS sizes previously profiled in PfPailin parasites that spanned -50.0 kb to +31.5 kb of the *k13* gene (*5, 69*), and determined the sizes of these MS in the progeny based on the WGS reads (**fig. S3**).

### Confirmation of progeny clones via microsatellite-based multiplexed fragment analysis

Genotyping to validate the genetic purity of individual progeny clones, pre and post phenotyping, was performed using multiplexed fragment analysis (**fig. S13**). PCRs were set up individually for C2M18, TAA81, BM5, C3M67, TA1 and C13M87, using forward primers fluorescently labeled with 6-FAM (blue), ATTO550/NED (yellow), or ATTO532/VIC (green) as described previously (*68*). PCR products were treated with ExoSAP-IT Express Reagent (Thermo Fisher Scientific) and sent for capillary electrophoresis with the Liz500 ladder as reference (Genewiz). Data were analyzed using Peak Scanner 2 (Thermo Fisher Scientific) to determine the microsatellite size of the fragment corresponding to either RF7 or NF54, which were run as parallel controls.

### Phenotypic assessment of drug sensitivities in genetic cross parents and progeny clones

#### Drug susceptibility assays for RF7 × NF54 parents

To profile the genetic cross parents’ susceptibility to a panel of 30 antimalarial drugs and preclinical compounds, we performed 72h dose-response assays on sorbitol-synchronized ring-stage cultures in at least 3 to 17 independent experiments with technical replicates. Assays employed a 10-point 2-fold serial dilution of the compounds and 0.1% (dimethyl sulfoxide) DMSO control-treated samples were included.

#### DHA ring-stage survival and drug susceptibility assays

To obtain tightly synchronized parasites for DHA ring-stage survival assays (RSAs), cultures were treated with 5% D-Sorbitol (Sigma) for 15 min at 37°C to remove mature parasites. After 40h, late-stage segmented schizonts were purified over 35% and 65% Percoll double density gradients, as described previously (*70*). Purified schizonts were incubated with fresh RBCs for 3h and treated with 5% D-Sorbitol to obtain 0 to 3-hour post invasion (hpi) early rings. Beginning with 1% parasitemia and 1% hematocrit, these early rings were exposed to DHA concentrations ranging from 5.5 nM to 2,800 nM, prepared as 2-fold serial dilutions or to 0.1% DMSO (vehicle control) in 96-well plates, with total culture volumes of 200 μL per well and technical duplicates. After a 4h incubation at 37°C, the wells were washed three times with complete medium to remove drug and transferred to new 96-well plates using the Freedom EVO MCA96 liquid-handling instrument (Tecan). Cultures were subsequently maintained for an additional 66h in drug-free medium.

#### PPQ, LMF, MFQ, MMV665939 and MMV675939 drug susceptibility assays

Synchronized ring-stage parasites were exposed for 72h to PPQ (dissolved in 0.5% lactic acid) at 2-fold serially diluted concentrations ranging from 0.8 nM to 25.6 μM, alongside drug-free wells, in technical duplicates. Standardized 72h dose-response assays were conducted similarly for LMF, MFQ, MMV665939, MMV675939 at 10-point concentrations starting from 600 nM, 200 nM, 32 μM, and 16 μM for these compounds, respectively. 0.1% DMSO-treated control wells were run in parallel. For all drug assays, the RF7 and NF54 parents were included as reference controls.

#### Flow cytometry for quantification of viable parasites

Parasitemias for drug-treated and vehicle control-treated wells were measured at 72h by flow cytometry, as described previous (*15*). Briefly, parasites were incubated with 1× SYBR Green (ThermoFisher) and 100 nM MitoTracker DeepRed (ThermoFisher) for 30 min at 37°C and quenched with 1× PBS. On average, we analyzed 10,000 cells per sample, using an iQue Screener Plus cytometer (Sartorius). Viable parasites were defined as the percentage of MitoTracker-positive and SYBR Green-positive labelled cells.

#### Calculation of RSA, PSA, AUC and IC_50_ values

For all drug assays conducted herein, we included kill controls in which 1 μM DHA-treated parasites were used as a background control to achieve complete parasite killing and this percent parasitemia was subtracted from the parasitemias measured for each well. Parasite survival in the presence of DHA or PPQ was expressed as the percentage of the background-subtracted parasitemias in the 700 nM DHA-treated samples or 200 nM PPQ-treated samples divided by the parasitemias of the DMSO-treated samples. Mean RSA survival rates >1% were defined as DHA resistant (*71*). The Area Under the Curve (AUC) values for DHA, PPQ and MMV675939 were calculated based on total parasite survival across the range of 21.9 nM to 2.8 μM for DHA, 3.1 nM to 25.6 μM for PPQ, and 31.3 nM to 16 μM for MMV675939. IC_50_ values were determined by applying a non-linear regression model (sigmoidal dose-response with variable slope) on the normalized % survival across the log-transformed drug concentrations, using Prism v8.3.1 (GraphPad).

### Genetic map construction

Duplicate markers were removed to obtain 2,091 markers using R/qtl. These markers were analyzed in JoinMap v5 (Kyazma B.V.) using the independence LOD parameter to generate a single linkage group for each chromosome, except for chr 12 where there were three linkage groups. Using the resulting 1,918 markers, we constructed a genetic map for RF7 x NF54 using Kosambi’s regression mapping function and adjusted for linkages with a recombination frequency < 0.5 and LOD scores >0 (**Table S6**).

### Bulk segregant analysis of bulk pools and drug-selected clones

Identification of QTLs in bulk pools treated with each of the drug conditions was conducted on the “18k” SNP set obtained from WGS data using G’ and QTL-seq approaches in the QTLseqr package in R. A sliding window of 100 kb was used for calculating the tricube-smoothed G’ and Δ(SNP-index) values of each SNP within that window. False discovery *q*-values were adjusted for each pairwise analysis to account for the differences in sequencing coverage. Outlier SNPs were filtered by Hampel’s rule. We also performed bulk segregant analysis on bulk progeny pools that were exposed to drug and then subsequently cloned. Progeny clones selected by PPQ, mdCQ, PYM or untreated controls (N=12 for untreated, N=12 for PYM, N=7 for mdCQ, N=6 for PPQ) were grouped and the RF7 allele frequency within each group of clones was averaged, thereby generating “pooled” clones that was used for bulk segregant analysis as shown in **Fig. 3E**.

### Individual clone-based quantitative trait loci mapping

After filtering out SNPs that were missing in >5 out of the 49 progeny that had been profiled for drug susceptibilities, we retained 15,869 SNPs and used these with the R/qtl v2 package to map QTL peaks. To identify significant QTLs for each phenotypic output, permutations of phenotypic data was performed 1,000 times to obtain a distribution of maximum LOD scores. These scores were then used to calculate the LOD threshold at 95% probability. Fine-mapping of the QTL segments was subsequently performed using Bayesian interval mapping at a 95% confidence level. To elucidate secondary QTLs, LOD scores were calculated after setting the major peak as an additive covariate and the LOD threshold at 95% probability was recalculated for this new model. The *k13*, *pfcrt* and *pm2/3* genes were applied separately as additive covariates for each of the QTL analyses. The percent variance associated with each QTL was determined by establishing linear models for each QTL and comparing additive vs. interactive QTL effects in R/qtl v1.

### Gene editing of *pfcrt* and *k13* by ZFN-based and/or CRISPR/Cas9 approaches

CRISPR/Cas9 editing to remove the C580Y mutation from the *k13* locus in RF7 clones harboring either 1 or 3 copies of *pm2/3,* using the all-in-one plasmid, pDC2-cam-coSpCas9-U6-gRNA-k13_bsm-hdhfr was performed as previously described (*12*). Zinc-finger nuclease-mediated editing to replace the 3D7 WT *pfcrt* allele with the Dd2+M343L *pfcrt* isoform (carrying 9 point mutations) was achieved in the HapD progeny as previously described (*72*). CRISPR/Cas9 editing of *pfcrt* to remove the M343L single point mutation in the HapU progeny harboring either 1, 2 or 3 copies of *pm2/3* was performed using the all-in-one plasmid, pDC2-cam-Cas9-U6-gRNA-pfcrt_ M343-bsd. Cloning was performed as for *k13*, except for gRNA cloning that was performed using In-Fusion cloning (Takara) and the drug-selection cassette of blasticidin instead of human *dhfr* was used. Cloning of gRNAs was performed using primer pair p8234+p8235. Donor templates were amplified and cloned into the final vector using the primer pairs p8236+p8237. Site-directed mutagenesis was performed using the allele-specific primer pairs p8238+p8239. All final plasmids were sequenced using the primer pair p7949+p7950 with internal sequencing primer p7989. The primer sequences are listed in **Table S13**.

To generate gene-edited lines, ring-stage parasites at 5-10% parasitemias were electroporated with 50 μg of circular plasmid DNA resuspended in Cytomix. Transfected parasites were selected by culturing in the presence of WR99210 (Jacobus Pharmaceuticals) for 6-8 days post electroporation. The RF7 clones were subjected to 5 nM WR99210, while the HapD and HapU parasites were selected under 2.5 nM WR99210 or 2 μg/mL blasticidin. Parasite cultures were monitored for recrudescence by microscopy for up eight weeks post electroporation. To screen for successful editing, the *k13* and *pfcrt* loci was amplified directly from recrudesced infected blood using the MyTaq Blood-PCR Kit (Bioline) (*68*). PCR products were submitted for Sanger sequencing and positively-edited bulk transfectants were cloned by limiting dilution. All gene-edited transgenic lines generated and their phenotypes are described in **Table S8**.

### Quantitative PCR for determination of *pm2* copy number

Multiplexed TaqMan qPCR of *pm2* labelled with FAM and single-copy *β-tubulin* labelled with HEX (as an internal control) were performed on gDNA extracted from the two parents, progeny clones and gene-edited lines, pre and post phenotyping, on a QuantStudio 3 real-time PCR system (Thermo Fisher Scientific), as described previously (*68*). Five standards of gene fragments, mixed at 1:1, 2:1, 3:1, 4:1, and 5:1 molar ratios of *pm2*:*β-tubulin*, as well as the 3D7 line (1 copy of *pm2*), were included as copy number controls. Each sample was assayed in technical duplicates. The *pm2* copy number for each progeny line was calculated by normalizing to the 3D7 control, using the relative quantification method (*68*). The primer sequences are listed in **Table S13**.

### Competitive fitness assays for *k13* and *pm2/3*

#### Isogenic co-cultures

Fitness assays were performed by co-culturing isogenic parasite lines in 1:1 ratios. These assays paired isogenic *k13* WT vs. mutant clones in RF7 clones that harbored either 1 or 3 copies of *pm2/3.* We also paired single vs. multicopy *pm2/3* RF7 clones. These three pairwise co-cultures were used to independently examine *k13* genotype or *pm2/3* copy number. Assays were initiated with tightly synchronized rings and conducted on four independent occasions with technical replicates. Cultures were maintained at 6 mL volumes over a period of 40 days (∼20 generations), and every four days a fraction of each co-culture was harvested for gDNA as described above.

#### k13 allelic discrimination qPCR assays

The percentage of WT or mutant *k13* alleles in each sample was determined using custom TaqMan fluorescence-labelled minor groove binder (MGB) probes in TaqMan allelic discrimination real-time PCR assays, as described previously (*12*). Probes were designed to specifically detect either the *k13* C580Y propeller mutation (HEX probe, Eurofins) or the WT allele (FAM probe, Eurofins). The primer sequences are listed in **Table S13**. Mixtures of WT and mutant plasmids in fixed ratios (0:100, 20:80, 40:60, 50:50, 60:40, 80:20, 100:0) were run as parallel controls to ensure accurate allelic detection. qPCR reactions for each sample were run in duplicate, with each 20 μL reaction consisting of 1× QuantiFAST reaction mix containing ROX reference dye (Qiagen), 0.66 µM forward and reverse primers, 0.16 µM FAM-MGB and HEX-MGB TaqMan probes, and 10 ng genomic DNA. Cycling conditions were as follows: 1 cycle of 30 s at 60°C and 5 min at 95°C; and 40 cycles of 30 s at 95°C and 1 min at 60°C. Every assay included no-template negative controls as well as positive controls (mixtures of WT and mutant plasmids in fixed ratios), which were run in triplicates. Amplification and detection of fluorescence were carried out on a QuantStudio3 real-time PCR system (Thermo Fisher Scientific) using the genotyping assay mode. Rn, the fluorescence of the FAM or HEX probe, was normalized to the fluorescence signal of the ROX reporter dye. Background-normalized fluorescence (Rn minus baseline, or ΔRn) was calculated as a function of cycle number.

#### pm2/3 allelic discrimination qPCR assays

TaqMan allelic discrimination was not suitable to quantify the percentage of single vs. two or more copies of *pm2/3*. To facilitate the differentiation of RF7 clones expressing 1 or 3 copies of *pm2/*3, we therefore conducted WGS of *pm2/3* multicopy versus single-copy RF7 clones. This analysis identified a SNP (gaA/gaT) present within the isoleucine-tRNA ligase gene (Pf3D7_1332900), corresponding to an E459D mutation, in the RF7 clone harboring a three-copy *pm2/3* amplicon, presumably as a result of genetic drift during extended culture and clonal expansion. Forward and reverse PCR primers, and custom TaqMan fluorescence-labelled probes were designed to target the E459 WT (FAM, Eurofins) or E459D mutant (HEX, Eurofins) alleles for this gene as listed (variant codon is underlined). The primer sequences are listed in **Table S13**.

The efficiency and sensitivity of these TaqMan primers were assessed using standard curves comprising 10-fold serially diluted templates ranging from 10 ng to 0.001 ng. Robustness was demonstrated by high efficiency (94-96%) and R^2^ values (0.99-1.00). The quantitative accuracy in genotype calling was assessed by performing multiplex qPCR assays using mixtures of WT and mutant plasmids in pre- defined fixed ratios (0:100, 20:80, 40:60, 50:50, 60:40, 80:20, and 100:0). These mixtures were obtained by cloning a ∼800 bp fragment for isoleucine-tRNA ligase centered around the mutant or WT SNP in pGEM plasmids. Triplicate data points clustered tightly and gave expected ratios of WT to mutant alleles, indicating high reproducibility in the data across the fitted curve (R^2^=0.86-0.87). Co-cultures consisting of the RF7 isogenic clones containing multicopy *pm2/3* with the E459D mutation, and the single copy *pm2/3* and WT E459 allele, were initiated at a 1:1 ratio and fitness assays conducted over a 40-day period as per the *k13* fitness experiment described above. qPCR reactions were run with probes targeting the isoleucine tRNA ligase marker that was diagnostic for single versus multicopy *pm2/3*, under the same conditions described above.

To determine the WT or mutant allele frequency in each sample, we retained only ΔRn values in samples where the threshold cycle (C_t_) was at least three cycles less than the no-template control. Next, we subtracted the ΔRn of samples from the ΔRn of the no-template negative control. We subsequently normalized the fluorescence to 100% using the positive control plasmids (tested in parallel reactions) to obtain the percentage of the WT and mutant alleles for each sample. The final percentage of the mutant allele was defined as the average of two values: the normalized percentage of the mutant allele, and 100% minus the normalized percentage of the WT allele.

#### Fitness cost assessment

The fitness cost associated with the *k13* mutation or multicopy *pm2/3* was calculated relative to its isogenic WT or single-copy counterpart, respectively, using the following equation: P’ = P*((1-x)^n^). P’ is equal to the percentage of line expressing the *k13* C580Y mutation or multicopy *pm2/3* at the assay endpoint. P is equal to the percentage of the co-culture that expresses the *k13* WT allele or single-copy *pm2/3* on day 0. n is equal to the number of generations from the assay start to finish, and x is equal to the fitness cost. For the TaqMan allelic discrimination real-time PCR assays described above, n was set to 20.

### CQ or PPQ uptake assays in PfCRT-containing proteoliposomes

Inhibition of CQ or PPQ uptake by MMV665939 and MMV675939 compounds in proteoliposomes containing PfCRT were studied using transport assays performed as described previously (*28*), with the following modifications. Purified PfCRT variants were reconstituted in preformed liposomes made of *E. coli* total lipids:cholesteryl hemisuccinate 97:3 (w/w) and the lumen of the proteoliposomes was composed of 100 mM KP_i_, pH 7.5 and 2 mM β-mercaptoethanol. Uptake of 100 nM [^3^H]-CQ (1 Ci/mmol), or [^3^H]-PPQ (1 Ci/mmol) was performed by diluting PfCRT-containing proteoliposomes (30 ng PfCRT per assay) in 50 μl of 100 mM Tris/MES, pH 5.5 in the absence or presence of 1μM atovaquone, CQ, PPQ, MMV665939 or MMV675939. 1 μM valinomycin was added to the reaction to generate a K^+^ diffusion potential-driven membrane potential. After 30s, the reactions were quenched by the addition of ice-cold 100 mM KP_i_ pH 6.0 and 100 mM LiCl before filtration. The radioactivity retained on the filters was determined with scintillation counting in a Hidex SL300 scintillation counter. Proteoliposomes that contain the CQ-resistant 7G8, or the PPQ-resistant 7G8+F145I or 7G8+C350R *pfcrt* isoforms were used herein. Negative controls of no drug (“-”) and atovaquone were included, where we expected no or minimal inhibition of CQ or PPQ uptake. CQ or PPQ was used as the positive control as PPQ is reported to interact with PfCRT and inhibit binding of CQ. Data were normalized to the specific signal (total counts per minute in PfCRT-containing proteoliposomes minus counts per minute in liposomes devoid of PfCRT). Data means ± SE are presented as percentage of data obtained in the absence of drug (“-”) which were normalized at 100%. Three independent experiments were performed as technical duplicates.

### Statistical Analysis

Unless stated otherwise, experiments were performed on at least four independent occasions, and exact sample sizes are given in the Figure legends or Supplementary Tables and Figures. Scatterplots and bar graphs are presented as means ± SE. Data were analyzed and plotted using Prism v8.3.1 (GraphPad). QTL mapping was analyzed and plotted in R using R/qtl versions 1 and 2 and ggplot2. Comparisons between two groups were analyzed by Mann-Whitney *U* tests if at least four repeats were obtained per group, or unpaired Student’s *t* test if otherwise. Statistical significance was defined by two-sided *P* values (\**P* < 0.05, \*\**P* < 0.01, \*\*\**P* < 0.001). *P* values less than 0.05 were considered statistically significant. ns indicates no significant difference.

## Author contributions

Conceptualization: SM, PS, and DAF. Methodology: SM, TY, MJS, MK, AKT, GM, PS, and DAF. Formal analysis: SM, TY, and DH. Investigation: SM, TY, DH, MJS, LSR, KEW, SKD, JLB, AYB, KJF, EGI, HP, FDR, and JK. Data Curation: SM and TY. Visualization: SM, TY and DH. Supervision: SM, ACU, FM, MQ, PS, and DAF. Writing: SM and DAF. All authors reviewed and approved of the submitted version.

## Data and materials availability

All data are available in the main text and/or the supplementary materials. Raw reads of WGS used in this study have been deposited in NCBI Sequenced Read Archive (SRA) and will be available upon publication of the study.

## Supplementary Materials

Figs. S1 to S13.

Tables S1 to S13.

## References

1. World Health Organization, WHO World Malaria Report 2022. https://www.who.int/teams/global-malaria-programme/reports/world-malaria-report-2022, (2022).

2. C. V. Plowe, Malaria chemoprevention and drug resistance: a review of the literature and policy implications. Malar J 21, 104 (2022).

3. B. Hanboonkunupakarn, N. J. White, Advances and roadblocks in the treatment of malaria. Br J Clin Pharmacol 88, 374–382 (2022).

4. A. M. Dondorp, F. Nosten, P. Yi, D. Das, A. P. Phyo, J. Tarning, K. M. Lwin, F. Ariey, W. Hanpithakpong, S. J. Lee, P. Ringwald, K. Silamut, M. Imwong, K. Chotivanich, P. Lim, T. Herdman, S. S. An, S. Yeung, P. Singhasivanon, N. P. Day, N. Lindegardh, D. Socheat, N. J. White, Artemisinin resistance in *Plasmodium falciparum* malaria. N Engl J Med 361, 455–467 (2009).

5. M. Imwong, M. Dhorda, K. Myo Tun, A. M. Thu, A. P. Phyo, S. Proux, K. Suwannasin, C. Kunasol, S. Srisutham, J. Duanguppama, R. Vongpromek, C. Promnarate, A. Saejeng, N. Khantikul, R. Sugaram, S. Thanapongpichat, N. Sawangjaroen, K. Sutawong, K. T. Han, Y. Htut, K. Linn, A. A. Win, T. M. Hlaing, R. W. van der Pluijm, M. Mayxay, T. Pongvongsa, K. Phommasone, R. Tripura, T. J. Peto, L. von Seidlein, C. Nguon, D. Lek, X. H. S. Chan, H. Rekol, R. Leang, C. Huch, D. P. Kwiatkowski, O. Miotto, E. A. Ashley, M. P. Kyaw, S. Pukrittayakamee, N. P. J. Day, A. M. Dondorp, F. M. Smithuis, F. H. Nosten, N. J. White, Molecular epidemiology of resistance to antimalarial drugs in the Greater Mekong subregion: an observational study. Lancet Infect Dis 20, 1470–1480 (2020).

6. M. Dhorda, C. Amaratunga, A. M. Dondorp, Artemisinin and multidrug-resistant *Plasmodium falciparum* - a threat for malaria control and elimination. Curr Opin Infect Dis 34, 432–439 (2021).

7. D. L. Saunders, P. Vanachayangkul, C. Lon, U.S. Army Military Malaria Research Program; National Center for Parasitology, Entomology, and Malaria Control; Royal Cambodian Armed Forces, Dihydroartemisinin-piperaquine failure in Cambodia. N Engl J Med 371, 484–485 (2014).

8. R. W. van der Pluijm, M. Imwong, N. H. Chau, N. T. Hoa, N. T. Thuy-Nhien, N. V. Thanh, P. Jittamala, B. Hanboonkunupakarn, K. Chutasmit, C. Saelow, R. Runjarern, W. Kaewmok, R. Tripura, T. J. Peto, S. Yok, S. Suon, S. Sreng, S. Mao, S. Oun, S. Yen, C. Amaratunga, D. Lek, R. Huy, M. Dhorda, K. Chotivanich, E. A. Ashley, M. Mukaka, N. Waithira, P. Y. Cheah, R. J. Maude, R. Amato, R. D. Pearson, S. Goncalves, C. G. Jacob, W. L. Hamilton, R. M. Fairhurst, J. Tarning, M. Winterberg, D. P. Kwiatkowski, S. Pukrittayakamee, T. T. Hien, N. P. Day, O. Miotto, N. J. White, A. M. Dondorp, Determinants of dihydroartemisinin-piperaquine treatment failure in *Plasmodium falciparum* malaria in Cambodia, Thailand, and Vietnam: a prospective clinical, pharmacological, and genetic study. Lancet Infect Dis 19, 952–961 (2019).

9. F. Ariey, B. Witkowski, C. Amaratunga, J. Beghain, A. C. Langlois, N. Khim, S. Kim, V. Duru, C. Bouchier, L. Ma, P. Lim, R. Leang, S. Duong, S. Sreng, S. Suon, C. M. Chuor, D. M. Bout, S. Menard, W. O. Rogers, B. Genton, T. Fandeur, O. Miotto, P. Ringwald, J. Le Bras, A. Berry, J. C. Barale, R. M. Fairhurst, F. Benoit-Vical, O. Mercereau-Puijalon, D. Menard, A molecular marker of artemisinin-resistant *Plasmodium falciparum* malaria. Nature 505, 50–55 (2014).

10. R. Amato, R. D. Pearson, J. Almagro-Garcia, C. Amaratunga, P. Lim, S. Suon, S. Sreng, E. Drury, J. Stalker, O. Miotto, R. M. Fairhurst, D. P. Kwiatkowski, Origins of the current outbreak of multidrug-resistant malaria in Southeast Asia: a retrospective genetic study. Lancet Infect Dis 18, 337–345 (2018).

11. F. A. Siddiqui, R. Boonhok, M. Cabrera, H. G. N. Mbenda, M. Wang, H. Min, X. Liang, J. Qin, X. Zhu, J. Miao, Y. Cao, L. Cui, Role of *Plasmodium falciparum* Kelch 13 protein mutations in *P. falciparum* populations from northeastern Myanmar in mediating artemisinin resistance. mBio 11, e01134–19 (2020).

12. B. H. Stokes, S. K. Dhingra, K. Rubiano, S. Mok, J. Straimer, N. F. Gnadig, I. Deni, K. A. Schindler, J. R. Bath, K. E. Ward, J. Striepen, T. Yeo, L. S. Ross, E. Legrand, F. Ariey, C. H. Cunningham, I. M. Souleymane, A. Gansane, R. Nzoumbou-Boko, C. Ndayikunda, A. M. Kabanywanyi, A. Uwimana, S. J. Smith, O. Kolley, M. Ndounga, M. Warsame, R. Leang, F. Nosten, T. J. Anderson, P. J. Rosenthal, D. Menard, D. A. Fidock, *Plasmodium falciparum* K13 mutations in Africa and Asia impact artemisinin resistance and parasite fitness. Elife 10, e66277 (2021).

13. T. Yang, L. M. Yeoh, M. V. Tutor, M. W. Dixon, P. J. McMillan, S. C. Xie, J. L. Bridgford, D. L. Gillett, M. F. Duffy, S. A. Ralph, M. J. McConville, L. Tilley, S. A. Cobbold, Decreased K13 abundance reduces hemoglobin catabolism and proteotoxic stress, underpinning artemisinin resistance. Cell Rep 29, 2917–2928 (2019).

14. J. Birnbaum, S. Scharf, S. Schmidt, E. Jonscher, W. A. M. Hoeijmakers, S. Flemming, C. G. Toenhake, M. Schmitt, R. Sabitzki, B. Bergmann, U. Frohlke, P. Mesen-Ramirez, A. Blancke Soares, H. Herrmann, R. Bartfai, T. Spielmann, A Kelch13-defined endocytosis pathway mediates artemisinin resistance in malaria parasites. Science 367, 51–59 (2020).

15. S. Mok, B. H. Stokes, N. F. Gnadig, L. S. Ross, T. Yeo, C. Amaratunga, E. Allman, L. Solyakov, R. Bottrill, J. Tripathi, R. M. Fairhurst, M. Llinas, Z. Bozdech, A. B. Tobin, D. A. Fidock, Artemisinin-resistant K13 mutations rewire *Plasmodium falciparum*’s intra-erythrocytic metabolic program to enhance survival. Nat Commun 12, 530 (2021).

16. F. A. Siddiqui, X. Liang, L. Cui, *Plasmodium falciparum* resistance to ACTs: Emergence, mechanisms, and outlook. Int J Parasitol Drugs Drug Resist 16, 102–118 (2021).

17. C. O. Egwu, P. Perio, J. M. Augereau, I. Tsamesidis, F. Benoit-Vical, K. Reybier, Resistance to artemisinin in *falciparum* malaria parasites: A redox-mediated phenomenon. Free Radic Biol Med 179, 317–327 (2022).

18. K. E. Ward, D. A. Fidock, J. L. Bridgford, *Plasmodium falciparum* resistance to artemisinin-based combination therapies. Curr Opin Microbiol 69, 102193 (2022).

19. A. Uwimana, N. Umulisa, M. Venkatesan, S. S. Svigel, Z. Zhou, T. Munyaneza, R. M. Habimana, A. Rucogoza, L. F. Moriarty, R. Sandford, E. Piercefield, I. Goldman, B. Ezema, E. Talundzic, M. A. Pacheco, A. A. Escalante, D. Ngamije, J. N. Mangala, M. Kabera, K. Munguti, M. Murindahabi, W. Brieger, C. Musanabaganwa, L. Mutesa, V. Udhayakumar, A. Mbituyumuremyi, E. S. Halsey, N. W. Lucchi, Association of *Plasmodium falciparum* kelch13 R561H genotypes with delayed parasite clearance in Rwanda: an open-label, single-arm, multicentre, therapeutic efficacy study. Lancet Infect Dis 21, 1120–1128 (2021).

20. A. Uwimana, E. Legrand, B. H. Stokes, J. M. Ndikumana, M. Warsame, N. Umulisa, D. Ngamije, T. Munyaneza, J. B. Mazarati, K. Munguti, P. Campagne, A. Criscuolo, F. Ariey, M. Murindahabi, P. Ringwald, D. A. Fidock, A. Mbituyumuremyi, D. Menard, Emergence and clonal expansion of *in vitro* artemisinin-resistant *Plasmodium falciparum* kelch13 R561H mutant parasites in Rwanda. Nat Med 26, 1602–1608 (2020).

21. B. Balikagala, N. Fukuda, M. Ikeda, O. T. Katuro, S. I. Tachibana, M. Yamauchi, W. Opio, S. Emoto, D. A. Anywar, E. Kimura, N. M. Q. Palacpac, E. I. Odongo-Aginya, M. Ogwang, T. Horii, T. Mita, Evidence of artemisinin-resistant malaria in Africa. N Engl J Med 385, 1163–1171 (2021).

22. B. H. Stokes, K. E. Ward, D. A. Fidock, Evidence of artemisinin-resistant malaria in Africa. N Engl J Med 386, 1385–1386 (2022).

23. B. Witkowski, V. Duru, N. Khim, L. S. Ross, B. Saintpierre, J. Beghain, S. Chy, S. Kim, S. Ke, N. Kloeung, R. Eam, C. Khean, M. Ken, K. Loch, A. Bouillon, A. Domergue, L. Ma, C. Bouchier, R. Leang, R. Huy, G. Nuel, J. C. Barale, E. Legrand, P. Ringwald, D. A. Fidock, O. Mercereau-Puijalon, F. Ariey, D. Menard, A surrogate marker of piperaquine-resistant *Plasmodium falciparum* malaria: a phenotype-genotype association study. Lancet Infect Dis 17, 174–183 (2017).

24. R. Amato, P. Lim, O. Miotto, C. Amaratunga, D. Dek, R. D. Pearson, J. Almagro-Garcia, A. T. Neal, S. Sreng, S. Suon, E. Drury, D. Jyothi, J. Stalker, D. P. Kwiatkowski, R. M. Fairhurst, Genetic markers associated with dihydroartemisinin-piperaquine failure in *Plasmodium falciparum* malaria in Cambodia: a genotype-phenotype association study. Lancet Infect Dis 17, 164–173 (2017).

25. V. Duru, N. Khim, R. Leang, S. Kim, A. Domergue, N. Kloeung, S. Ke, S. Chy, R. Eam, C. Khean, K. Loch, M. Ken, D. Lek, J. Beghain, F. Ariey, P. J. Guerin, R. Huy, O. Mercereau-Puijalon, B. Witkowski, D. Menard, *Plasmodium falciparum* dihydroartemisinin-piperaquine failures in Cambodia are associated with mutant K13 parasites presenting high survival rates in novel piperaquine *in vitro* assays: retrospective and prospective investigations. BMC Med 13, 305 (2015).

26. S. Agrawal, K. A. Moser, L. Morton, M. P. Cummings, A. Parihar, A. Dwivedi, A. C. Shetty, E. F. Drabek, C. G. Jacob, P. P. Henrich, C. M. Parobek, K. Jongsakul, R. Huy, M. D. Spring, C. A. Lanteri, S. Chaorattanakawee, C. Lon, M. M. Fukuda, D. L. Saunders, D. A. Fidock, J. T. Lin, J. J. Juliano, C. V. Plowe, J. C. Silva, S. Takala-Harrison, Association of a novel mutation in the *Plasmodium falciparum* chloroquine resistance transporter with decreased piperaquine sensitivity. J Infect Dis 216, 468–476 (2017).

27. L. S. Ross, S. K. Dhingra, S. Mok, T. Yeo, K. J. Wicht, K. Kumpornsin, S. Takala-Harrison, B. Witkowski, R. M. Fairhurst, F. Ariey, D. Menard, D. A. Fidock, Emerging Southeast Asian PfCRT mutations confer *Plasmodium falciparum* resistance to the first-line antimalarial piperaquine. Nat Commun 9, 3314 (2018).

28. J. Kim, Y. Z. Tan, K. J. Wicht, S. K. Erramilli, S. K. Dhingra, J. Okombo, J. Vendome, L. M. Hagenah, S. I. Giacometti, A. L. Warren, K. Nosol, P. D. Roepe, C. S. Potter, B. Carragher, A. Kossiakoff, M. Quick, D. A. Fidock, F. Mancia, Structure and drug resistance of the *Plasmodium falciparum* transporter PfCRT. Nature 576, 315–320 (2019).

29. B. Riegel, P. D. Roepe, Altered drug transport by *Plasmodium falciparum* chloroquine resistance transporter isoforms harboring mutations associated with piperaquine resistance. Biochemistry 59, 2484–2493 (2020).

30. K. J. Wicht, S. Mok, D. A. Fidock, Molecular mechanisms of drug resistance in *Plasmodium falciparum* malaria. Annu Rev Microbiol 74, 431–454 (2020).

31. S. Bopp, P. Magistrado, W. Wong, S. F. Schaffner, A. Mukherjee, P. Lim, M. Dhorda, C. Amaratunga, C. J. Woodrow, E. A. Ashley, N. J. White, A. M. Dondorp, R. M. Fairhurst, F. Ariey, D. Menard, D. F. Wirth, S. K. Volkman, Plasmepsin II-III copy number accounts for bimodal piperaquine resistance among Cambodian *Plasmodium falciparum*. Nat Commun 9, 1769 (2018).

32. A. M. Vaughan, R. S. Pinapati, I. H. Cheeseman, N. Camargo, M. Fishbaugher, L. A. Checkley, S. Nair, C. A. Hutyra, F. H. Nosten, T. J. Anderson, M. T. Ferdig, S. H. Kappe, *Plasmodium falciparum* genetic crosses in a humanized mouse model. Nat Methods 12, 631–633 (2015).

33. K. A. Button-Simons, S. Kumar, N. Carmago, M. T. Haile, C. Jett, L. A. Checkley, S. Y. Kennedy, R. S. Pinapati, D. A. Shoue, M. McDew-White, X. Li, F. H. Nosten, S. H. Kappe, T. J. C. Anderson, J. Romero-Severson, M. T. Ferdig, S. J. Emrich, A. M. Vaughan, I. H. Cheeseman, The power and promise of genetic mapping from *Plasmodium falciparum* crosses utilizing human liver-chimeric mice. Commun Biol 4, 734 (2021).

34. X. Su, M. T. Ferdig, Y. Huang, C. Q. Huynh, A. Liu, J. You, J. C. Wootton, T. E. Wellems, A genetic map and recombination parameters of the human malaria parasite *Plasmodium falciparum*. Science 286, 1351–1353 (1999).

35. A. Miles, Z. Iqbal, P. Vauterin, R. Pearson, S. Campino, M. Theron, K. Gould, D. Mead, E. Drury, J. O’Brien, V. Ruano Rubio, B. MacInnis, J. Mwangi, U. Samarakoon, L. Ranford-Cartwright, M. Ferdig, K. Hayton, X. Z. Su, T. Wellems, J. Rayner, G. McVean, D. Kwiatkowski, Indels, structural variation, and recombination drive genomic diversity in *Plasmodium falciparum*. Genome Res 26, 1288–1299 (2016).

36. MalariaGen, M. M. Abdel Hamid, M. H. Abdelraheem, D. O. Acheampong, A. Ahouidi, M. Ali, J. Almagro-Garcia, A. Amambua-Ngwa, C. Amaratunga, L. Amenga-Etego, B. Andagalu, T. Anderson, V. Andrianaranjaka, I. Aniebo, E. Aninagyei, F. Ansah, P. O. Ansah, T. Apinjoh, P. Arnaldo, E. Ashley, S. Auburn, G. A. Awandare, H. Ba, V. Baraka, A. Barry, P. Bejon, G. I. Bertin, M. F. Boni, S. Borrmann, T. Bousema, M. Bouyou-Akotet, O. Branch, P. C. Bull, H. Cheah, K. Chindavongsa, T. Chookajorn, K. Chotivanich, A. Claessens, D. J. Conway, V. Corredor, E. Courtier, A. Craig, U. D’Alessandro, S. Dama, N. Day, B. Denis, M. Dhorda, M. Diakite, A. Djimde, C. Dolecek, A. Dondorp, S. Doumbia, C. Drakeley, E. Drury, P. Duffy, D. F. Echeverry, T. G. Egwang, S. M. M. Enosse, B. Erko, R. M. Fairhurst, A. Faiz, C. A. Fanello, M. Fleharty, M. Forbes, M. Fukuda, D. Gamboa, A. Ghansah, L. Golassa, S. Goncalves, G. L. A. Harrison, S. A. Healy, J. A. Hendry, A. Hernandez-Koutoucheva, T. T. Hien, C. A. Hill, F. Hombhanje, A. Hott, Y. Htut, M. Hussein, M. Imwong, D. Ishengoma, S. A. Jackson, C. G. Jacob, J. Jeans, K. J. Johnson, C. Kamaliddin, E. Kamau, J. Keatley, T. Kochakarn, D. S. Konate, A. Konate, A. Kone, D. P. Kwiatkowski, M. P. Kyaw, D. Kyle, M. Lawniczak, S. K. Lee, M. Lemnge, P. Lim, C. Lon, K. M. Loua, C. I. Mandara, J. Marfurt, K. Marsh, R. J. Maude, M. Mayxay, O. Maiga-Ascofare, O. Miotto, T. Mita, V. Mobegi, A. O. Mohamed, O. A. Mokuolu, J. Montgomery, C. M. Morang’a, I. Mueller, K. Murie, P. N. Newton, T. Ngo Duc, T. Nguyen, T. N. Nguyen, T. Nguyen Thi Kim, H. Nguyen Van, H. Noedl, F. Nosten, R. Noviyanti, V. N. Ntui, A. Nzila, L. I. Ochola-Oyier, H. Ocholla, A. Oduro, I. Omedo, M. A. Onyamboko, J. B. Ouedraogo, K. Oyebola, W. A. Oyibo, R. Pearson, N. Peshu, A. P. Phyo, C. V. Plowe, R. N. Price, S. Pukrittayakamee, H. H. Quang, M. Randrianarivelojosia, J. C. Rayner, P. Ringwald, A. Rosanas-Urgell, E. Rovira-Vallbona, V. Ruano-Rubio, L. Ruiz, D. Saunders, A. Shayo, P. Siba, V. J. Simpson, M. S. Sissoko, C. Smith, X. Z. Su, C. Sutherland, S. Takala-Harrison, A. Talman, L. Tavul, N. V. Thanh, V. Thathy, A. M. Thu, M. Toure, A. Tshefu, F. Verra, J. Vinetz, T. E. Wellems, J. Wendler, N. J. White, G. Whitton, W. Yavo, R. W. van der Pluijm, Pf7: an open dataset of Plasmodium falciparum genome variation in 20,000 worldwide samples. Wellcome Open Res 8, 22 (2023).

37. Y. Yuthavong, B. Tarnchompoo, T. Vilaivan, P. Chitnumsub, S. Kamchonwongpaisan, S. A. Charman, D. N. McLennan, K. L. White, L. Vivas, E. Bongard, C. Thongphanchang, S. Taweechai, J. Vanichtanankul, R. Rattanajak, U. Arwon, P. Fantauzzi, J. Yuvaniyama, W. N. Charman, D. Matthews, Malarial dihydrofolate reductase as a paradigm for drug development against a resistance-compromised target. Proc Natl Acad Sci USA 109, 16823–16828 (2012).

38. M. Vanaerschot, L. Lucantoni, T. Li, J. M. Combrinck, A. Ruecker, T. R. S. Kumar, K. Rubiano, P. E. Ferreira, G. Siciliano, S. Gulati, P. P. Henrich, C. L. Ng, J. M. Murithi, V. C. Corey, S. Duffy, O. J. Lieberman, M. I. Veiga, R. E. Sinden, P. Alano, M. J. Delves, K. Lee Sim, E. A. Winzeler, T. J. Egan, S. L. Hoffman, V. M. Avery, D. A. Fidock, Hexahydroquinolines are antimalarial candidates with potent blood-stage and transmission-blocking activity. Nat Microbiol 2, 1403–1414 (2017).

39. A. N. Cowell, E. S. Istvan, A. K. Lukens, M. G. Gomez-Lorenzo, M. Vanaerschot, T. Sakata-Kato, E. L. Flannery, P. Magistrado, E. Owen, M. Abraham, G. LaMonte, H. J. Painter, R. M. Williams, V. Franco, M. Linares, I. Arriaga, S. Bopp, V. C. Corey, N. F. Gnadig, O. Coburn-Flynn, C. Reimer, P. Gupta, J. M. Murithi, P. A. Moura, O. Fuchs, E. Sasaki, S. W. Kim, C. H. Teng, L. T. Wang, A. Akidil, S. Adjalley, P. A. Willis, D. Siegel, O. Tanaseichuk, Y. Zhong, Y. Zhou, M. Llinas, S. Ottilie, F. J. Gamo, M. C. S. Lee, D. E. Goldberg, D. A. Fidock, D. F. Wirth, E. A. Winzeler, Mapping the malaria parasite druggable genome by using *in vitro* evolution and chemogenomics. Science 359, 191–199 (2018).

40. J. M. Murithi, I. Deni, C. F. A. Pasaje, J. Okombo, J. L. Bridgford, N. F. Gnadig, R. L. Edwards, T. Yeo, S. Mok, A. Y. Burkhard, O. Coburn-Flynn, E. S. Istvan, T. Sakata-Kato, M. G. Gomez-Lorenzo, A. N. Cowell, K. J. Wicht, C. Le Manach, G. F. Kalantarov, S. Dey, M. Duffey, B. Laleu, A. K. Lukens, S. Ottilie, M. Vanaerschot, I. N. Trakht, F. J. Gamo, D. F. Wirth, D. E. Goldberg, A. R. Odom John, K. Chibale, E. A. Winzeler, J. C. Niles, D. A. Fidock, The *Plasmodium falciparum* ABC transporter ABCI3 confers parasite strain-dependent pleiotropic antimalarial drug resistance. Cell Chem Biol 29, 824–839 e826 (2022).

41. MalariaGEN *Plasmodium falciparum* Community Project, Genomic epidemiology of artemisinin resistant malaria. Elife 5, e08714 (2016).

42. D. A. Fidock, P. J. Rosenthal, Artemisinin resistance in Africa: How urgent is the threat? Med 2, 1287–1288 (2021).

43. N. J. White, Severe malaria. Malar J 21, 284 (2022).

44. Z. Wang, Y. Wang, M. Cabrera, Y. Zhang, B. Gupta, Y. Wu, K. Kemirembe, Y. Hu, X. Liang, Brashear, S. Shrestha, X. Li, J. Miao, X. Sun, Z. Yang, L. Cui, Artemisinin resistance at the China-Myanmar border and association with mutations in the K13 propeller gene. Antimicrob Agents Chemother 59, 6952–6959 (2015).

45. Y. Zhao, Z. Liu, M. T. Soe, L. Wang, T. N. Soe, H. Wei, A. Than, P. L. Aung, Y. Li, X. Zhang, Y. Hu, H. Wei, Y. Zhang, J. Burgess, F. A. Siddiqui, L. Menezes, Q. Wang, M. P. Kyaw, Y. Cao, L. Cui, Genetic variations associated with drug resistance markers in asymptomatic *Plasmodium falciparum* infections in Myanmar. Genes 10, 692 (2019).

46. S. Dahlstrom, P. E. Ferreira, M. I. Veiga, N. Sedighi, L. Wiklund, A. Martensson, A. Farnert, C. Sisowath, L. Osorio, H. Darban, B. Andersson, A. Kaneko, G. Conseil, A. Bjorkman, J. P. Gil, *Plasmodium falciparum* multidrug resistance protein 1 and artemisinin-based combination therapy in Africa. J Infect Dis 200, 1456–1464 (2009).

47. D. K. Raj, J. Mu, H. Jiang, J. Kabat, S. Singh, M. Sullivan, M. P. Fay, T. F. McCutchan, X. Z. Su, Disruption of a *Plasmodium falciparum* multidrug resistance-associated protein (PfMRP) alters its fitness and transport of antimalarial drugs and glutathione. J Biol Chem 284, 7687–7696 (2009).

48. O. Miotto, R. Amato, E. A. Ashley, B. MacInnis, J. Almagro-Garcia, C. Amaratunga, P. Lim, D. Mead, S. O. Oyola, M. Dhorda, M. Imwong, C. Woodrow, M. Manske, J. Stalker, E. Drury, S. Campino, L. Amenga-Etego, T. N. Thanh, H. T. Tran, P. Ringwald, D. Bethell, F. Nosten, A. P. Phyo, S. Pukrittayakamee, K. Chotivanich, C. M. Chuor, C. Nguon, S. Suon, S. Sreng, P. N. Newton, M. Mayxay, M. Khanthavong, B. Hongvanthong, Y. Htut, K. T. Han, M. P. Kyaw, M. A. Faiz, C. I. Fanello, M. Onyamboko, O. A. Mokuolu, C. G. Jacob, S. Takala-Harrison, C. V. Plowe, N. P. Day, A. M. Dondorp, C. C. Spencer, G. McVean, R. M. Fairhurst, N. J. White, D. P. Kwiatkowski, Genetic architecture of artemisinin-resistant *Plasmodium falciparum*. Nat Genet 47, 226–234 (2015).

49. T. Kampoun, S. Srichairatanakool, P. Prommana, P. J. Shaw, J. L. Green, E. Knuepfer, A. A. Holder, C. Uthaipibull, Apicoplast ribosomal protein S10-V127M enhances artemisinin resistance of a Kelch13 transgenic *Plasmodium falciparum*. Malar J 21, 302 (2022).

50. X. Li, S. Kumar, M. McDew-White, M. Haile, I. H. Cheeseman, S. Emrich, K. Button-Simons, F. Nosten, S. H. I. Kappe, M. T. Ferdig, T. J. C. Anderson, A. M. Vaughan, Genetic mapping of fitness determinants across the malaria parasite *Plasmodium falciparum* life cycle. PLoS Genet 15, e1008453 (2019).

51. W. L. Hamilton, R. Amato, R. W. van der Pluijm, C. G. Jacob, H. H. Quang, N. T. Thuy-Nhien, T. T. Hien, B. Hongvanthong, K. Chindavongsa, M. Mayxay, R. Huy, R. Leang, C. Huch, L. Dysoley, C. Amaratunga, S. Suon, R. M. Fairhurst, R. Tripura, T. J. Peto, Y. Sovann, P. Jittamala, B. Hanboonkunupakarn, S. Pukrittayakamee, N. H. Chau, M. Imwong, M. Dhorda, R. Vongpromek, X. H. S. Chan, R. J. Maude, R. D. Pearson, T. Nguyen, K. Rockett, E. Drury, S. Goncalves, N. J. White, N. P. Day, D. P. Kwiatkowski, A. M. Dondorp, O. Miotto, Evolution and expansion of multidrug-resistant malaria in southeast Asia: a genomic epidemiology study. Lancet Infect Dis 19, 943–951 (2019).

52. R. M. Hoglund, L. Workman, M. D. Edstein, N. X. Thanh, N. N. Quang, I. Zongo, J. B. Ouedraogo, S. Borrmann, L. Mwai, C. Nsanzabana, R. N. Price, P. Dahal, N. C. Sambol, S. Parikh, F. Nosten, E. A. Ashley, A. P. Phyo, K. M. Lwin, R. McGready, N. P. Day, P. J. Guerin, N. J. White, K. I. Barnes, J. Tarning, Population pharmacokinetic properties of piperaquine in *falciparum* malaria: an individual participant data meta-Analysis. PLoS Med 14, e1002212 (2017).

53. M. Chugh, V. Sundararaman, S. Kumar, V. S. Reddy, W. A. Siddiqui, K. D. Stuart, P. Malhotra, Protein complex directs hemoglobin-to-hemozoin formation in *Plasmodium falciparum*. Proc Natl Acad Sci USA 110, 5392–5397 (2013).

54. D. Loesbanluechai, N. Kotanan, C. de Cozar, T. Kochakarn, M. R. Ansbro, K. Chotivanich, N. J. White, P. Wilairat, M. C. S. Lee, F. J. Gamo, L. M. Sanz, T. Chookajorn, K. Kumpornsin, Overexpression of plasmepsin II and plasmepsin III does not directly cause reduction in *Plasmodium falciparum* sensitivity to artesunate, chloroquine and piperaquine. Int J Parasitol Drugs Drug Resist 9, 16–22 (2019).

55. S. K. Dhingra, J. L. Small-Saunders, D. Menard, D. A. Fidock, *Plasmodium falciparum* resistance to piperaquine driven by PfCRT. Lancet Infect Dis 19, 1168–1169 (2019).

56. K. J. Wicht, J. L. Small-Saunders, L. M. Hagenah, S. Mok, D. A. Fidock, Mutant PfCRT can mediate piperaquine resistance in African *Plasmodium falciparum* with reduced fitness and increased susceptibility to other antimalarials. J Infect Dis 226, 2021–2029 (2022).

57. S. H. Shafik, S. A. Cobbold, K. Barkat, S. N. Richards, N. S. Lancaster, M. Llinas, S. J. Hogg, R. L. Summers, M. J. McConville, R. E. Martin, The natural function of the malaria parasite’s chloroquine resistance transporter. Nat Commun 11, 3922 (2020).

58. J. Okombo, S. Mok, T. Qahash, T. Yeo, J. Bath, L. M. Orchard, E. Owens, I. Koo, I. Albert, M. Llinas, D. A. Fidock, Piperaquine-resistant PfCRT mutations differentially impact drug transport, hemoglobin catabolism and parasite physiology in *Plasmodium falciparum* asexual blood stages. PLoS Pathog 18, e1010926 (2022).

59. C. P. Sanchez, E. D. T. Manson, S. Moliner Cubel, L. Mandel, S. K. Weidt, M. P. Barrett, M. Lanzer, The knock-down of the chloroquine resistance transporter PfCRT is linked to oligopeptide handling in *Plasmodium falciparum*. Microbiol Spectr 10, e0110122 (2022).

60. M. Imwong, K. Suwannasin, S. Srisutham, R. Vongpromek, C. Promnarate, A. Saejeng, A. P. Phyo, S. Proux, T. Pongvongsa, N. Chea, O. Miotto, R. Tripura, C. Nguyen Hoang, L. Dysoley, N. Ho Dang Trung, T. J. Peto, J. J. Callery, R. W. van der Pluijm, C. Amaratunga, M. Mukaka, L. von Seidlein, M. Mayxay, N. T. Thuy-Nhien, P. N. Newton, N. P. J. Day, E. A. Ashley, F. H. Nosten, F. M. Smithuis, M. Dhorda, N. J. White, A. M. Dondorp, Evolution of multidrug resistance in *Plasmodium falciparum*: a longitudinal study of genetic resistance markers in the Greater Mekong subregion. Antimicrob Agents Chemother 65, e0112121 (2021).

61. M. Venkatesan, N. B. Gadalla, K. Stepniewska, P. Dahal, C. Nsanzabana, C. Moriera, R. N. Price, A. Martensson, P. J. Rosenthal, G. Dorsey, C. J. Sutherland, P. Guerin, T. M. E. Davis, D. Menard, I. Adam, G. Ademowo, C. Arze, F. N. Baliraine, N. Berens-Riha, A. Bjorkman, S. Borrmann, F. Checchi, M. Desai, M. Dhorda, A. A. Djimde, B. B. El-Sayed, T. Eshetu, F. Eyase, C. Falade, J. F. Faucher, G. Froberg, A. Grivoyannis, S. Hamour, S. Houze, J. Johnson, E. Kamugisha, S. Kariuki, J. R. Kiechel, F. Kironde, P. E. Kofoed, J. LeBras, M. Malmberg, L. Mwai, B. Ngasala, F. Nosten, S. L. Nsobya, A. Nzila, M. Oguike, S. D. Otienoburu, B. Ogutu, J. B. Ouedraogo, P. Piola, L. Rombo, B. Schramm, A. F. Some, J. Thwing, J. Ursing, R. P. M. Wong, A. Zeynudin, I. Zongo, C. V. Plowe, C. H. Sibley, G. Asaq Molecular Marker Study, Polymorphisms in *Plasmodium falciparum* chloroquine resistance transporter and multidrug resistance 1 genes: parasite risk factors that affect treatment outcomes for *P. falciparum* malaria after artemether-lumefantrine and artesunate-amodiaquine. Am J Trop Med Hyg 91, 833–843 (2014).

62. P. Tumwebaze, S. Tukwasibwe, A. Taylor, M. Conrad, E. Ruhamyankaka, V. Asua, A. Walakira, J. Nankabirwa, A. Yeka, S. G. Staedke, B. Greenhouse, S. L. Nsobya, M. R. Kamya, G. Dorsey, P. J. Rosenthal, Changing antimalarial drug resistance patterns identified by surveillance at three sites in Uganda. J Infect Dis 215, 631–635 (2017).

63. P. Chotsiri, N. J. White, J. Tarning, Pharmacokinetic considerations in seasonal malaria chemoprevention. Trends Parasitol 38, 673–682 (2022).

64. R. W. van der Pluijm, R. Tripura, R. M. Hoglund, A. Pyae Phyo, D. Lek, A. Ul Islam, A. R. Anvikar, P. Satpathi, S. Satpathi, P. K. Behera, A. Tripura, S. Baidya, M. Onyamboko, N. H. Chau, Y. Sovann, S. Suon, S. Sreng, S. Mao, S. Oun, S. Yen, C. Amaratunga, K. Chutasmit, C. Saelow, R. Runcharern, W. Kaewmok, N. T. Hoa, N. V. Thanh, B. Hanboonkunupakarn, J. J. Callery, A. K. Mohanty, J. Heaton, M. Thant, K. Gantait, T. Ghosh, R. Amato, R. D. Pearson, C. G. Jacob, S. Goncalves, M. Mukaka, N. Waithira, C. J. Woodrow, M. P. Grobusch, M. van Vugt, R. M. Fairhurst, P. Y. Cheah, T. J. Peto, L. von Seidlein, M. Dhorda, R. J. Maude, M. Winterberg, N. T. Thuy-Nhien, D. P. Kwiatkowski, M. Imwong, P. Jittamala, K. Lin, T. M. Hlaing, K. Chotivanich, R. Huy, C. Fanello, E. Ashley, M. Mayxay, P. N. Newton, T. T. Hien, N. Valecha, F. Smithuis, S. Pukrittayakamee, A. Faiz, O. Miotto, J. Tarning, N. P. J. Day, N. J. White, A. M. Dondorp, C. Tracking Resistance to Artemisinin, Triple artemisinin-based combination therapies versus artemisinin-based combination therapies for uncomplicated *Plasmodium falciparum* malaria: a multicentre, open-label, randomised clinical trial. Lancet 395, 1345–1360 (2020).

65. A. Amambua-Ngwa, L. Amenga-Etego, E. Kamau, R. Amato, A. Ghansah, L. Golassa, M. Randrianarivelojosia, D. Ishengoma, T. Apinjoh, O. Maiga-Ascofare, B. Andagalu, W. Yavo, M. Bouyou-Akotet, O. Kolapo, K. Mane, A. Worwui, D. Jeffries, V. Simpson, U. D’Alessandro, D. Kwiatkowski, A. A. Djimde, Major subpopulations of *Plasmodium falciparum* in sub-Saharan Africa. Science 365, 813–816 (2019).

66. A. K. Tripathi, G. Mlambo, S. Kanatani, P. Sinnis, G. Dimopoulos, *Plasmodium falciparum* gametocyte culture and mosquito infection through artificial membrane feeding. J Vis Exp, 10.3791/61426 (2020).

67. R. Xi, A. G. Hadjipanayis, L. J. Luquette, T. M. Kim, E. Lee, J. Zhang, M. D. Johnson, D. M. Muzny, D. A. Wheeler, R. A. Gibbs, R. Kucherlapati, P. J. Park, Copy number variation detection in whole-genome sequencing data using the Bayesian information criterion. Proc Natl Acad Sci USA 108, E1128–1136 (2011).

68. M. Kanai, T. Yeo, V. Asua, P. J. Rosenthal, D. A. Fidock, S. Mok, Comparative analysis of *Plasmodium falciparum* genotyping via SNP detection, microsatellite profiling, and whole-genome sequencing. Antimicrob Agents Chemother 66, e0116321 (2022).

69. M. Imwong, K. Suwannasin, C. Kunasol, K. Sutawong, M. Mayxay, H. Rekol, F. M. Smithuis, T. M. Hlaing, K. M. Tun, R. W. van der Pluijm, R. Tripura, O. Miotto, D. Menard, M. Dhorda, N. P. J. Day, N. J. White, A. M. Dondorp, The spread of artemisinin-resistant *Plasmodium falciparum* in the Greater Mekong subregion: a molecular epidemiology observational study. Lancet Infect Dis 17, 491–497 (2017).

70. B. Witkowski, C. Amaratunga, N. Khim, S. Sreng, P. Chim, S. Kim, P. Lim, S. Mao, C. Sopha, B. Sam, J. M. Anderson, S. Duong, C. M. Chuor, W. R. Taylor, S. Suon, O. Mercereau-Puijalon, R. M. Fairhurst, D. Menard, Novel phenotypic assays for the detection of artemisinin-resistant *Plasmodium falciparum* malaria in Cambodia: *in-vitro* and *ex-vivo* drug-response studies. Lancet Infect Dis 13, 1043–1049 (2013).

71. C. Amaratunga, A. T. Neal, R. M. Fairhurst, Flow cytometry-based analysis of artemisinin-resistant *Plasmodium falciparum* in the ring-stage survival assay. Antimicrob Agents Chemother 58, 4938–4940 (2014).

72. S. J. Gabryszewski, C. Modchang, L. Musset, T. Chookajorn, D. A. Fidock, Combinatorial genetic modeling of *pfcrt*-mediated drug resistance evolution in *Plasmodium falciparum*. Mol Biol Evol 33, 1554–1570 (2016).

