## Supplementary Figures for "Mapping the genomic landscape of multidrug resistance in *Plasmodium falciparum* and its impact on parasite fitness"

**This PDF file includes:**

Figs. S1 to S13
Legends for Tables S1 to S13

**Other Supplementary Materials for this manuscript include the following:**

Tables S1 to S13 (in a separate .xlsx file)

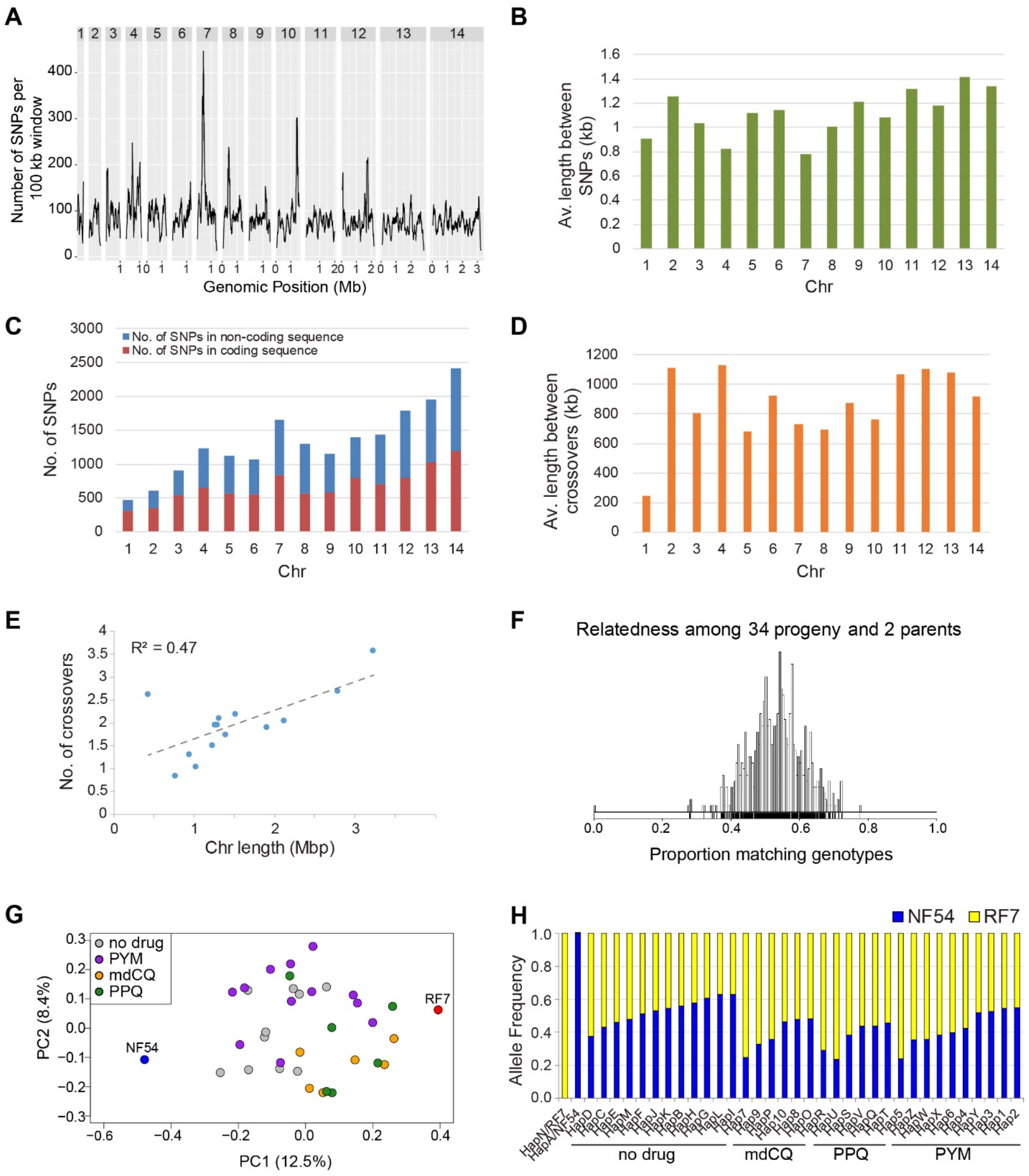
A
Number of SNPs per 100 kb window
Genomic Position (Mb)
B
Av. length between SNPs (kb)
Chr
C
No. of SNPs in non-coding sequence
No. of SNPs in coding sequence
No. of SNPs
Chr
D
Av. length between crossovers (kb)
Chr
E
R² = 0.47
No. of crossovers
Chr length (Mbp)
F
Relatedness among 34 progeny and 2 parents
Proportion matching genotypes
G
no drug
PYM
mdCQ
PPQ
NF54
RF7
PC2 (8.4%)
PC1 (12.5%)
H
NF54
RF7
Allele Frequency
no drug
mdCQ
PPQ
PYM

**Fig. S1. Genomic distribution of SNP density and characteristics for the 18,489 genome-wide SNPs that differ between RF7 and NF54 parents in the core genome and the analysis of recombination events and allele frequencies in the recombinant progeny.** (**A**) No. of SNPs per 100 kb windows plotted based on genomic location. (**B**) Average physical length (in Kb) between SNPs per chromosome. (**C**) No. of coding vs. non-coding SNPs per chromosome. (**D**) Average physical length in kb between crossovers per chromosome for the core genome. Data is averaged across the 34 recombinant progeny. The average physical distance between meiotic crossover events was similar between chromosomes, with the exception of chromosome 1 that showed a relatively high proportion of coding sequence SNPs and more frequent crossovers. (**E**) No. of crossovers vs. chromosome length averaged across the 34 recombinant progeny. (**F**) Genetic relatedness between the 34 recombinant progeny clones and two parents measured by identity-by-descent in multiple pairwise comparisons. (**G**) Principal coordinate analysis of progeny and cross parents based on genetic dissimilarities calculated from the pairwise distance matrix of the 15,869 SNPs (after filtering for markers present in >46/51 progeny). (**H**) Averaged RF7 and NF54 allele frequencies across the genome for each of the 34 recombinant progeny, ordered by drug-treatment condition. PYM, pyrimethamine; mdCQ, monodesethyl chloroquine; PPQ, piperaquine.

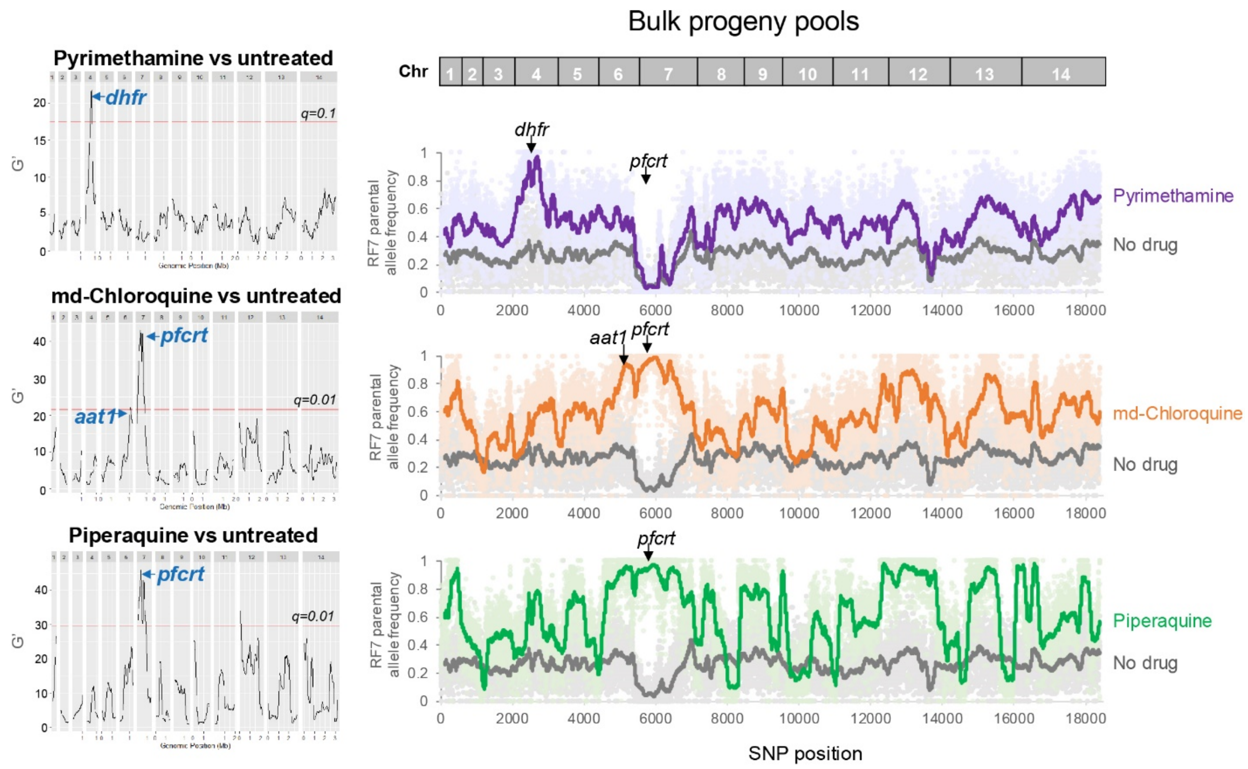


**Fig. S2. Bulk segregant analyses of bulk progeny pools comparing each drug-treatment condition vs. the untreated control.** Shown are the genetic loci enriched by each drug-treated condition and the RF7 allele frequency of bulk progeny pools that were exposed to pyrimethamine, monodesethyl (md-) chloroquine or piperaquine compared to untreated controls in multiple pairwise comparisons. Regions with a False Discovery Rate (FDR) *q*-value < 0.1 (pyrimethamine), 0.01 (md-chloroquine), or 0.01 (piperaquine) were considered statistically significant QTLs. Highlighted are the genes of interest within these QTL segments.

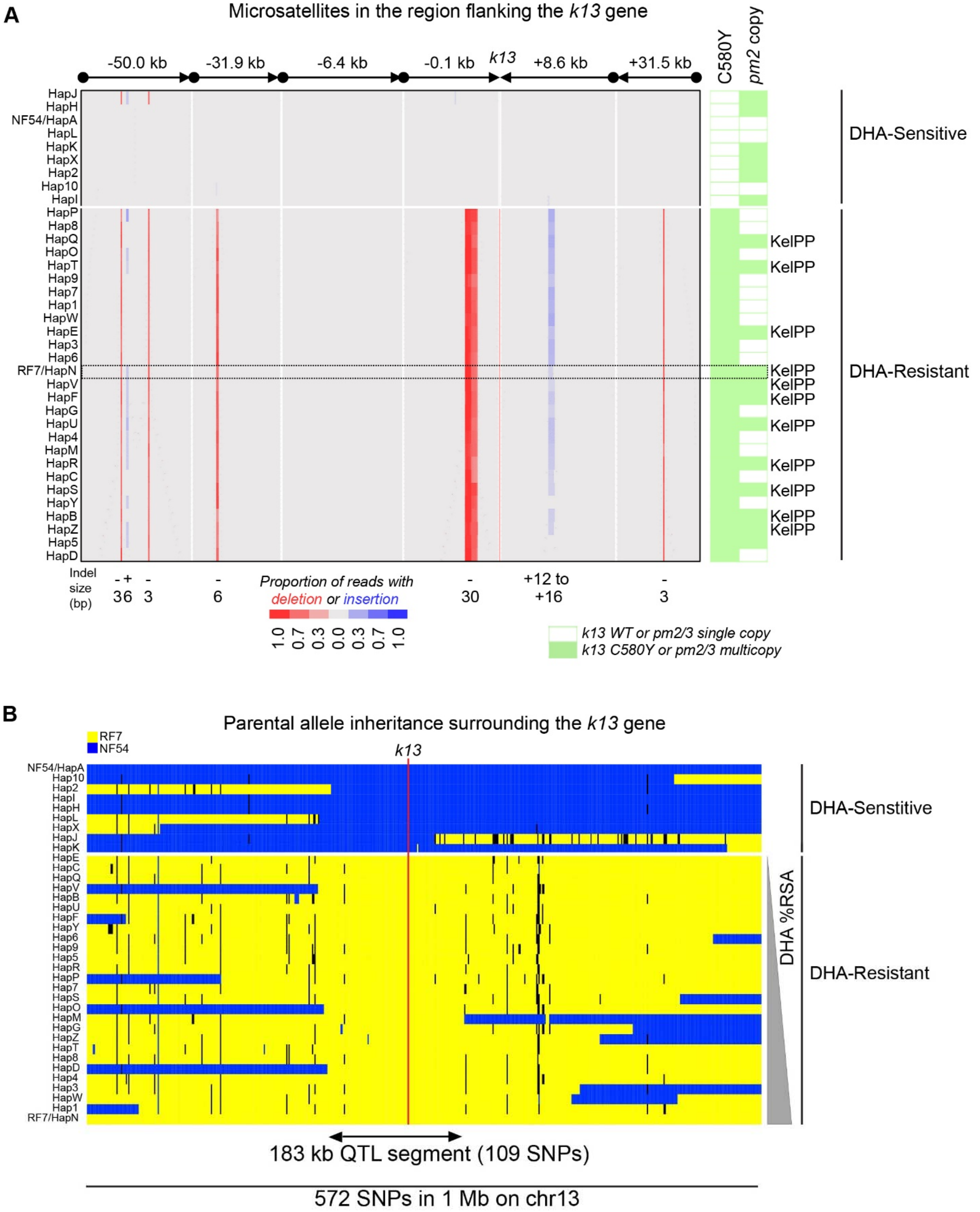
A
Microsatellites in the region flanking the k13 gene
-50.0 kb
-31.9 kb
-6.4 kb
-0.1 kb
k13
+8.6 kb
+31.5 kb
C580Y
pm2 copy
HapJ
HapH
NF54/HapA
HapL
HapK
HapX
Hap2
Hap10
HapI
HapP
Hap8
HapQ
HapO
HapT
Hap9
Hap7
Hap1
HapW
HapE
Hap3
Hap6
RF7/HapN
HapV
HapF
HapG
HapU
Hap4
HapM
HapR
HapC
HapS
HapY
HapB
HapZ
Hap5
HapD
DHA-Sensitive
DHA-Resistant
KelPP
Indel size (bp)
- + 36
- 3
- 6
- 30
+12 to +16
- 3
Proportion of reads with deletion or insertion
1.0 0.7 0.3 0.0 0.3 0.7 1.0
k13 WT or pm2/3 single copy
k13 C580Y or pm2/3 multicopy
B
Parental allele inheritance surrounding the k13 gene
RF7
NF54
k13
NF54/HapA
Hap10
Hap2
HapI
HapH
HapL
HapX
HapJ
HapK
HapE
HapC
HapQ
HapV
HapB
HapU
HapF
HapY
Hap6
Hap9
Hap5
HapR
HapP
Hap7
HapS
HapO
HapM
HapG
HapZ
HapT
Hap8
HapD
Hap4
Hap3
HapW
Hap1
RF7/HapN
DHA-Senstitive
DHA-Resistant
DHA %RSA
183 kb QTL segment (109 SNPs)
572 SNPs in 1 Mb on chr13

**Fig. S3. Microsatellite marker and SNP analysis of parental allele inheritance surrounding *k13* gene among the recombinant progeny.** (**A**) Identification of genetic cross progeny clones harboring the KEL1/PLA1/PfPailin (KelPP) co-lineage by the *k13* genotype, *pm2* copy number determined by WGS and qPCR, and microsatellite size analysis of the -150 kb to +30 kb region surrounding *k13* gene. We defined progeny as KelPP if they fulfilled these three criteria. Shown are the sizes of the insertions (red) and deletions (blue) occurring in the microsatellites flanking *k13* gene for each of the 34 recombinant progeny and the RF7 x NF54 parents. The colors reflect the proportion of WGS reads that contain the event. The RF7 parent is used as a reference as it has the MS signature unique to the Cambodian KEL1/PLA1/PfPailin parasites. (**B**) Allelic map showing the inheritance of either RF7 or NF54 parental alleles for the 572 SNPs located ± 500 kb from *k13* gene in the chr13 segment among the 34 recombinant progeny. The progeny are grouped by DHA susceptibility (RSA > 1% for DHA resistance) and then ordered by the % RSA levels.

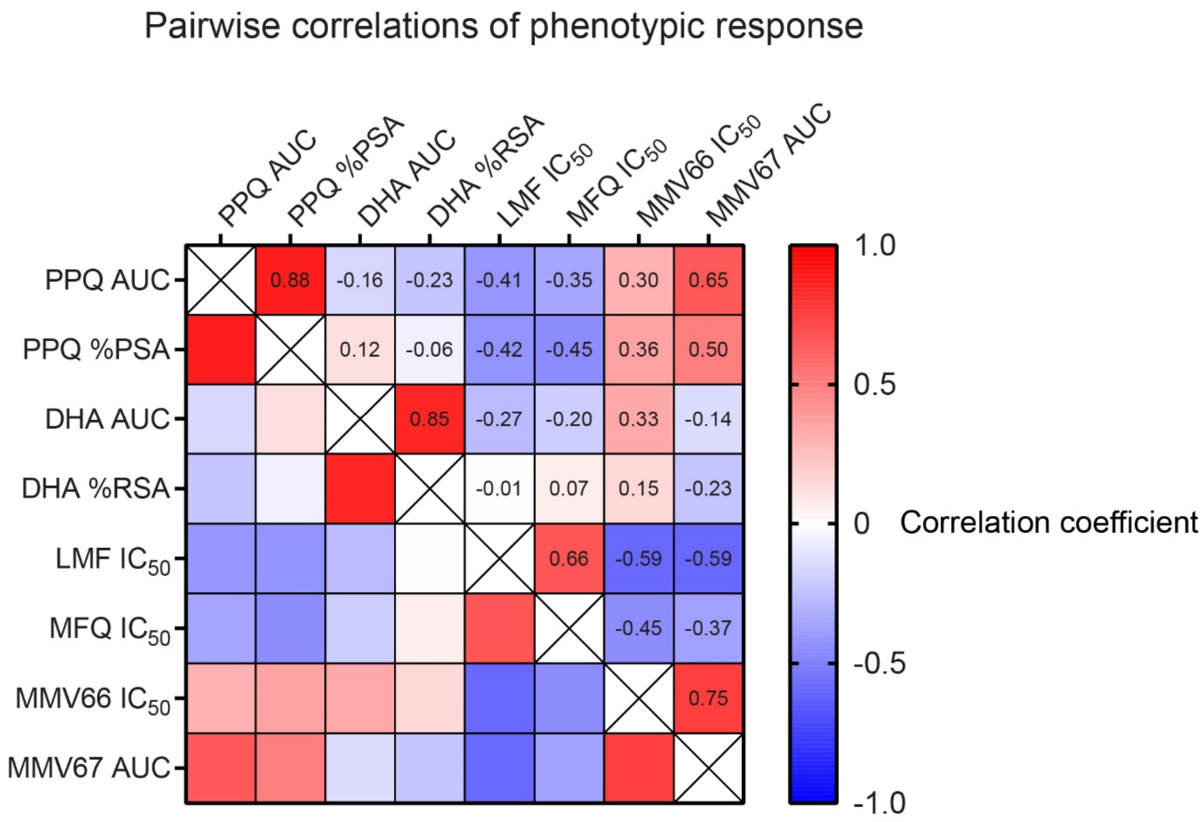


**Fig. S4. Pairwise correlation coefficients for each phenotypic response measured in the progeny clones.** Heatmap of the pairwise correlation coefficients, R, calculated between each of the phenotypic drug responses studied herein in the genetic cross progeny (N=49).

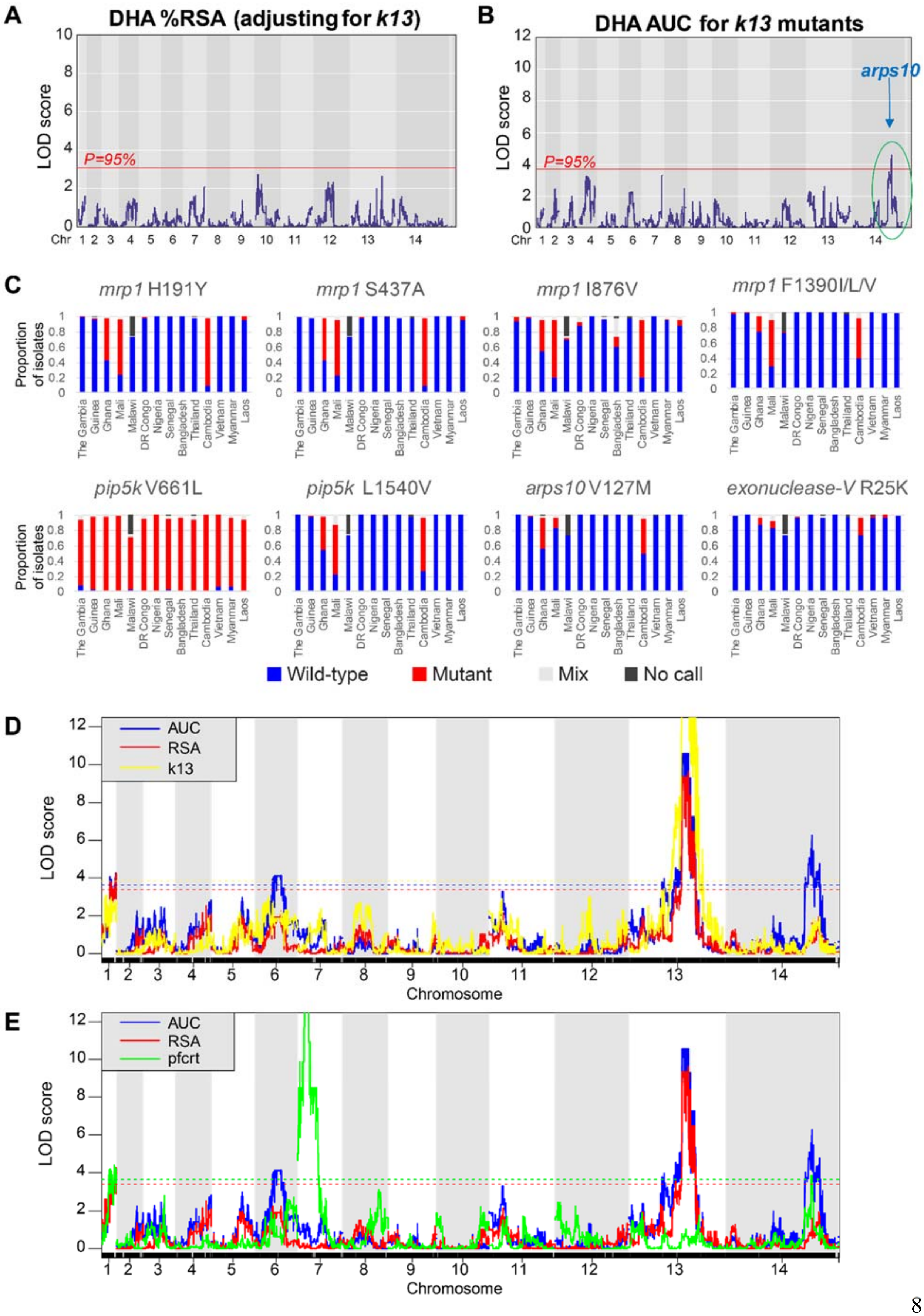

A
DHA %RSA (adjusting for k13)
LOD score
P=95%
Chr
B
DHA AUC for k13 mutants
arps10
C
mrp1 H191Y
mrp1 S437A
mrp1 I876V
mrp1 F1390I/L/V
pip5k V661L
pip5k L1540V
arps10 V127M
exonuclease-V R25K
Proportion of isolates
The Gambia
Guinea
Ghana
Mali
Malawi
DR Congo
Nigeria
Senegal
Bangladesh
Thailand
Cambodia
Vietnam
Myanmar
Laos
Wild-type
Mutant
Mix
No call
D
AUC
RSA
k13
Chromosome
E
AUC
RSA
pfcrt
Chromosome

**Fig. S5. QTLs associated with *k13* and *pfcrt* genotypes and the prevalence of mutations in genes located in chr1 and chr14 segments among clinical isolates.** (**A**) LOD plot for % RSA after adjusting for *k13* as a covariate suggests co-inheritance of chr1 segment with *k13*. (**B**) LOD plot for AUC levels in the subset of *k13* mutants (N=37 progeny) suggests an epistatic interaction of chr14 segment with *k13*. (**C**) Frequency of mutations in clinical samples (Pf3k) of genes in chr1 and chr14 segments: *mrp1, pip5k, arps10, alpha-beta hydrolase.* (**D** and **E**) LOD plots of *k13* (**D**) and *pfcrt* (**E**) genotypes showing linkage of chr1 and chr14 segments with *pfcrt* but not with *k13* using these genotypes as an outcome in the QTL analysis. Shown are the significant QTLs above the 95% probability threshold (dotted lines) for each analysis. RSA: red line; blue line: AUC; yellow line: *k13*; green line: *pfcrt.*

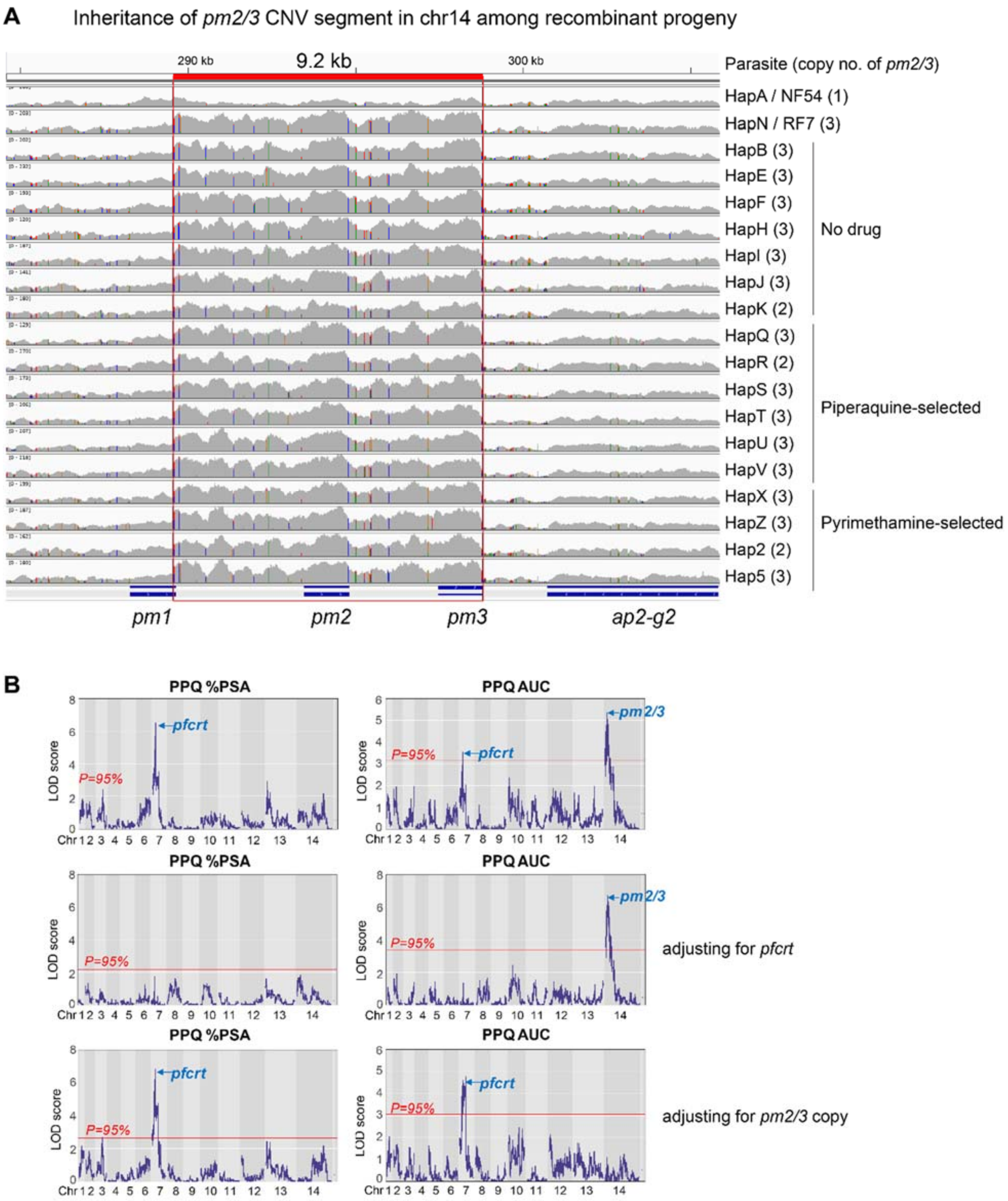
A
Inheritance of pm2/3 CNV segment in chr14 among recombinant progeny
290 kb
9.2 kb
300 kb
Parasite (copy no. of pm2/3)
HapA / NF54 (1)
HapN / RF7 (3)
HapB (3)
HapE (3)
HapF (3)
HapH (3)
HapI (3)
HapJ (3)
HapK (2)
No drug
HapQ (3)
HapR (2)
HapS (3)
HapT (3)
HapU (3)
HapV (3)
Piperaquine-selected
HapX (3)
HapZ (3)
Hap2 (2)
Hap5 (3)
Pyrimethamine-selected
pm1
pm2
pm3
ap2-g2
B
PPQ %PSA
PPQ AUC
LOD score
P=95%
pfcrt
pm2/3
Chr 1 2 3 4 5 6 7 8 9 10 11 12 13 14
adjusting for pfcrt
adjusting for pm2/3 copy

**Fig. S6. The chr14 segment showing inheritance of *pm2/3* amplicon in progeny and QTLs mapped to PPQ response after controlling for *pfcrt* or *pm2/3*.** (**A**) WGS read counts showing the tight conservation of a 9,232 bp segment spanning positions 289,567 to 298,799 on chr14 that contained the *pm2/3* amplicon among recombinant progeny. (**B**) LOD plots showing the QTLs above the 95% probability threshold identified for PPQ after adjusting for *pfcrt* or *pm2/3* as covariates.

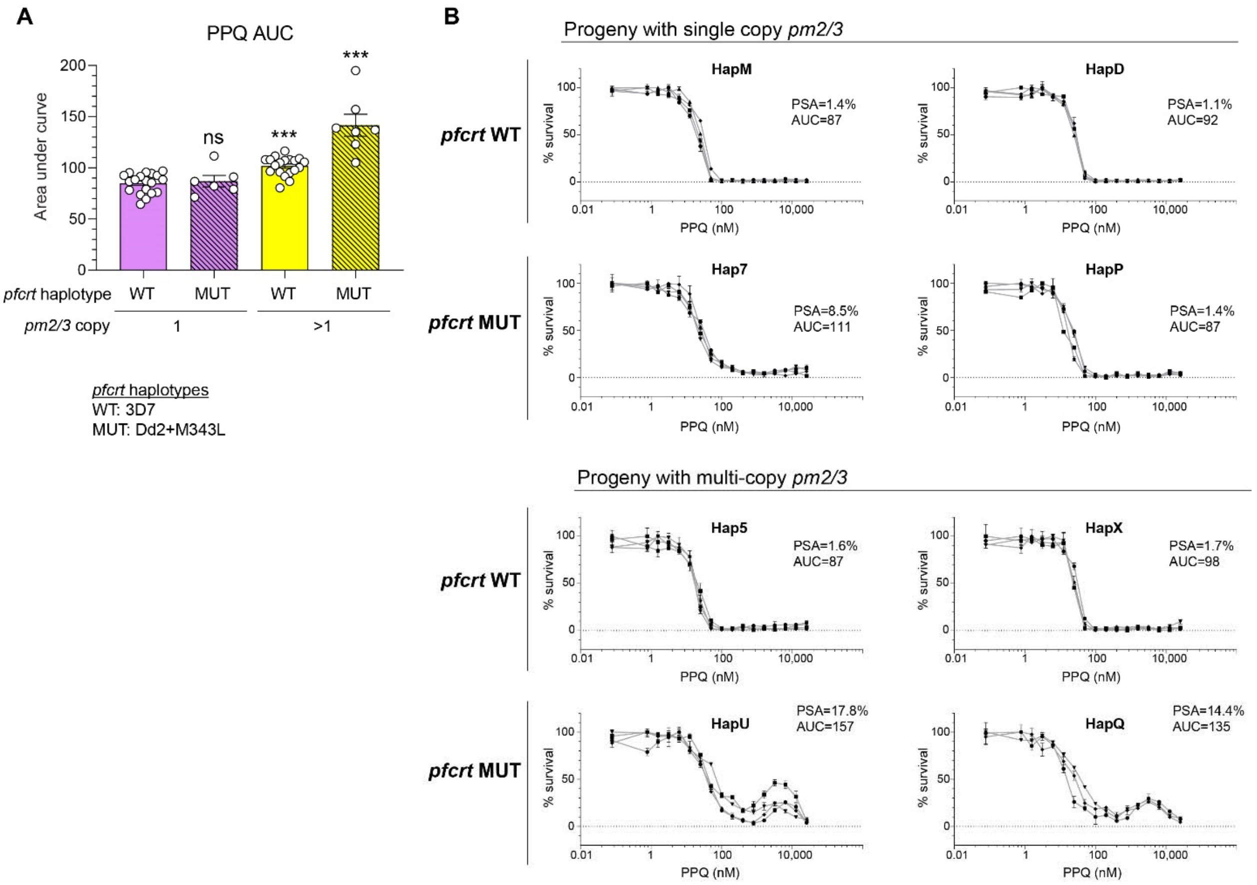


**Fig. S7. PPQ AUC levels and dose response curves for representative recombinant progeny.** (**A**) AUC levels for recombinant progeny segregated by their *pm2/3* copy number and *pfcrt* genotypes. Significance in AUC levels between progeny belonging to each group vs. the group with WT *pfcrt* and single *pm2/3* copy was tested by Mann-Whitney U (N=6-19). ***$P$<0.001. (**B**) PPQ dose response curves for representative recombinant progeny grouped by their *pm2/3* copy number or *pfcrt* genotypes, showing that the biphasic PPQ curve associates with the inheritance of mutant Dd2+M343L *pfcrt* and multicopy *pm2/3*. Each line represents the mean % parasite survival ± SE for one independent experiment performed in technical duplicates. N,n=3-4,2.

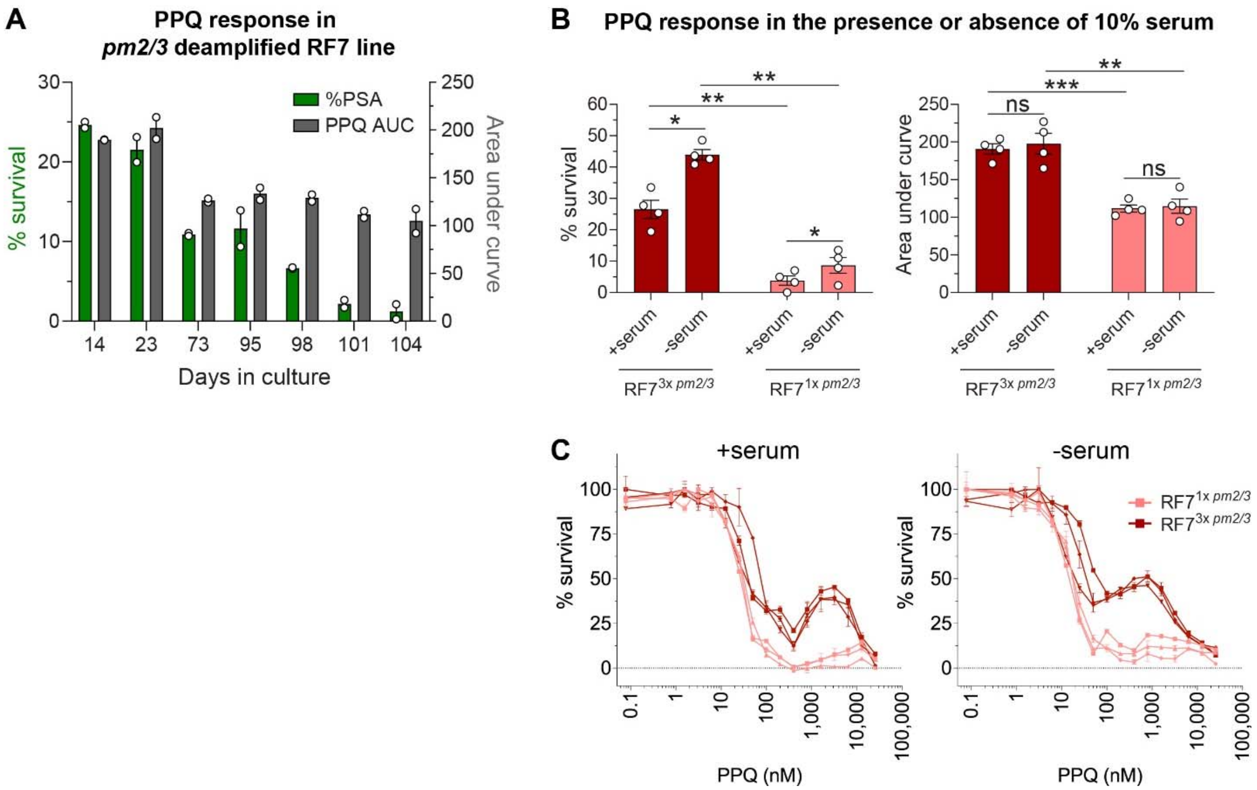


**Fig. S8. Genetic and non-genetic factors affecting PPQ response - *pm2/3* copy and serum levels.** (**A**) Decline in PPQ % PSA and AUC levels accompanied with the loss of *pm2/3* copy during long-term *in vitro* culture adaptation of RF7 line in the absence of PPQ pressure. (**B**) PPQ response in the presence or absence of 10% human serum showing that culture media supplemented with serum significantly reduces the % PSA by ~2-fold in both PPQ-resistant and PPQ-sensitive lines but does not affect AUC levels. Significance in % PSA or AUC levels between the +serum vs. no serum conditions or between the 3 vs. 1 copy *pm2/3* RF7 clones were tested by Student's paired t-test (N,n=4,2). **P*<0.05. ***P*<0.01, ****P*<0.001. (**C**) PPQ dose response curve in RF7 clones showing that serum can affect the PPQ biphasic response. Each line represents the mean % parasite survival ± SE for one independent experiment performed in technical duplicates. N,n=3,2. Our observation suggests that serum components can potentiate PPQ activity.

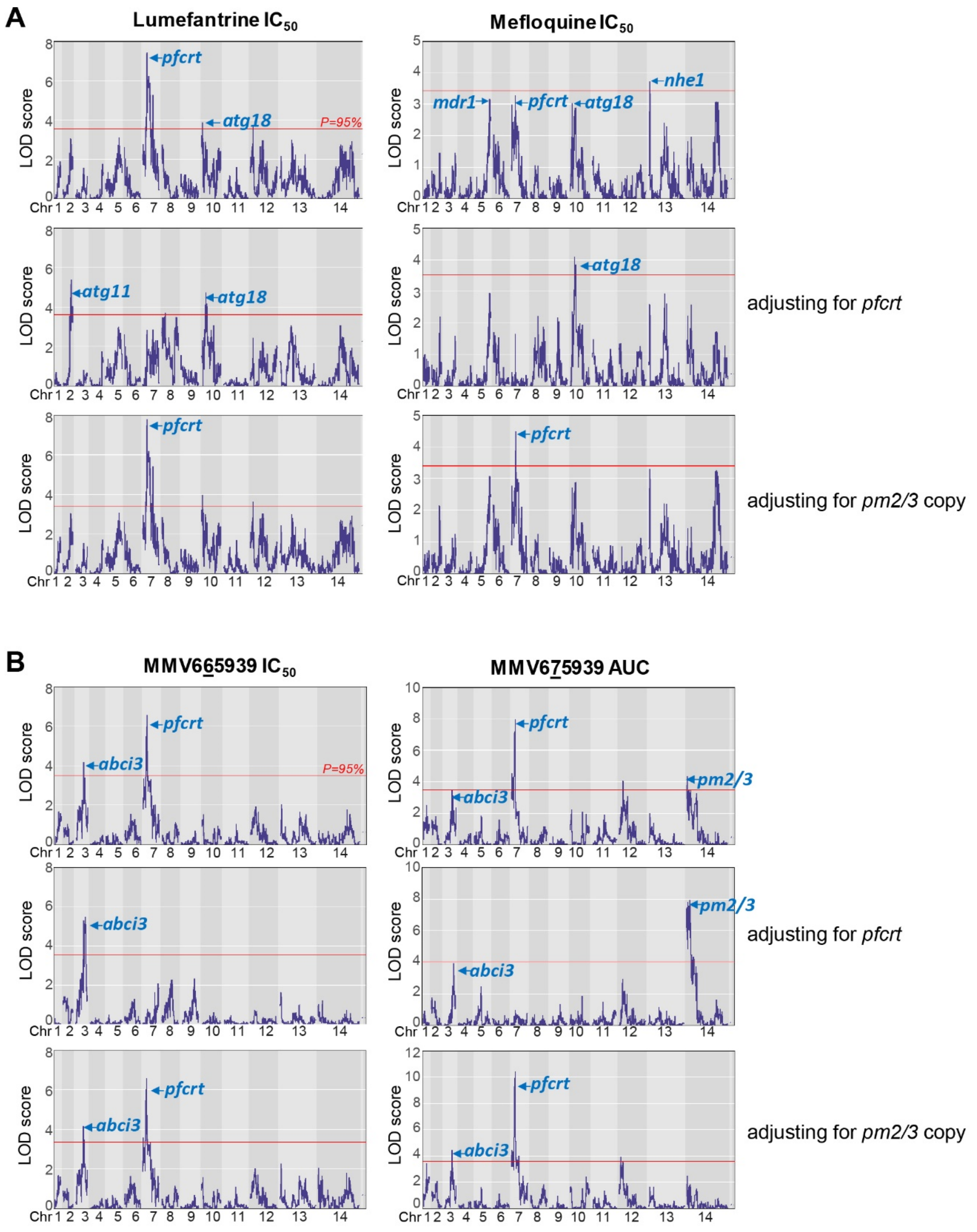


**Fig. S9. QTLs that associate with parasite susceptibility or resistance to antimalarial compounds after controlling for *pfcrt* or *pm2/3*.** (**A** and **B**) LOD score plots for LMF and MFQ (**A**), and MMV665939 and MMV675939 (**B**) after adjusting for *pfcrt* or *pm2/3* as covariates. Shown are the significant QTLs above the 95% probability threshold (red line) for each analysis.

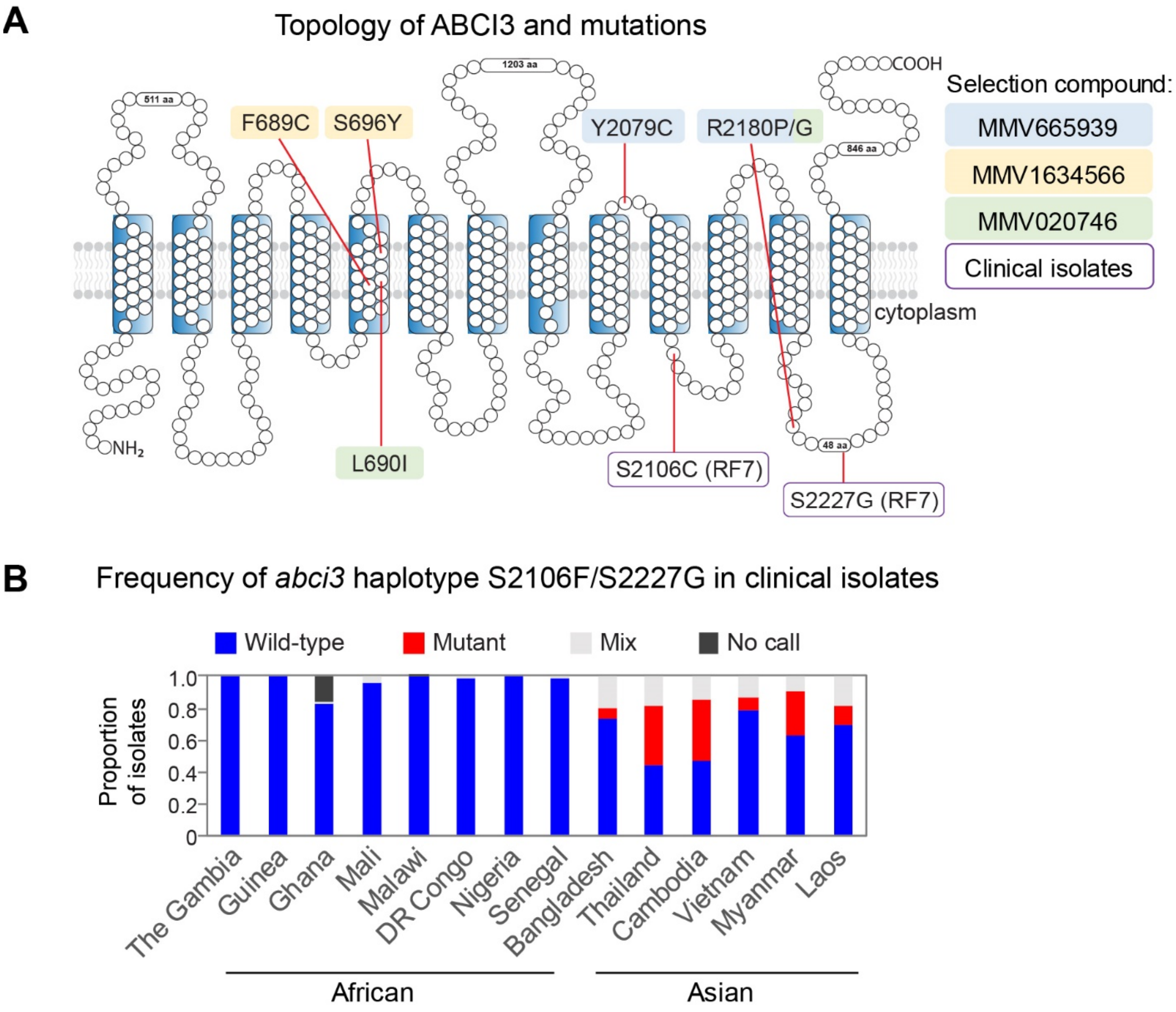


**Fig. S10. Topology of ABCI3 protein and the prevalence of ABCI3 mutations in field isolates.** (**A**) Topology of ABCI3 protein showing the locations of point mutations that arose either via *in vitro* resistance selections using MMV665939 and other preclinical compounds or polymorphisms that naturally occur in field isolates. Not shown are the mutations L79F, F2010L and H2181D, which arose during *in vitro* resistance selections with DDD01034957 but were not validated using gene editing. (**B**) Prevalence of the most frequently occurring ABCI3 haplotype S2106C/S2227G in the Pf3K dataset (2,512 isolates), showing its abundance in Southeast Asian parasites.

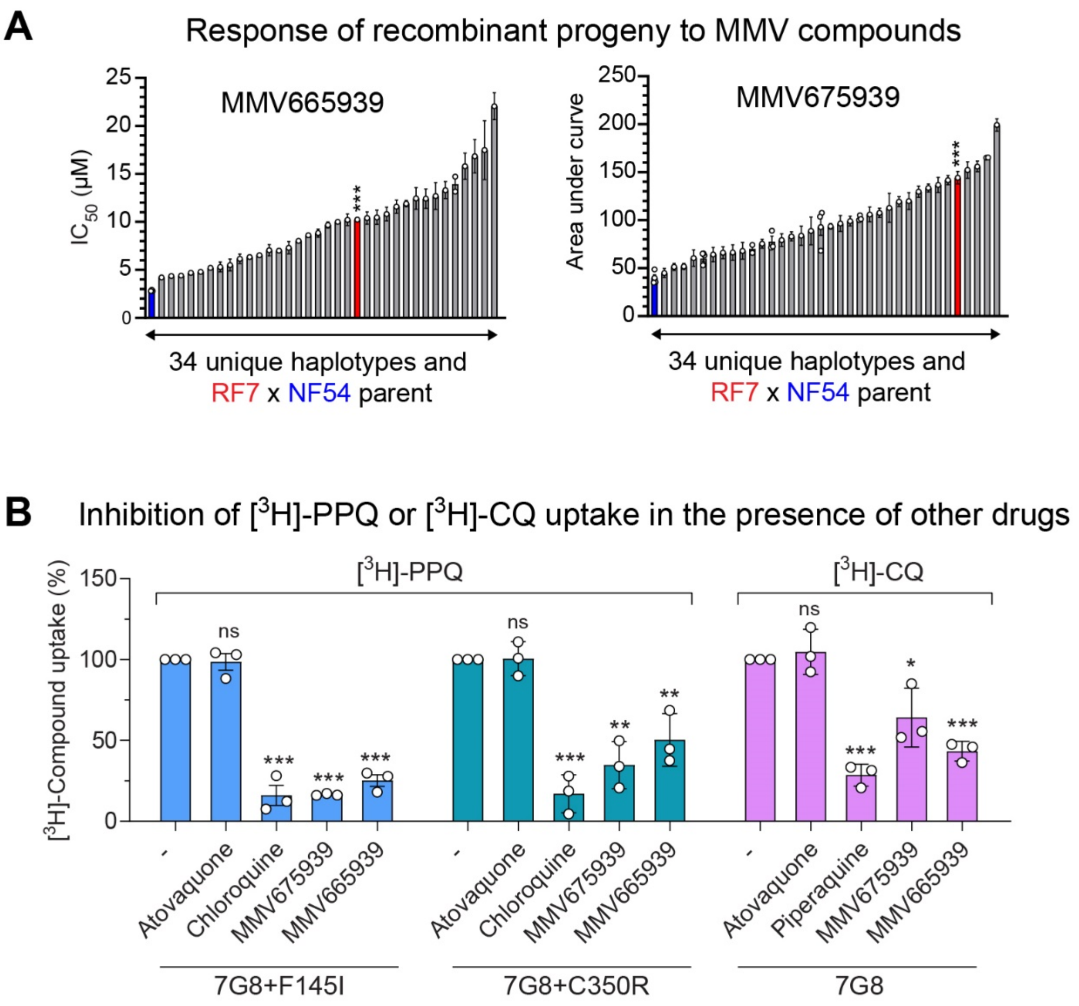


**Fig. S11. MMV6659369 and MMV675939 response in progeny and interaction of these compounds with *pfcrt* using drug uptake assays.** (**A**) Sorted bar plots for MMV6675939 $IC_{50}$ or MMV675939 Area Under the Curve (AUC) values for the progeny clones. Each bar represents the mean $IC_{50}$ or AUC ± SE for a recombinant haplotype. Data are ordered by the resistance levels separately for each drug index. For certain haplotype groups in which identical clones were obtained, multiple points depict more than one sibling progeny being phenotyped. Significance between the genetic cross parents' responses were tested by Mann-Whitney U (N,n=3-14,2). (**B**) Inhibition of 100 nM [$^3$H]-CQ or [$^3$H]-PPQ uptake in the presence of 1 µM atovaquone, chloroquine, piperaquine, MMV665939 or MMV675939, measured in proteoliposomes that contain the CQ-resistant 7G8, or the PPQ-resistant 7G8+F145I or 7G8+C350R *pfcrt* isoforms. Data shown are the means ± SE for three independent experiments performed in technical duplicates. Values were normalized to the signal in proteoliposomes devoid of PfCRT. Negative controls of no drug ("-") and atovaquone were included, where we expected no inhibition of [$^3$H]-CQ or [$^3$H]-PPQ uptake. CQ or PPQ were used as the positive control as PPQ has earlier been shown to interact with PfCRT and inhibit the binding of CQ. Significance in uptake between each drug treatment vs. the no drug ("-") condition was tested using unpaired Student's t-tests. *$P$<0.05, **$P$<0.01, ***$P$<0.001.

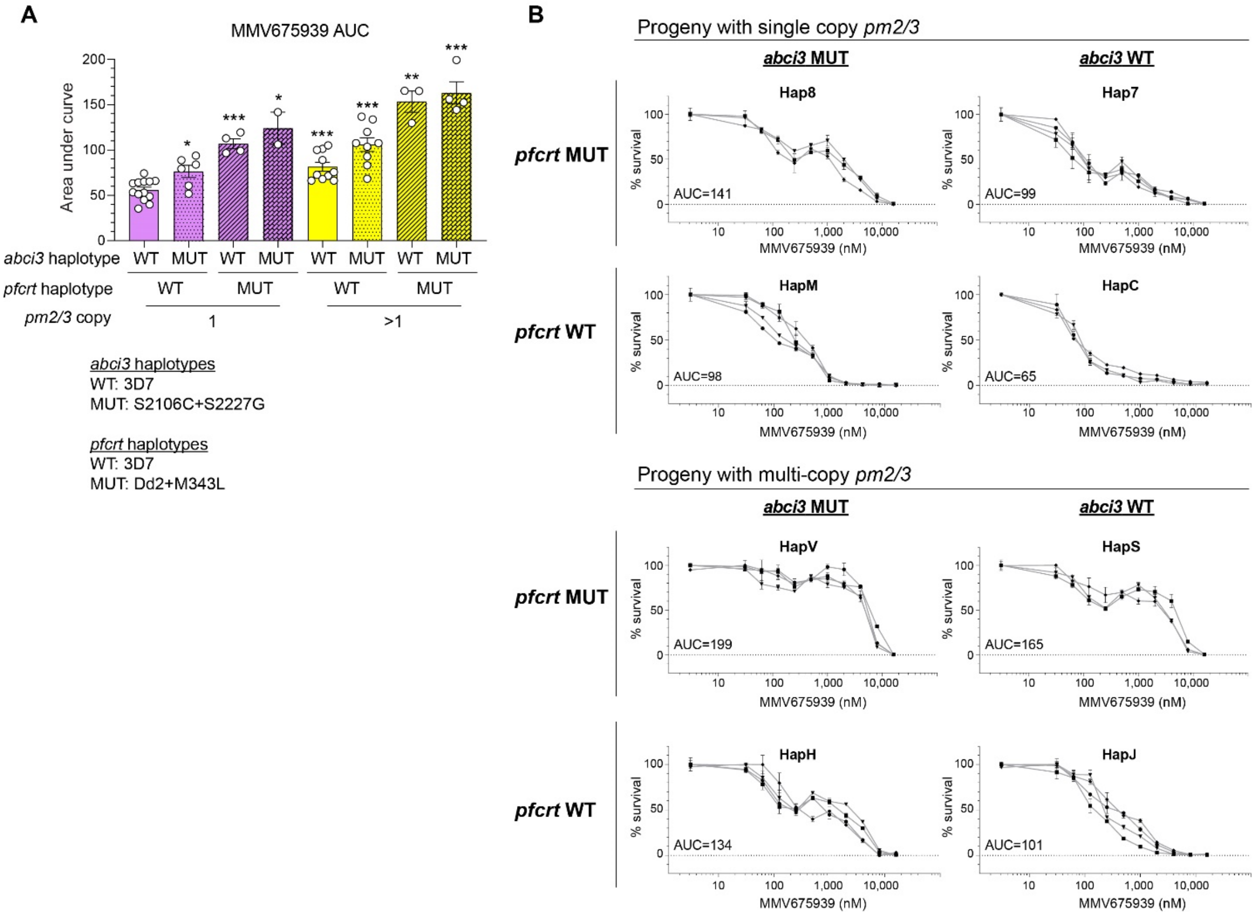


**Fig. S12. MMV675939 AUC levels and dose response curves for representative recombinant progeny.** (**A**) AUC values for recombinant progeny segregated by their *pm2/3* copy number*, pfcrt* and *abci3* genotypes. Significance in AUC levels between progeny belonging to each group vs. the group with WT *abci3* and *pfcrt*, and single *pm2/3* copy was tested by Mann-Whitney U (N=2-13). **P*<0.05, ***P*<0.01, ****P*<0.001. (**B**) MMV675939 dose response curves for representative recombinant progeny grouped by their *pm2/3* copy number*, pfcrt* and *abci3* genotypes showing that either of these three markers can confer a biphasic response. Each plotted line represents the mean % parasite survival ± SE for one independent experiment performed in technical duplicates. Parasites from each haplotype were tested on 3-4 separate occasions, as shown.

| | GrpA | | | GrpB | | |
|---|---|---|---|---|---|---|
| Microsatellite marker | C2M18 | TAA81 | BM5 | C3M67 | TA1 | C13M87 |
| Dye | FAM (blue) | VIC (green) | NED (yellow) | FAM (blue) | VIC (green) | NED (yellow) |
| Size of RF7 (bp) | 126 | 130 | 148 | 128 | 167 | 119 |
| Size of NF54 (bp) | 161 | 121 | 140 | 150 | 185 | 141 |
| Size diff. of RF7 vs. NF54 (bp) | -35 | 9 | 8 | 22 | 18 | 22 |
| Chr | 2 | 5 | 8 | 3 | 6 | 13 |
| Position | 206 | 1214 | 1225 | 800 | 900 | 1828 |

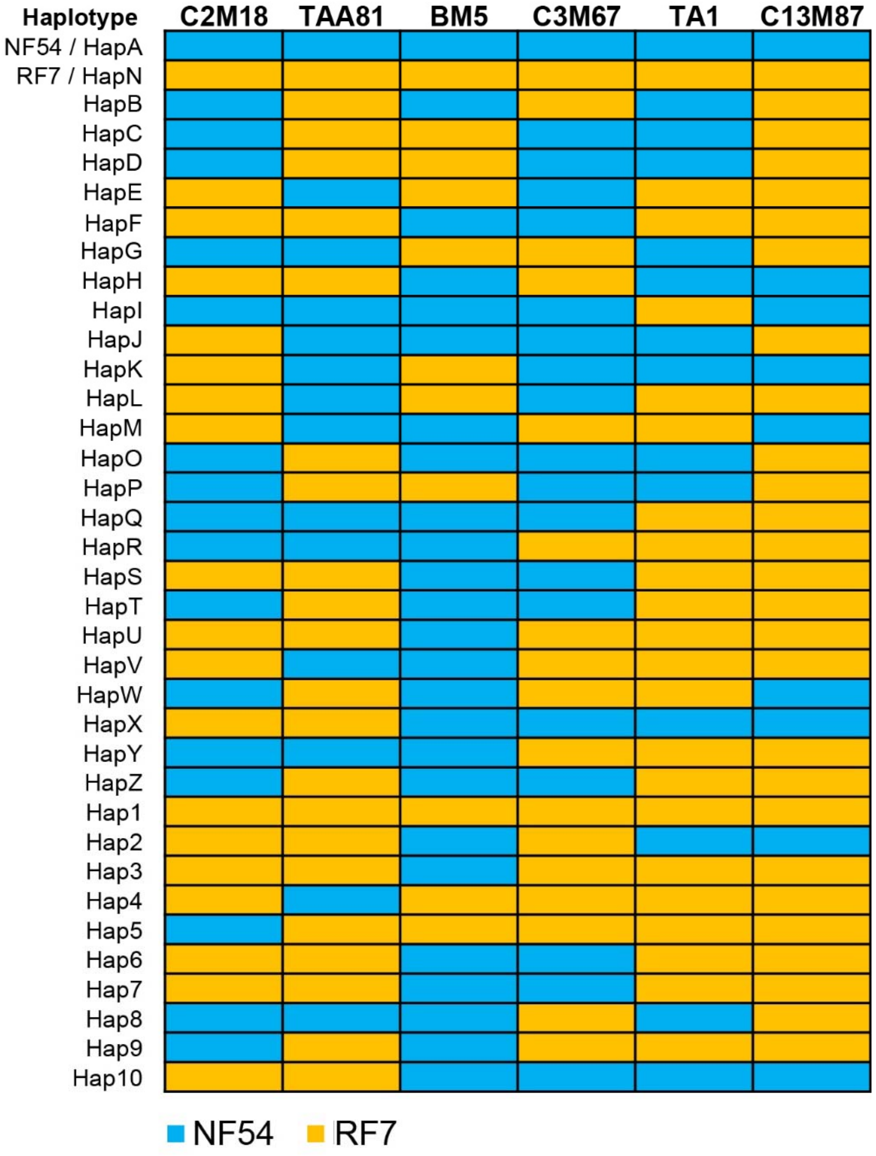


**Fig. S13. Microsatellite markers used herein to distinguish the recombinant progeny of RF7 × NF54.** The six markers, C2M18, TAA1, BM5, C3M67, TA1, and C13M87, were selected from WGS data as they differed by >7 bp in size between the parents. These markers were used subsequently for multiplexed fragment analyses to differentiate the recombinant progeny clones and to validate clonality pre- and post-drug assays.

**Legends for Tables S1 to S13.**

**Table S1.** List of first-line antimalarials and preclinical compounds tested on the genetic cross parents, RF7 × NF54, and statistical results. Listed are the drug inhibition concentration ($IC_{50}$) values for 50% survival, ring-stage survival % (RSA), piperaquine survival % (PSA), and area under the under the curve (AUC) values. Student's t-tests were conducted with corrections for multiple testing using the Holm-Sidak method for each compound. Drugs studied herein are shown in bold.

**Table S2.** *Plasmodium falciparum* oocyst, sporozoite, asexual counts in mosquito and mouse infections, and number of recombinant progeny obtained for the RF7 × NF54 genetic cross.

**Table S3.** Whole-genome sequencing metrics for the two genetic cross parents, RF7 and NF54, and the 49 progeny clones that were phenotyped and used in QTL mapping, and grouped by haplotype.

**Table S4.** The physical position, % missing or heterozygous and parental allele frequency among the 34 recombinant progeny, and segregation distortion score for each SNP marker used in the QTL analysis.

**Table S5.** List of QTL segments identified by bulk segregant analysis that were enriched by specific drugs in drug-to-drug pairwise comparisons.

**Table S6.** Genetic map of RF7 × NF54 for the core genome.

**Table S7.** Genotypes of major drug resistance markers, *pm2/3* copy number and KEL1/PLA1/PfPailin co-lineage for the RF7 × NF54 parental lines, progeny and genetically-edited clones.

**Table S8.** Drug response phenotypes of the genetic cross parents, RF7 and NF54, progeny clones, and genetically-modified lines to artemisinin, piperaquine, lumefantrine, mefloquine, MMV665939 and MMV675939.

**Table S9.** Significant QTLs associated with dihydroartemisinin (DHA) resistance and gene information.

**Table S10.** Significant QTLs associated with piperaquine (PPQ) resistance and gene information.

**Table S11.** Significant QTLs associated with lumefantrine (LMF) and mefloquine (MFQ) differential susceptibility and gene information.

**Table S12.** Significant QTLs associated with MMV665939 (MMV66) and MMV675939 (MMV67) resistance and gene information.

**Table S13.** List of oligonucleotides for gene-editing and real-time qPCR used in this study.
